# *De novo* design of highly selective miniprotein inhibitors of integrins αvβ6 and αvβ8

**DOI:** 10.1101/2023.06.12.544624

**Authors:** Anindya Roy, Lei Shi, Ashley Chang, Xianchi Dong, Andres Fernandez, John C. Kraft, Jing Li, Viet Q. Le, Rebecca Viazzo Winegar, Gerald Maxwell Cherf, Dean Slocum, P. Daniel Poulson, Garrett E. Casper, Mary L. Vallecillo-Zúniga, Jonard Corpuz Valdoz, Marcos C. Miranda, Hua Bai, Yakov Kipnis, Audrey Olshefsky, Tanu Priya, Lauren Carter, Rashmi Ravichandran, Cameron M. Chow, Max R. Johnson, Suna Cheng, McKaela Smith, Catherine Overed-Sayer, Donna K. Finch, David Lowe, Asim K. Bera, Gustavo Matute-Bello, Timothy P Birkland, Frank DiMaio, Ganesh Raghu, Jennifer R. Cochran, Lance J. Stewart, Melody G. Campbell, Pam M. Van Ry, Timothy Springer, David Baker

**Affiliations:** Department of Biochemistry and Institute for Protein Design, University of Washington, Seattle, WA 98195, USA; Encodia Inc, 5785 Oberlin Drive, San Diego, CA 92121; Department of Chemistry and Biochemistry, Brigham Young University, Provo, UT 84602, USA; Program in Cellular and Molecular Medicine, Children’s Hospital Boston, and Departments of Biological Chemistry and Molecular Pharmacology and of Medicine, Harvard Medical School, Boston, United States; State Key Laboratory of Pharmaceutical Biotechnology, School of Life Sciences, Nanjing University, Nanjing, China; Engineering Research Center of Protein and Peptide Medicine,Ministry of Education; Division of Basic Sciences, Fred Hutchinson Cancer Center, Seattle, WA 98109, USA; Department of Bioengineering, Stanford University, Stanford CA 94305; Denali Therapeutics, South San Francisco, CA, USA; Department of Materials Science and Engineering, University of Washington, Seattle, WA 98195, USA; Department of Pharmacology, Northwestern University Feinberg School of Medicine; Chicago, IL 60611, USA; Research and Early Development, Respiratory and Immunology, BioPharmaceuticals R&D, AstraZeneca, Cambridge, United Kingdom; Alchemab Therapeutics Ltd, Cambridge, United Kingdom; Evox Therapeutics Limited, Oxford Science Park, Medawar Centre, East Building, Robert Robinson Avenue, Oxford, OX4 4HG; Center for Lung Biology, Division of Pulmonary, Critical Care and Sleep Medicine, University of Washington; Division of Pulmonary, Critical Care and Sleep Medicine, Department of Medicine, University of Washington, Seattle, Washington; Dept of Medicine, University of Washington, Seattle WA, USA; Howard Hughes Medical Institute, University of Washington, Seattle, WA 98195, USA; Department of Medicine Solna, Division of Immunology and Allergy, Karolinska Institutet and Karolinska University Hospital, Stockholm, Sweden; Department of Bioengineering, University of Washington, Seattle, WA 98195, USA; Bioscience COPD/IPF, Research and Early Development, Respiratory and Immunology, BioPharmaceuticals R&D, AstraZeneca, Cambridge, UK

## Abstract

The RGD (Arg-Gly-Asp)-binding integrins αvβ6 and αvβ8 are clinically validated cancer and fibrosis targets of considerable therapeutic importance. Compounds that can discriminate between the two closely related integrin proteins and other RGD integrins, stabilize specific conformational states, and have sufficient stability enabling tissue restricted administration could have considerable therapeutic utility. Existing small molecules and antibody inhibitors do not have all of these properties, and hence there is a need for new approaches. Here we describe a method for computationally designing hyperstable RGD-containing miniproteins that are highly selective for a single RGD integrin heterodimer and conformational state, and use this strategy to design inhibitors of αvβ6 and αvβ8 with high selectivity. The αvβ6 and αvβ8 inhibitors have picomolar affinities for their targets, and >1000-fold selectivity over other RGD integrins. CryoEM structures are within 0.6-0.7Å root-mean-square deviation (RMSD) to the computational design models; the designed αvβ6 inhibitor and native ligand stabilize the open conformation in contrast to the therapeutic anti-αvβ6 antibody BG00011 that stabilizes the bent-closed conformation and caused on-target toxicity in patients with lung fibrosis, and the αvβ8 inhibitor maintains the constitutively fixed extended-closed αvβ8 conformation. In a mouse model of bleomycin-induced lung fibrosis, the αvβ6 inhibitor potently reduced fibrotic burden and improved overall lung mechanics when delivered via oropharyngeal administration mimicking inhalation, demonstrating the therapeutic potential of *de novo* designed integrin binding proteins with high selectivity.

## Introduction

The highly homologous integrins αvβ6 and αvβ8 bind to latent transforming growth factor-β1 and β3 (L-TGF-β1 and L-TGF-β3) leading to release of active TGF-β1 and -β3.^1–3^ Upregulation of αvβ6- and/or αvβ8-mediated TGF-β activation is a driver of multiple diseases, including idiopathic pulmonary fibrosis (IPF)^4–6^, primary sclerosing cholangitis (PSC),^7^ and several solid tumors^8–10^, but deconvoluting the contribution of αvβ6 and αvβ8 to the etiology of these diseases has been challenging due to limitations in current interventions. Selective antibodies targeting RGD integrins have been generated by immunizing mice,^11–13^ but this approach lacks precise control over the target binding site on the integrin. Control over the target site is important because differential modulation of αvβ6 integrin conformations (bent-closed, extended-closed, and extended-open) by orthosteric and allosteric inhibitors has dramatically different outcomes on receptor internalization^11,14,15^ and led to different safety signals in preclinical and clinical studies.^16^ For example, the mAb BG00011 and small molecule MORF-720 both target the non-internalized, bent-closed αvβ6 conformation^11,15^ and have on-target/αvβ6-mediated toxicity,^17–19^ while the small molecules PLN-74809^20^ and GSK3008348^21^ stabilize the extended-open αvβ6 conformation that induces αvβ6 internalization, and have not shown any drug-related serious adverse events in clinical trials.^22,23^ Since eight integrin heterodimers, including αvβ6 and αvβ8, share the conserved RGD binding sequence, it has not been possible to generate selective RGD-mimetic small molecules for individual integrins, making it challenging to dissect the role a single integrin plays in a particular disease.^24^ Therefore, there is a need for a new integrin therapeutic modality with (i) high selectivity for a single RGD integrin heterodimer, (ii) atomic-level control over the precise location of the target binding site and the protein-protein interaction interfaces to control the evoked integrin conformation, (iii) hyperstability to enable tissue restricted administration (inhaled and oral), and (iv) a smaller hydrodynamic size than IgG antibodies to enable better tissue penetration.

### Computational design strategy

We set out to overcome the limitations of integrin-targeted small molecules and antibodies by developing a computational approach that generates small (<75 amino acids) hyperstable *de novo* integrin binding proteins that have high integrin selectivity and specific receptor binding interfaces optimal for treating disease. Integrin αvβ6 and αvβ8 both bind to a RGDLXX(L/I) motif in the pro-domains of L-TGF-β1 and β3 with low nM affinity (Fig. 1a).^1,25^ As in other structures of RGD-containing peptides bound to integrins, the arginine and aspartate side chains make multiple hydrogen bond and salt-bridge interactions to residues at the interface between the integrin alpha and beta subunits (Fig. 1b, 1c). For both αvβ6 and αvβ8, C-terminal to the RGD, the peptide adopts an alpha-helix-like turn with two leucines (or Ile for β8) fitting into a hydrophobic pocket formed by a β6/β8 subunit specificity determining loop 2 (SDL2, Figure 1d, 1e).^1,2,25^ In the unliganded state, SDL2 of αvβ6 is ordered with multiple backbone hydrogen bonds (PDB ID 4UM8), whereas SDL2 of unliganded αvβ8 is mainly flexible.^1,2,25^ To engineer selectivity, we focused on two main areas on the β subunit that differ between the two targets: the region that contacts the LXX(L/I) motif in the L-TGF-β3 peptide (Fig. 1d, 1e) and a charge reversal on the β subunit (Fig. 1f). There are several key differences in the hydrophobic packing pattern of LXX(L/I) motif and SDL2 of β6 compared to β8 (Fig. 1d, 1e, 1k). Y185 from SDL2 of αvβ6 packs optimally with Leu (LXX(**L**/I), L247) of the L-TGF-β3 peptide (PDB ID 4UM9, Fig. 1d), while the equivalent position on the SDL2 of αvβ8 (L174) packs much less tightly with Ile (LXX(L/**I)**, I221) of L-TGF-β1 (PDB ID 6OM2, Fig. 1e). There is also a key charge reversal on the β subunit; β8 contains K304 whereas the equivalent position on β6 is E316 (Fig. 1f). We hypothesized that minibinders interacting with the Y185/L174 and E316/K304 regions of αvβ6 and αvβ8, respectively, might be able to achieve selectivity between the two proteins.

**Figure 1.**
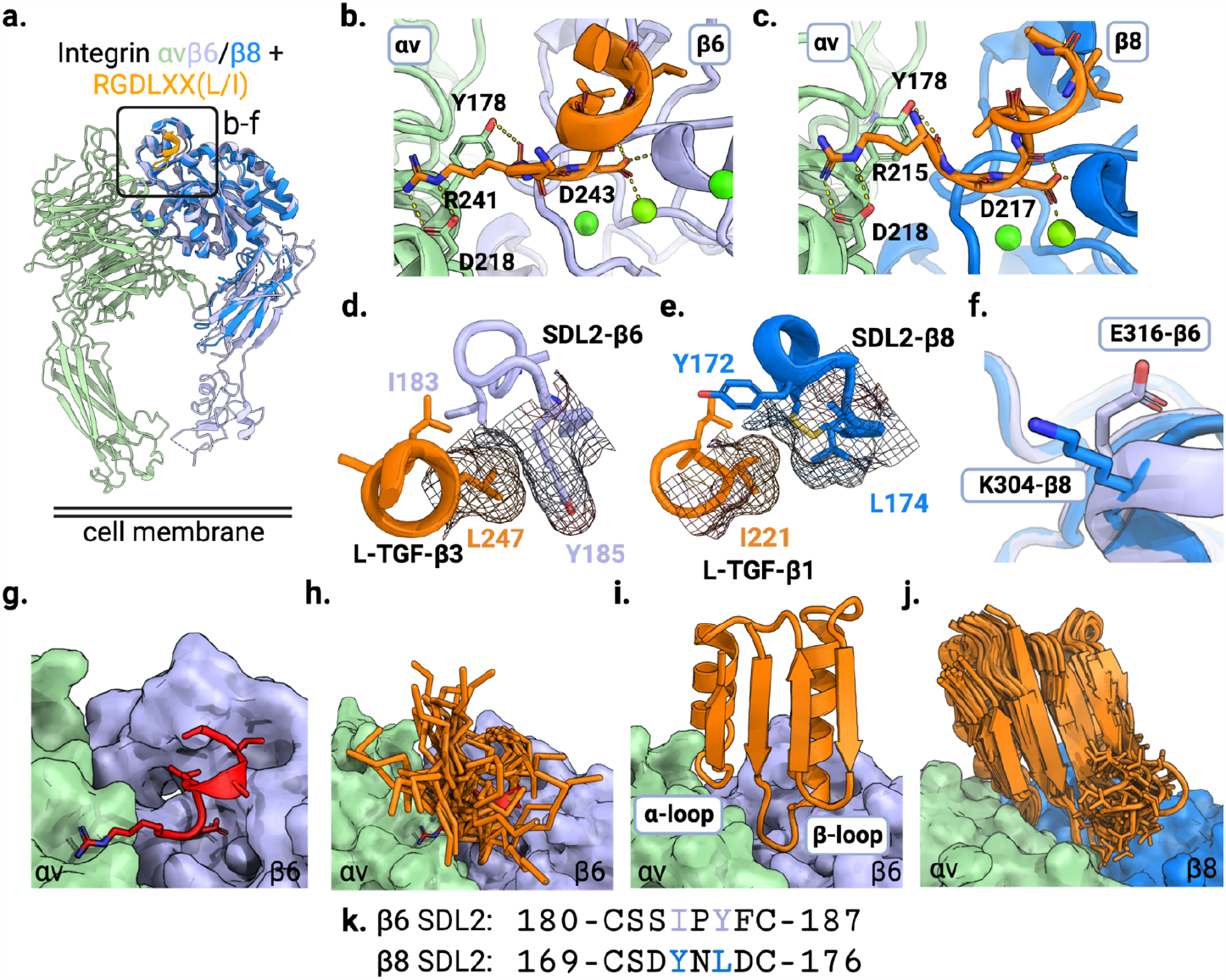
Computational design of αvβ6 and αvβ8 selective minibinders. **a)** Crystal structure of β8 (PDB ID 6OM2) overlaid on the structure of αvβ6 integrin in complex with the L-TGF-β3 peptide RGDLXX(L/I) (PDB ID 4UM9). Inset highlights the zoomed-in regions shown in panels b-f. The αv subunit is shown in green, β6 is in lavender, β8 is in blue, and the RGDLXX(L/I) peptide is in orange. **b**,**c)** Shared polar interactions between the RGD motif and (b) αvβ6 and (c) αvβ8 integrin (L-TGF-β3/αvβ6 complex PDB ID 4UM9, L-TGF-β1/αvβ8 PDB ID 6OM2). **d)** Hydrophobic packing of LXX(L/I) motif of L-TGF-β3 peptide with SDL2 of αvβ6 (PDB ID 4UM9). Leu247 packs optimally against Y185 from SDL2 of β6. **e)** Hydrophobic packing of LXX(L/I) of L-TGF-β1 peptide with SDL2 of αvβ8 (PDB ID 6OM2). I221 of L-TGF-β1 peptide packs less tightly against L174 on SDL2 of β8 compared to the homologous interactions in panel d. **f)** Charge reversal on β subunit: β8 contains K304 whereas the equivalent position on β6 is E316. **g)** Surface structure of the αvβ6 integrin in complex with L-TGF-β3 peptide (red cartoon representation, PDB ID 4UM9). **h)** Low RMSD matches to the L-TGF-β3 peptide bound to αvβ6 were harvested from the PDB database (orange stick representations). **i)** Non-clashing fragments with αvβ6 were then incorporated in the α/β ferredoxin folds (orange ribbon representation) using Rosetta. **j)** Loop extension strategy to design an αvβ8 selective minibinder: to make more extensive contacts to the β8 subunit the β-loop was resampled by one residue insertion (blue surface representation for β8 subunit, PDB ID 6OM2). In addition to the loop extension, the LXX(L/I) motif was allowed to be redesigned using Rosetta. **k)** Partial sequence alignment of SDL2 of the β6/β8 subunits is shown highlighting two key positions packing against the LXX(L/I) motif of the L-TGF-β ligand (I183 and Y185 in SLD2-β6, and Y172 and L174 in SDL2-β8).

To implement this design strategy, we sought to generate small proteins that incorporate the central RGD affinity loop, make favorable contacts with both α and β subunits, and interact closely with the two structurally diverging regions described above. We started from the crystal structure of human αvβ6 in complex with an RGD-containing L-TGF-β3 peptide (PDB ID 4UM9),^1^ and screened the PDB database *in silico* for topologies and structure segments capable of hosting the 8 residue extended turn conformation of the peptide (RGDLGALA, Fig. 1g). Low RMSD matches to the peptide backbone conformation, along with the five flanking residues on both the N- and C-termini, were superimposed on the bound peptide conformation in the complex structure, and those making backbone level clashes with the integrin were discarded (Fig. 1h). We found small α/β ferredoxin folds (Fig. 1i) were able to scaffold the RGDLXX(L/I) binding loop without clashing with the integrin while making additional contacts with both α and β subunit (α- and β-loop respectively, Fig. 1i). Structures were assembled from fragments following rules for constructing ideal proteins,^26^ sampling different alpha helix, beta sheet, and loop lengths, while constraining torsion angles in the region corresponding to the RGD peptide to those observed in the co-crystal structure using Rosetta (Fig. 1g). Following two rounds of design and optimization (see supplementary info for details), two high affinity variants were selected for further characterization: B6B8_BP (av6_3_E13T) and B6_BP (av6_3_A39KG64R) (Extended Data Figs. 5, 6). Any mutation to the LXX(L/I) motif was depleted during affinity maturation, confirming the importance of this motif for selectivity towards αvβ6 (Extended Data Fig. 5b). Additionally, the A39K mutation makes a salt bridge interaction with E316 of the β6 subunit (Extended Data Fig. 5f) where there is a charge reversal for the β8 subunit (K304) (Fig. 1f).

To achieve selectivity for the β8 subunit, we redesigned the β-loop to take advantage of the K304 charge reversal on the β subunit (Fig. 1f). We generated 200 models with different lengths and conformations of the β-loop using RosettaRemodel^27^ and the resulting models were superimposed on the L-TGF-β1 / αvβ8 complex structure (PDB ID 6OM2) by superposition on the RGD peptide (Fig. 1j). As packing of the L-TGF-β1 LXX(L/I) motif with SDL2 of αvβ8 integrin is suboptimal (Fig. 1e), we hypothesized that a minibinder mimicking this interaction would be able to accommodate bulkier residues at these positions, giving additional selectivity. Both the β-loop and LXX(L/I) motif were redesigned using Rosetta and a total of 9 designs with lowest predicted binding energy were selected following structure prediction using AlphaFold.^28^ Four out of 9 designs showed preferential binding to αvβ8 integrin with B8_BP_dslf showing the highest affinity and selectivity towards αvβ8 (Extended Data Fig. 7). Mutations from the B8_BP_dslf sequence (MAVY) to LATI (avb8_#12, which corresponds to the L-TGF-β1 peptide sequence) completely abrogated selectivity towards αvβ8 on the yeast surface displayed design (Extended Data Fig. 7b), indicating these residues are critical for selectivity against αvβ8 vs. αvβ6. B8_BP_dslf is a monomeric and hyperstable protein when expressed in *E. coli* and binds to human αvβ8 with 1.9 nM affinity, with no appreciable binding to human αvβ6 up to 1 μM (Extended Data Fig. 6b). In agreement with yeast surface display data, purified avb8_12 with the reversion mutations loses selectivity towards αvβ8 and binds to αvβ6 with a Kd of 1.13 nM (Extended Data Table 1, Extended Data Fig. 6), confirming the importance of the LXX(L/I) motif for selectivity. B8_BP_dslf has a MAVY motif that packs against SDL2 of αvβ8. We systematically varied each position within the LATI motif to determine which residue plays a critical role in determining selectivity (Fig. 2e). We found a single mutation containing the LAT**Y** (B8_BP-LATY) motif binds to αvβ8 with an affinity of 500 pM with no appreciable binding to αvβ6 at 500 nM concentration (Fig. 2e, Extended Data Fig. 6b).

**Figure 2.**
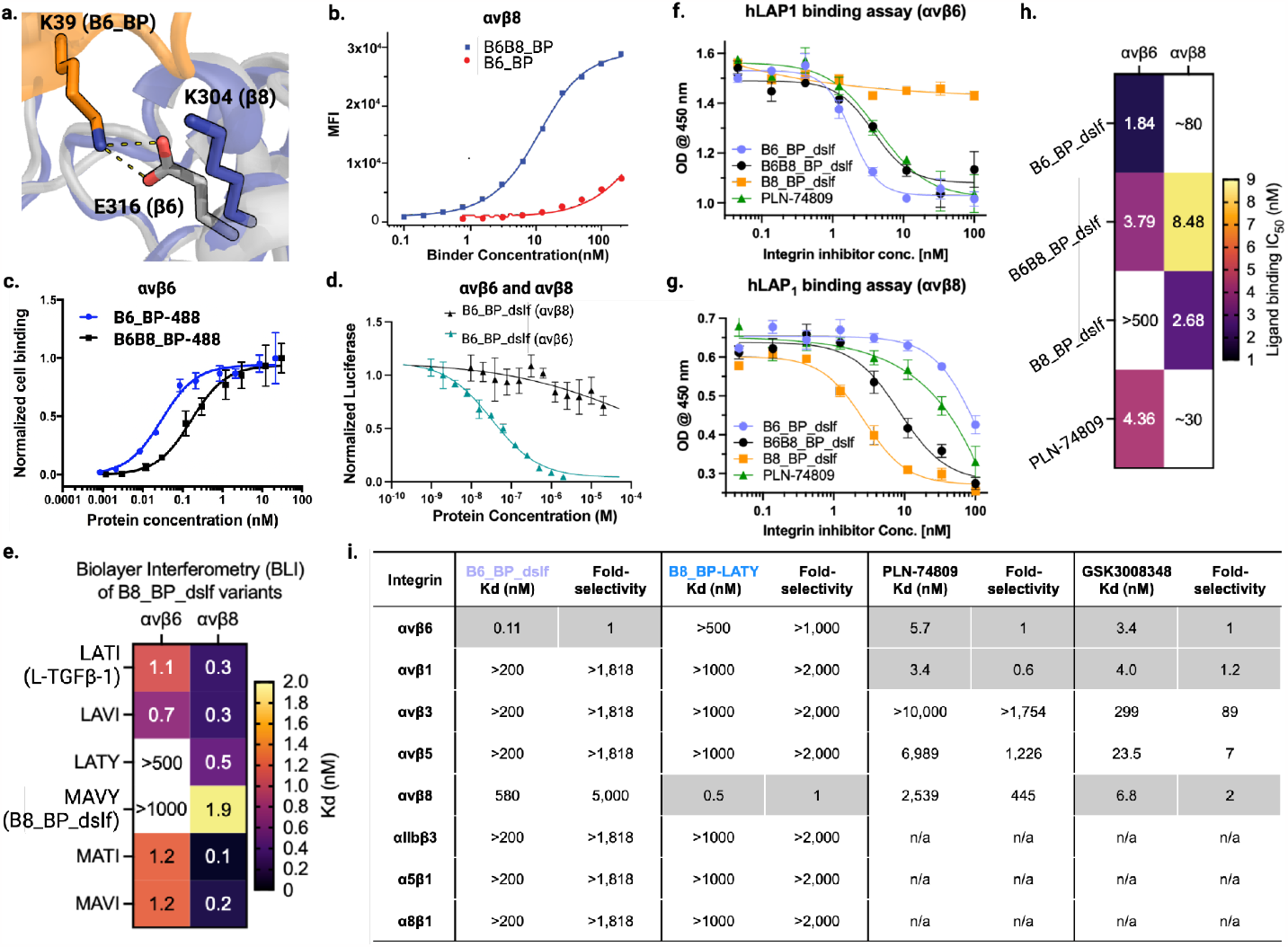
Selectivity of designed binders for αvβ6 and αvβ8. **a)** The A39K mutation confers selectivity towards αvβ6 compared to αvβ8 where there is a charge reversal (Glu316 for β6 shown as a gray stick, Lys304 for β8 shown as a blue stick). **b)** Cell surface titration of B6B8_BP and B6_BP against K562 cells stably transfected with αvβ8. B6B8_BP lacking the A39K mutation binds to αvβ8 with a Kd of ∼7.3 nM whereas B6_BP containing the A39K mutation binds to αvβ8 >500 nM. **c)** Cell surface titration of AlexaFluor-488-labelled B6B8_BP and B6_BP using αvβ6 (+) human epidermoid A431 carcinoma cells. B6_BP binds to A431 cells with higher potency than B6B8_BP (30 pM vs 167 pM). **d)** B6_BP_dslf selectively inhibits αvβ6-mediated TGF-β1 activation. αvβ6 and αvβ8 transfectants were co-incubated with CAGA-reporter cells and GARP/TGF-β1 transfectants and inhibitors. B6_BP_dslf inhibits TGF-β activation with an IC50 of 32.8 nM. **e)** Binding affinities (Kd) of B8_BP_dslf (MAVY) point mutants to integrins αvβ6 and αvβ8, determined by BLI. The LATI motif is in native L-TGF-β1. **f, g)** Competitive inhibition of h-LAP1 binding to (f) αvβ6 and (g) αvβ8 by designed inhibitors and control small molecule PLN-74809.^20^ **h)** Heatmap of IC50 values for h-LAP1 binding assays in f and g. **i)** Binding affinities (Kd) and fold-selectivity values of B6_BP_dslf and B8_BP-LATY to all eight RGD integrins compared to small molecules PLN-74809^20^ and GSK3008348.^21^ Binding data for PLN-74809 and GSK3008348 are taken from Decaris et al. 2021.^20^ Rows shaded in gray indicate the RGD integrin(s) for which each molecule is selective (i.e., B6_BP_dslf and B8_BP-LATY are both mono-selective whereas PLN-74809 and GSK3008348 are dual- and tri-selective, respectively). n/a, not available.

### Selectivity profiles of αvβ6 and αvβ8 binders towards other RGD binding integrins

B6_BP and B8_BP_dslf are highly selective to αvβ6 and αvβ8, respectively. We investigated the selectivity of the designed binders against seven other RGD-binding integrins. B6B8_BP and B6_BP do not cross-react with RGD-binding integrins αvβ1, αvβ3, αvβ5, α5β1, α8β1, and αiibβ3 at concentrations up to 200 nM in cell surface binding experiments using K562 cells stably transfected with different RGD-binding integrins, corresponding to >1000-fold selectivity (Extended Data Fig. 8a, 8c). In B6_BP, the β-loop is positioned to confer selectivity between the two integrins, where residue K39 faces E316 on the β6 subunit and K304 on β8 (Fig. 2a). As intended, B6_BP is more selective for αvβ6 than B6B8_BP. B6_BP binds αvβ6 with a higher affinity than αvβ8 on the surface of K562 cells, with a *K*_*d*_ of 0.11 (± .09) and 580 (± 40) nM, respectively (Fig. 2b, Extended Data Fig. 8b). B6B8_BP, which has an alanine at this position (A39), is less selective for αvβ6, and binds to αvβ6 and αvβ8 with a *K*_*d*_ of 1.7 (± 0.2) and 7.3 (± 1.2) nM, respectively (Fig. 2b, Extended Data Fig. 8b). We generated fluorescently labeled B6B8_BP and B6_BP by conjugating AlexaFluor-488 to an engineered C-terminal cysteine via maleimide chemistry. The fluorescently labeled proteins were titrated against αvβ6 (+) human epidermoid carcinoma A431 cells. B6B8_BP and B6_BP bind to A431 cells with *K*_*d*_ values of 167 (± .028) pM and 30 (± .004) pM, respectively (Fig. 2c).

As B6_BP starts unfolding close to ∼90°C (data not shown), we sought to further increase the stability of the engineered inhibitors.^29,30^ Four variants with additional disulfide bonds stapling the N and C terminus had considerably increased thermostability (Extended Data Fig. 9), and bound to αvβ6 with subnanomolar affinity; mutation of the RGD to KGE abrogated binding to αvβ6 confirming that the RGD loop is necessary for binding (Extended Data Fig. 6a). We further characterized one variant, B6_BP_dslf, and found that it selectively inhibited αvβ6-mediated TGF-β activation (IC50 32.8 ± 3.4 nM) using CAGA reporter cells^31^ and GARP/TGF-β1 transfectants, and had marginal effect on αvβ8-mediated TGF-β activation in the tested concentration range, confirming the selectivity towards αvβ6 (Fig. 2d, Extended Data Fig. 10d, 10e). αvβ8-selective B8_BP_dslf does not bind to any other RGD binding integrins as confirmed by BLI (Extended Data Fig. 6b).

We compared the potency and selectivity of our designed αvβ6 and αvβ8 minibinders to the small-molecule dual αvβ6/αvβ1 inhibitor (PLN-74809) currently in clinical trials as an oral IPF therapy,^20^ by assessing their ability to outcompete binding of hLAP_1_, the endogenous ligand of αvβ6 and αvβ8. For αvβ6 integrin, B6_BP_dslf had the lowest IC_50_ (1.84 nM), followed by B6B8_BP_dslf (3.79 nM), PLN-74809 (4.36 nM), and no significant binding of B8_BP_dslf to αvβ6 integrin was detected (Fig. 2f, 2h). For αvβ8, B8_BP_dslf outcompeted hLAP_1_ with the lowest IC_50_ (2.68 nM), followed in order of potency by B6B8_BP_dslf (8.48 nM), PLN-74809, and B6_BP_dslf (Fig. 2g, 2h). Taken together, these data confirm that B6_BP_dslf and B8_BP_dslf have exquisite selectivity and affinity for their respective integrin targets, with considerably greater RGD integrin selectivity than the small molecules PLN-74809 and GSK3008348 (Fig. 2i).

### Negative stain EM reveals B6_BP_dslf stabilizes the αvβ6 open headpiece conformation

Integrin αvβ6 adopts the well-characterized range of integrin conformations including bent-closed, extended-closed, and extended-open, which have been linked to activation and binding site accessibility.^3^ However, αvβ8 has not been observed in this range of conformations and instead has been shown to bind and activate L-TGF-β while exclusively occupying the extended-closed conformation.^2^ To confirm the conformational effects of minibinder binding on global αvβ6 conformations, we used single-particle negative stain electron microscopy to image complexes of minibinder binding on glycosylated soluble construct of the αvβ6 headpiece. All minibinder-integrin complexes were formed in a buffer containing excess Mn^2+^ ions to push the conformational equilibrium towards extended-open and to ensure the availability of the MIDAS cation, which is known to be crucial for ligand binding. As expected, the 2D class averages showed that both B6_BP_dslf and B8_BP_dslf bind at the canonical ligand binding site at the alpha / beta subunit cleft in αvβ6 and αvβ8, respectively (Fig. 3a). Furthermore, B6_BP_dslf induces αvβ6 headpiece opening whereas B8_BP_dslf does not have an effect on the global conformation of αvβ8 and the headpiece remains closed (Fig. 3a, Extended Data Fig. 18).

**Figure 3.**
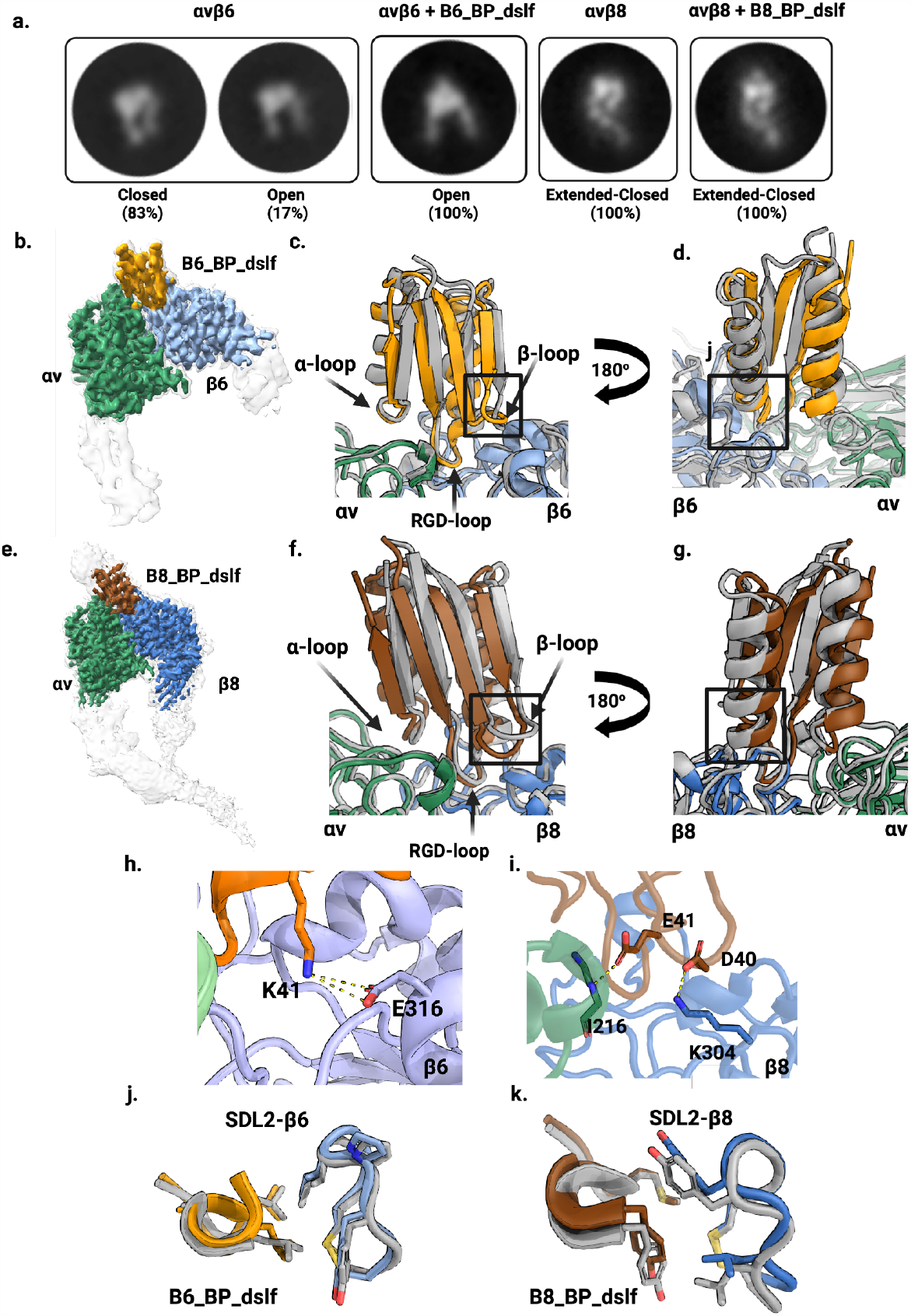
Structural characterization. **a)** Representative 2D class averages of integrin with and without minibinder. For αvβ6, both closed and open headpiece conformations are present in the unbound state, but in the presence of the minibinder the open conformation is dominant. For αvβ8, no open headpieces were observed, with or without minibinder. **b)** CryoEM density map of αvβ6 bound to minibinder B6_BP_dslf. B6_BP_dslf (goldenrod) binds the integrin ligand binding cleft between the αv (green) and β6 (light blue) subunits and induces or stabilizes the open conformation. The sharpened, locally refined cryoEM map is shown in color, superimposed with the unsharpened map showing all domains of the αvβ6 headpiece in semi-transparent white. **c**,**d)** Overlay of the designed αvβ6 + B6_BP_dslf model (gray) and the experimentally determined cryoEM model (colors). Although the overall angle of the minibinder is shifted, the RGD loop positioning is as predicted. Insets in c) and d) are magnified in panels h) and i), respectively. **e)** CryoEM density map of αvβ8 bound to minibinder B8_BP_dslf. Similar to B6_BP_dslf, B8_BP_dslf (brown) binds the integrin ligand binding cleft between the αv (green) and β8 (blue) subunits, however the conformation of αvβ8 remains in the closed headpiece conformation. The sharpened, locally refined cryoEM map is shown in color, superimposed with the unsharpened map showing all domains in the αvβ8 ectodomain construct in semi-transparent white. **f**,**g)** An overlay of the designed αvβ8 + B8_BP_dslf model (gray) and the experimentally determined model. Although the overall angle of the minibinder is shifted, the RGD loop positioning is as predicted. Insets in f) and g) are magnified in panels i) and k), respectively. **h**,**i)** Key designed interactions between β-loop and β6/β8 subunit are observed in the cryoEM structure: K41 from B6_BP_dslf forms a salt bridge with E316 from the β6 subunit (panel h). E41 from β-loop makes backbone level hydrogen bond with I216 from β8 subunit and D40 makes salt bridge interaction with K304 from β8 subunit (panel i). **j)** Experimental vs designed (gray) packing pattern of the LXXL motif and SDL2 of αvβ6. **k)** Experimental vs designed (gray) packing pattern of the MAVY motif and SDL2 of αvβ8.

### cryoEM structure characterization

To investigate the accuracy of our designed structures, we used single-particle cryoelectron microscopy (cryoEM) to determine structures of a stable complex of human αvβ6 ectodomain bound to B6_BP_dslf and human αvβ8 ectodomain bound to B8_BP_dslf. Using focused refinement, the nominal overall resolution is 3.4Å for the αvβ6 - B6_BP_dslf complex and 2.9Å for the αvβ8 - B8_BP_dslf complex, although the resolution varies considerably due to the intrinsic flexibility of both integrins (Supplementary Table 7, Extended Data Fig. 19).^2^ Both integrin-minibinder complexes have extensive binding interfaces (Extended Data Table 3). The structure of the αvβ6 - B6_BP_dslf complex identifies several glycosylation sites that had not previously been observed structurally. Despite extensive 3D classification of the αvβ6 - B6_BP_dslf complex (Extended Data Fig. 19), we did not observe any subclasses of the integrin-minibinder complex in a closed headpiece conformation. As expected based on the negative stain class averages, the αvβ8 - B8_BP_dslf was found to be exclusively in the extended-closed conformation.

The secondary structural elements of the designed B6_BP_dslf minibinder model are in close agreement with the cryoEM map (complex RMSD 0.6 Å vs design, Fig. 3b, 3c, 3d) and the three minibinder loops make contact with the alpha, the beta, or both subunits of integrin, although some interactions vary slightly from the initial design (Extended Data Table 3). B6_BP_dslf was designed using a closed integrin headpiece (PDB ID 4UM9),^1^ however, our cryoEM map revealed that, when bound to minibinder, the βI domain of the β6 subunit rearranges to the same open conformation as when bound to ligand (Fig. 3b). In the cryoEM map, the overall orientation of the minibinder is shifted relative to initial design, but the RGD loop positioning is as predicted (Fig. 3c, 3d). As expected, the RGD loop spans the subunit binding interface with RGD-Arg10 forming a hydrogen bond with D218 of the αv-subunit and RGD-Asp12 of the minibinder in a position to coordinate with the MIDAS cation. The engineered positively charged, affinity-enhancing point mutation in B6_BP_dslf, A41K (A39K in B6_BP), interacts with negatively charged E316 of the β2 subunit to form a salt bridge (Fig. 3h). Although the second charge reversal mutation (G64R in B6_BP, G66R in B6_BP_dslf) does not form the anticipated salt bridge with D148 in our structure, we note that this binding surface has a strong negative charge and speculate that this stabilizes the positively charged Arg. As predicted, we observe hydrophobic packing of LXX(**L**/I) L16 with Y185 of the β6 subunit (Fig. 3j). Of the 13 interacting pairs of residues present in the cryoEM model, 11 are present in the computational design model, including all three salt bridges. The two unanticipated interactions were backbone interactions with integrin: minibinder R10 and αv A21 and minibinder RGD-Asp12 and β6 S127 (Extended Data Table 3).

The cryoEM model of αvβ8 - B8_BP_dslf complex is also very close to the computational design model (Fig. 3e, 3f, 3g, complex RMSD 0.7 Å). In the cryoEM model of the αvβ8 - B8_BP_dslf complex, there are 12 interacting pairs of residues between integrin and the minibinder (Extended Data Table 3); as in the design model, the α-loop interacts with αv, β-loop with β8, and the RGD loop spans the two subunits (Fig. 3f). Y172 of the β8-SDL2 loops bends inward to form a hydrophobic patch similar to the conformation in L-TGF-β-bound structures (Fig. 3k).^2^ We showed above that altering the known binding motif LXX(L/I) to LAT**Y** confers selectivity for αvβ8 (Fig. 2e) and the cryoEM structure reveals the molecular basis for this selectivity: we find that Y16 forms stabilizing interactions with A115 of the β8 subunit and interacts with the less bulky L174 in the β8-SDL2 loop (Fig. 3k). The equivalent position in the β6-SDL2 loop, Y185, is too bulky and we hypothesize that the steric clash would interfere with binding.

### *In vivo* tumor targeting using fluorescently labeled B6_BP

As B6_BP binds to A431 cells with higher affinity and is more selective than B6B8_BP for αvβ6, we selected B6_BP for further *in vivo* experiments. We prepared tumor bearing rodents by injecting 6-8 week old female athymic nude mice with A431 cells (αvβ6 (+)) and HEK 293T (αvβ6 (-)) into the left and right shoulders, respectively. When the tumors reached 5-10 mm in diameter, mice were injected via the tail vein with 1.5 nmols of AlexaFluor-680 labeled B6_BP (AF680-B6_BP). AF680-B6_BP rapidly accumulated in the αvβ6 positive tumors and reached a high tumor-to-muscle fluorescence contrast ratio within 3 hours post-injection (Fig. 4a, Extended Data Fig. 11). There was no detectable fluorescence at the αvβ6 negative HEK-293T tumors (Fig. 4a). We also performed a semiquantitative *ex vivo* biodistribution analysis of AF680-B6_BP at 6 hours post-tail vein injection. Analysis of fluorescence intensities of different tissues revealed accumulation of AF680-B6_BP to αvβ6 positive tumors and kidney (tumor-to-kidney ratio 1:1.04) with no significant off-target binding including αvβ6 negative tumors (Fig. 4b). Quantification of whole body imaging data for AF680-B6_BP (Fig. 4b) suggests glomerular filtration through the kidneys into the urine is the primary route of elimination.^32^ We also characterized the pharmacokinetics of B6_BP_dslf in the lungs and serum of healthy male C57BL/6 mice following a single dose via different routes of administration. B6_BP_dslf was rapidly cleared from the blood following IV and IP administration with a half-life of ∼10 min, and following inhaled administration, B6_BP_dslf had a half-life in the lungs of ∼1 hr (Extended Data Fig. 12).

**Figure 4:**
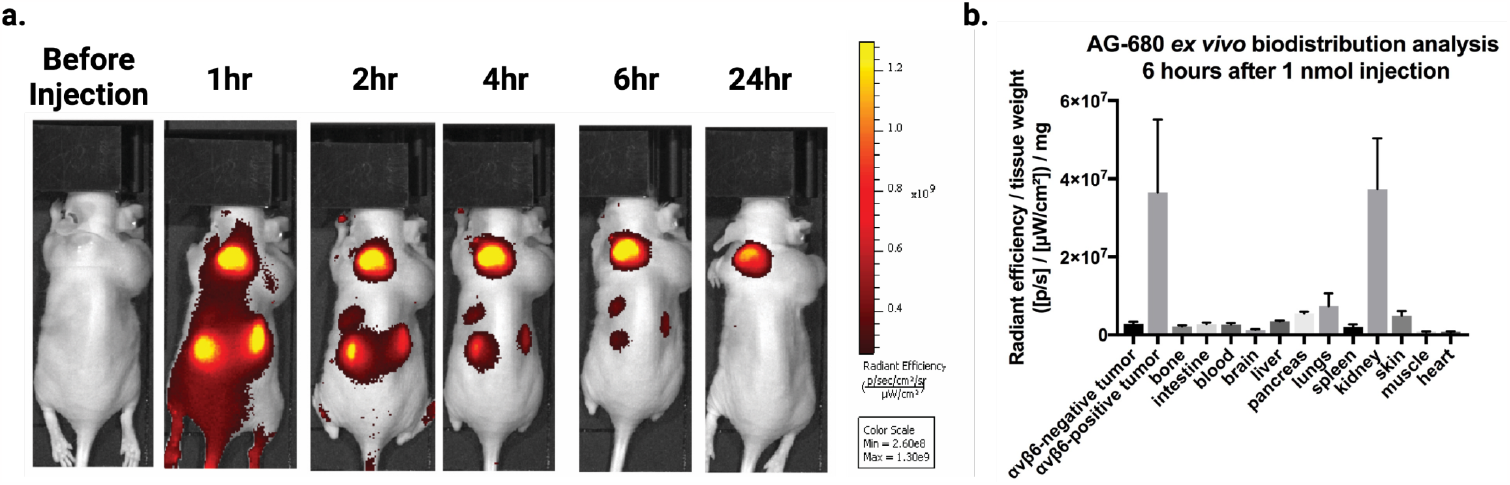
*In vivo* imaging of αvβ6 (+) A431 tumors using fluorescently-labeled B6_BP. a) Athymic nude mice were injected with αvβ6 (+) A431 cells on the left shoulder and αvβ6 (-) HEK293T cells on the right shoulder. AlexaFluor-680-labelled B6_BP (AF680-B6_BP) was injected via the tail vein to image the tumors over time as indicated (see Extended Data for additional images, n=5). b) Semiquantitative *ex vivo* biodistribution assay of AF-680-B6_BP at 6 hours post-tail vein injection. B6_BP selectively accumulates in αvβ6 (+) tumors and primarily clears via glomerular filtration in the kidneys.

### *In vivo* efficacy of B6_BP_dslf in bleomycin-induced IPF

We investigated the therapeutic efficacy of B6_BP_dslf using the bleomycin-induced pulmonary fibrosis (PF) mouse model. 12-week-old C57BL/6 male mice were administered bleomycin intratracheally at 1 U/kg body weight to induce fibrosis in a “mild” and “severe” manner (see Methods for details) to mirror the progressive stages of PF. Therapeutic dosing was confirmed by first showing fibrotic development at day 7 in our longitudinal High-Resolution micro-Computed Tomography (HR-μCT) renders (Extended Data Fig. 13c, 15a) and therefore the data described here verifies treatment began after the development of fibrosis. As proof-of-principle, we administered B6_BP_dslf intraperitoneally at 100 μg/kg in the mild bleomycin model and at 1 mg/kg in the severe bleomycin model starting on day 7 after bleomycin instillation and found significant improvement in lung health and function through HR-μCT and lung function measurements (Extended Data Figs. 13-15). For both doses, B6_BP_dslf halted fibrotic progression as evident by the reduced Ashcroft scores and improved respiratory mechanics such as static compliance and forced vital capacity (FVC) (Extended Data Figs. 13-15).

A clinical trial involving an antibody targeting αvβ6 (BG00011) from Biogen has been discontinued because it exacerbated disease at higher doses among other serious adverse effects (SAE) including mortality.^17^ The SAEs were attributed to increased alveolar inflammation, increased MMP12 generation, and emphysema due to the long half-life of BG00011.^33,34^ An inhaled, tissue-restricted therapy delivered directly to the site of action of fibrosis in the lung could result in a considerably safer and more effective option than systemic inhibition of αvβ6-mediated TGF-β activation; therefore, we pursued a respiratory system delivery. To mimic inhalation^35^, mice were administered B6_BP_dslf via oropharyngeal administration (OA, 43.6 and 185.2 μg/kg) every-other-day starting at day 7 post-bleomycin instillation (using the severe bleomycin application method), ending on day 19, for a total of 7 treatments. Non-treated mice were given neither bleomycin nor B6_BP_dslf and serve as a healthy lung control to identify B6_BP_dslf efficacy. Three-dimensional renders and representative slices of the HR-μCT scans show increased healthy lung tissue available for segmentation in the 185.2 μg/kg B6_BP_dslf group compared to bleomycin controls (Fig. 5a, upper and middle panels). Quantification of HR-μCT scans shows a significant rescue of healthy lung volume following 185.2 μg/kg OA treatment and a shift away from fibrotic intensities (Extended Data Figs. 16a,b). Ashcroft scoring of Masson-trichrome stained lung sections by a blinded veterinary pathologist was significantly reduced in the 185.2 μg/kg B6_BP-disulf treatment group compared to the BLM and 43.62 μg/kg B6_BP-disulf treatment group (Fig. 5b). Western blot analysis of the whole lung homogenate lysates shows a reduction in TGF-β mechanistic biomarkers: collagen 1, pSMAD2, and fibronectin (Figs. 4e, f; Extended Data Figs. 16e, f). Further analysis of fibrosis using the Sircol™ Collagen Assay shows the 185.2 μg/kg OA treatment significantly attenuates both soluble and insoluble collagen deposition, indicative of newly synthesized collagen and more mature crosslinked collagen, respectively (Figs. 4g, h). FVC (Fig. 5c) and static compliance (Extended Data Fig. 16c) were significantly improved with 185.2 μg/kg OA treatment, and respiratory mechanics show a less restrictive nature (Fig. 5d). We investigated the 185.2 μg/kg dose further through recovered bronchoalveolar lavage fluid (BALF) using cytokine array analysis, histological immunofluorescence and Sirius Red staining. Commonly implicated cytokines in the progression and severity of PF noted in patients and upregulated in the bleomycin model of PF including IL-6, TNF-α, and TIMP-1, were significantly reduced in 185.2 μg/kg treated BALF samples (Fig. 5i)^36–39^. Immunofluorescence imaging shows a marked reduction in TGF-β-related fibrotic markers collagen type I and α-smooth muscle actin (α-SMA) (Extended Data Fig. 16d). Sirius Red staining corroborates a reduction of histological total collagen levels (Extended Data Fig, 16g).

**Figure 5:**
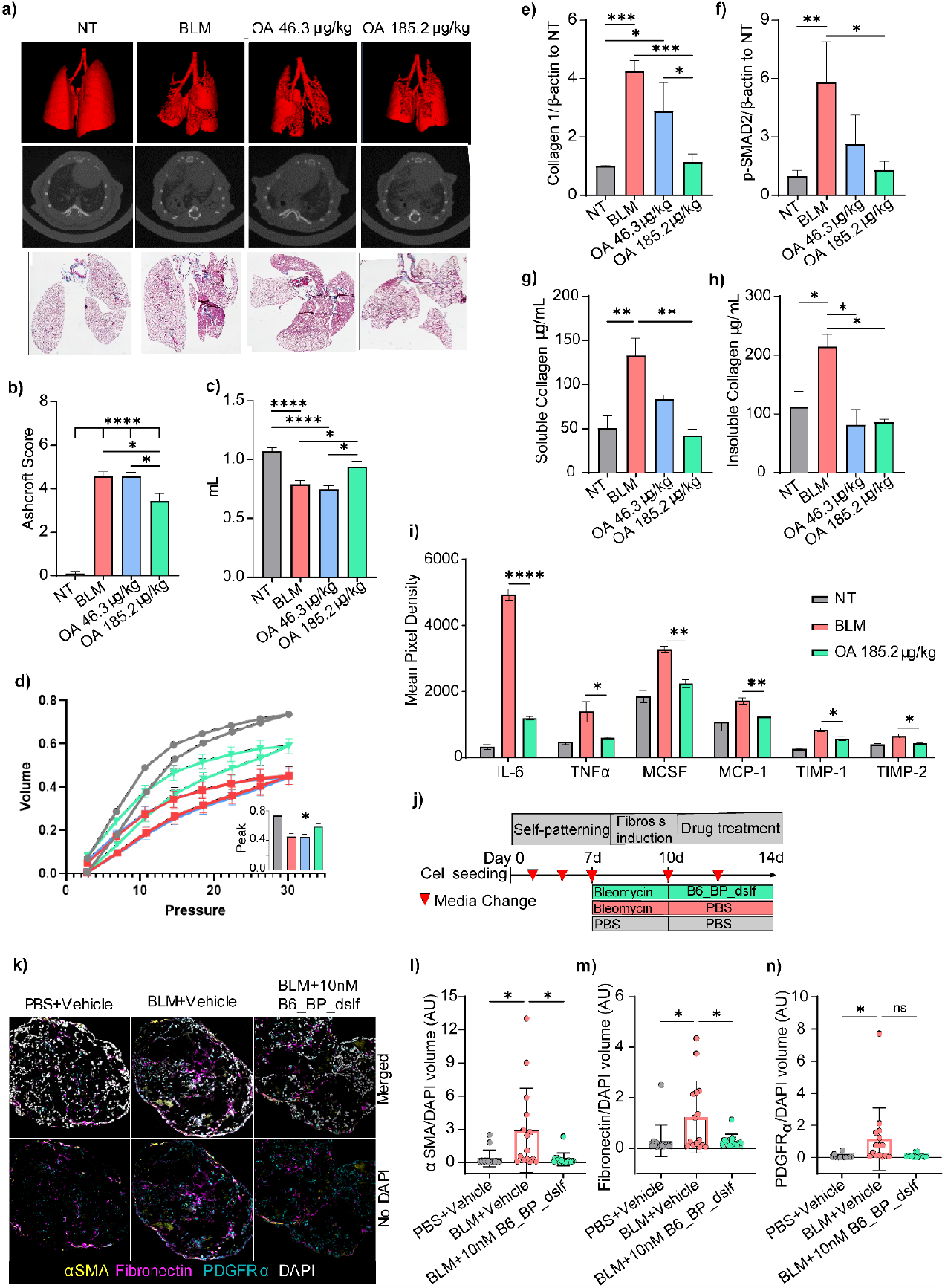
*In vivo* efficacy of OA-administered B6_BP_dslf in bleomycin-induced IPF. **a)** Three dimensional renderings of HR-uCT scans (top panel), representative HR-uCT scans (middle panel), and representative Masson-trichrome images for nontreated (NT), bleomycin treated (BLM) and inhaled B6_BP_dslf group show improvements in tissue health following B6_BP_dslf OA treatment. **b)** Average Ashcroft Scoring of Masson-trichrome images (mean ± SEM, NT n=5, BLM n=3, B6_BP_dslf 46.3 ug/kg n=3, B6_BP_dslf 185.2 ug/kg n=4) **c)** Forced vital capacity as measured by SCIREQ flexiVent FX (mean ± SEM, NT n=9, BLM n=10, B6_BP_dslf 46.3 ug/kg n=4, B6_BP_dslf 185.2 ug/kg n=6). **d)** Pressure-Volume curves measured by SCIREQ flexiVent with peak volumes in the inset graph (NT n=20, BLM n=14, B6_BP_dslf 46.3 ug/kg n=4, B6_BP_dslf 185.2 ug/kg n=7). **e)** Whole lung tissue homogenate western blot analysis of Collagen1 and **f)** p-SMAD2 show a dose-dependent reduction of these pro-fibrotic markers following B6_BP_dslf OA treatment (mean ± SD NT n=3, BLM n=3, B6_BP_dslf 46.3 ug/kg n=3, B6_BP_dslf 185.2 ug/kg n=3). **g)** Soluble and **h)** Insoluble collagen levels are lower following B6_BP_dslf OA treatment (mean ± SD NT n=3, BLM n=3, B6_BP_dslf 46.3 ug/kg n=3, B6_BP_dslf 185.2 ug/kg n=3). **i)** Cytokine array analysis of common cytokines implicated in inflammation and IPF (all NT vs. BLM are significant, p-value<.05, mean ± SD, NT n=4, BLM n=4, B6_BP_dslf 46.3 ug/kg n=4, B6_BP_dslf 185.2 ug/kg n=4). **j)** Time course of hFLO organoid growth, bleomycin induction, and 10 nM B6_BP_dslf treatment. **k)** Fluorescent confocal microscopy imaging of hFLO sections immunostained with pro-fibrotic markers αSMA, fibronectin, and PDGFRα. Volumetric analysis of **l)** αSMA, **m)** fibronectin, and **n)** PDGFRα normalized to DAPI signal (mean ± SD, N=15 cell aggregates were used per treatment, \**P*<0.05 determined using two-tailed Welch’s t-test).

With no significant changes in the total number of cells (Extended Data Figs. 16h-k) and reduction of inflammatory cytokines in BALF (Fig. 5i), inhaled B6_BP_dslf has potential to inhibit TGF-β induced fibrosis without exacerbating inflammation as compared to a systemically delivered antibody with a longer *in vivo* half-life^4^. A median mass aerodynamic diameter (MMAD) of ∼1-5 μm is necessary to reach the lower respiratory tract^40^ for an inhaled nebulized drug. To confirm the aerosol formulability, we nebulized B6_BP_dslf using an Aeroneb nebulizer and collected aerosols of two different MMADs: 2.5-4 μm and 4-6 μm. For both of these particle sizes, B6_BP_dslf is monomeric, hyperstable, and binds to αvβ6 with similar affinity pre-nebulization (Extended Data Fig. 17), paving the way for the development of this molecule as an inhaled nebulized therapy for IPF.

### Evidence of human efficacy using lung organotypic model

To assess the viability of B6_BP_dslf in a human organoid-based bleomycin system, we used the human fluorescent lung alveolar organoid (hFLO) triculture model as described previously ^41^. hFLO organoids were allowed to self-pattern for 7 days after which fibrosis was induced using bleomycin for 3 days prior to treatment with 10 nM B6_BP_dslf for 4 days. Upon immunofluorescent analysis, we observed increased pro-fibrotic markers α-SMA, fibronectin, and PDGFRα in bleomycin-treated organoids. Among these, we observed a statistically significant reduction in α-SMA and fibronectin levels in the organoids treated with B6_BP_dslf showing the efficacy of the treatment in lowering pro-fibrotic markers in human organoids (Fig. 5k-n).

## Discussion

The limited effectiveness of current treatments has renewed interest in developing inhaled therapeutics for IPF.^35,42^ To the best of our knowledge, the designed αvβ6 inhibitor described here (B6_BP_dslf) binds to its target with higher affinity and selectivity than any previously reported linear or cyclic peptide, or disulfide cross-linked knottin inhibitors of αvβ6,^43–47^ and is comparable to the leading antibody BG00011, which is no longer under development for IPF (Extended Data Table 2) due to exacerbation of the disease and death.^17,19^ While inhaled tissue-restricted delivery of drugs at the site of action minimizes overall toxicity, dose, and adverse effects, a challenge in development of inhaled biologics is instability or aggregation at the liquid-air interface. The properties of B6_BP_dslf are unchanged following aerosolization: the protein is monomeric, thermostable, has the same CD spectrum, and binds to αvβ6 with similar affinity. In addition to the highly potent anti-fibrotic effects demonstrated here in the bleomycin-induced lung fibrosis model, the reduced systemic exposure due to the short serum half-life (∼10 min), high selectivity and affinity for αvβ6, stabilization of the open αvβ6 conformation, ease of production using *E. coli*, hyper-thermostability, and aerosol formulatability give B6_BP_dslf an improved target product profile as a therapeutic candidate for IPF. Our designed αvβ6 inhibitor could also help combat progressive respiratory disease associated with current and future coronavirus infections.^48–51^ Further improvement in the pharmacokinetic/pharmacodynamic properties of these molecules can be achieved by site-selective chemical PEGylation or fusion to an immunoglobulin Fc domain for immune-oncology indications.^8,9^ More generally, a frequent challenge in drug development is the targeting of a single member within a large family of closely related proteins. This can be difficult to achieve with small molecules that share a conserved binding site, and the development of antibody panels capable of fine discrimination require considerable amounts of negative selection. Our structure-based *de novo* design strategy has high accuracy, as demonstrated by the cryoEM structures and achieves high selectivity by integrating both previously known binding motifs and introducing completely new interactions in a hyperstable small scaffold. As exemplified by our successful design of a potent and selective αvβ8 minibinder, this approach should be widely applicable to developing binders with high selectivity and affinity to individual members of the many therapeutically important families of cell surface receptors. Taken together, B6_BP_dslf, with its exquisite RGD integrin selectivity, hyperstability, *in vivo* efficacy in the bleomycin mouse model of lung fibrosis, and the potential for the inhaled route to deliver high drug concentrations to the lungs with minimal systemic exposure and thus provide an effective and safer treatment option for IPF patients than oral αvβ6 inhibitors, warrants further clinical investigation.

## Data availability

Structures from the first round design and B6B8_BP_dslf monomer have been deposited in the Protein Data Bank with accession numbers 7LMV and 7LMX, respectively. Structures of αvβ6+B6_BP_dslf and αvβ8+B8_BP_dslf have been deposited in the Electron Microscopy Data Bank (EMDB) with the accession numbers XXXX, XXXX, XXXX, and XXXX, and Protein Data Bank with the acession numbers XXXX and XXXX. Raw data has been deposited in the Electron Microscopy Public Image Archive (EMPIR) with the acession numbers XXXX, and XXXX. Raw counts/FastQ files for two rounds of directed evolution and model of the co-complex structures for some of the mutants are also uploaded on the github account (https://github.com/aroy10/avb6-publication). Any additional data is available from the authors upon request.

## Competing financial interests

A.R., L.S, X.D., J.L. T.S., D.B. are co-inventors on an International patent (Serial # PCT/US2020/057016) filed by University of Washington covering molecules and their uses described in this manuscript. C. O. is an employee of AstraZeneca and may own stock or stock options. M.G.C is an inventor on “Antibodies that bind integrin avb8 and uses thereof”, U.S. Patent US20210277125A1. A.R., H.B., J.C.K., M.S.S., M.C. and D.B. are inventors on a provisional patent describing the sequence and usage of αvβ8 integrins binders. A.R., L.S., J.C.K, H.B. and D.B. are co-founders of Lila Biologics and own stock or stock options in the company.

## Author contributions

A.R., L.S., J.C.K., L.J.S., T.S., A.C., A.F, M.G.C, P.V.R, G.R., and D.B designed the research. L.S. performed first and second round computational designs and affinity maturation of the first round design. A.R. designed the disulfide bonded version of the protein, performed directed evolution, generated material and reagents used for in vitro and in vivo experiments, performed biophysical characterization of the designed inhibitors, and wrote the manuscript. A.R., H.B. designed and characterized αvβ8 selective binders. M.S.S expressed and characterized all B8_BP_dslf variants and measured affinity and selectivity of those mutants against αvβ6 and αvβ8. X.D and T.S. provided the coordinates for human αvβ6 used for designing the inhibitor. X.D. and A.B. solved and refined the crystal structure BP and B6B8_BP_dslfide. X.D. determined affinities of the inhibitors against αvβ6 and αvβ8. J.L. measured binding of the final variants against all RGD binding integrins. A.F. optimized the expression and purification of the integrin samples used for all EM studies and prepared, collected, and processed all negative stain data. A.F. and M.G.C prepared, collected, and processed all cryoEM data, built the molecular models for the integrin-minibinder complexes, and analyzed all EM data. V.Q.L. tested inhibition of αvβ6 and αvβ8 mediated TGF-β activation by B6_BP_dslf using a CAGA co-culture assay. D.F, D.L., and C.O designed and performed the TMLC assay. TMLC assay was performed by C.O at AstraZeneca following standard MTA agreement. M.C.M. optimized expression, purification conditions and developed a one-step purification protocol for the B6B8_BP_dslf. L.C., R.R., C.M.C., T.P. performed large scale overexpression of the inhibitor. A.C. and R.V. performed the bleomycin induced PF model in mice. A.C., R.V., L.J.S., G.R., and P.V.R designed and analyzed the data for bleomycin induced PF models. P.V.R and G.R. supervised bleomycin induced PF model study. M.C. performed the binding of the designed inhibitor to A431 cells and in vivo imaging and analyzed data. J.C. designed and supervised the in vivo imaging experiments. A.R., H.B., and J.C.K. developed the sandwich ELISA bioassay for quantifying minibinders in biological samples; J.C.K. and S.C. ran the sandwich ELISA on serum and lung samples. A.R. and J.C.K. designed and analyzed the B6_BP_dslf_disfulf pharmacokinetics in mice and the h-LAP_1_ binding assay for integrins αvβ6 and αvβ8. Y.K. helped with the generation of the SSM library for the designed inhibitor. D.B. supervised, oversaw research, and coordinated research presented here. A.R., J.C.K., L.J.S., A.C., M.G.C. T.S. and D.B. wrote the paper with input from P.V.R. and G.R. Inputs from all authors were included in the manuscript as well.

## Acknowledgements

Funding for this research was supported by the National Institutes of Health under Grants No. R01 GM092802, R01 AR067288 and R35 GM147414. The US government has certain rights in inventions described here. A.R. and H.B. are Washington Research Foundation Innovation Postdoctoral Fellows at Institute of Protein Design and would like to acknowledge kind financial support provided by Washington Research Foundation. Funding support was also provided by The Audacious Project at the Institute for Protein Design (D.B., L.C., R.R., C.C.), Alexandria Venture Investments Translational Investigator Fund (A.R.), Washington State Supplement Funding to Support the Institute for Protein Design (L.J.S.), the Washington Research Foundation Translational Research Grants (D.B., A.R., P.V.R., and R.V.W.), the Howard Hughes Medical Institute (D.B. and Y.K.), Medimmune (C.K.S, D.K.F, and D.L.), and the Pew Biomedical Scholars award (M.G.C) V.Q.L. was supported by NIH K01DK124443 and A.F. was supported by NIH 5T32 GM008268-33. We would also like to thank Cassie Bryan, Parisa Hosseinzadeh, Karla-Luise Herpoldt, Franziska Seeger, George Ueda, Brian Weitzner and many other Baker lab members for useful discussion and support. Computing resources were provided by the volunteers who have donated the spare CPU cycles of their cellular telephones and computers to the Rosetta@Home project; the Hyak supercomputer at the University of Washington; and the Rhino cluster at the Fred Hutchinson Cancer Center. We would like to thank Luki Goldschmidt, Patrick Vecchiato, and Ben McGough for IT support, Liz Soberg for assistance with animal studies, and Caleigh Azumaya and Anvesh Dasari for assistance with cryoEM data collection setup. Electron microscopy data were generated using the Fred Hutchinson Cancer Center Electron Microscopy shared resource, supported in part by the Cancer Center Support Grant P30 CA015704-40. We would like to thank Jessica M. Snyder, D.V.M., at the Department of Comparative Medicine at University of Washington for conducting the histological scoring.

