## Supplementary for "*De novo* design of highly selective miniprotein inhibitors of integrins αvβ6 and αvβ8"

Extended Data Figures, Tables, and Legends

Extended Data Fig. 1: Crystal structures of designed αvβ6 inhibitors from first and second round design strategies.
Extended Data Fig. 2: Metal-dependent binding of designed proteins to human αvβ6.
Extended Data Fig. 3: Flow cytometry screening of second-round designed protein binders using human αvβ6.
Extended Data Fig. 4: Sorting scheme for the site-saturation mutagenesis (SSM) library of av6_3.
Extended Data Fig. 5: Experimental optimization of av6_3 for binding and heatmap of key evolved residues.
Extended Data Fig. 6: BLI binding of purified mutants against titrations of biotinylated human αvβ6/αvβ8.
Extended Data Fig. 7: Selectivity of designed αvβ8 inhibitors against αvβ6 and αvβ8 using yeast surface display.
Extended Data Fig. 8: Selectivity of designed αvβ6 inhibitors against other RGD-binding integrins.
Extended Data Fig. 9: Thermal stability of B6_BP_dslf under reducing and non-reducing conditions.
Extended Data Fig. 10: TMLC and CAGA-reporter co-culture assays assessing αvβ6-mediated inhibition of TGF-β1.
Extended Data Fig. 11: *In vivo* imaging of αvβ6 (+) A431 tumors using fluorescently-labeled B6_BP.
Extended Data Fig. 12: Lung and serum pharmacokinetics of B6_BP_dslf in healthy male C57BL/6 mice.
Extended Data Fig. 13: HR-μCT imaging, histopathology, and lung function after IP B6_BP_dslf in "mild" bleo-model.
Extended Data Fig. 14: CT frequency, lung function, and cellular responses after IP B6_BP_dslf in "mild" bleo-model.
Extended Data Fig. 15: HR-μCT imaging and lung function after IP B6_BP_dslf in "severe" bleo-model.
Extended Data Fig. 16: Lung function and cellular responses after OA B6_BP_dslf in "severe" bleo-model.
Extended Data Fig. 17: Stability of B6_BP_dslf following nebulization.
Extended Data Fig. 18: Negative stain EM class averages, SDS-PAGE, SEC, and CryoEM micrographs and statistics.
Extended Data Fig. 19: Processing schematic for cryoEM of αvβ6 + B6_BP_dslf and αvβ8 + B8_BP_dslf complexes.
Extended Data Table 1: Kinetic analysis of BLI binding of purified mutants against titrations of biotinylated human αvβ6.
Extended Data Table 2: Affinity and selectivity comparison of B6_BP against other leading reported αvβ6 inhibitors.
Extended Data Table 3: Predicted and observed interactions between integrins αvβ6/αvβ8 and designed minibinders.
Supplementary Table 1: Statistics of X-ray diffraction and structure refinement.
Supplementary Table 2: Exclusion criteria for animals in bleomycin-induced lung fibrosis model.
Supplementary Table 3: Sequences of all designs and evolved variants reported in this paper.
Supplementary Table 4: Forward mutagenic primer sequences used for directed evolution.
Supplementary Table 5: Reverse mutagenic primer sequences used for directed evolution.
Supplementary Table 6: Integrin αvβ6 and αvβ8 headpiece and ectodomain amino acid sequences.
Supplementary Table 7: CryoEM data collection, refinement, and validation statistics.
Materials and Methods
References

### Extended Data Figures

a.

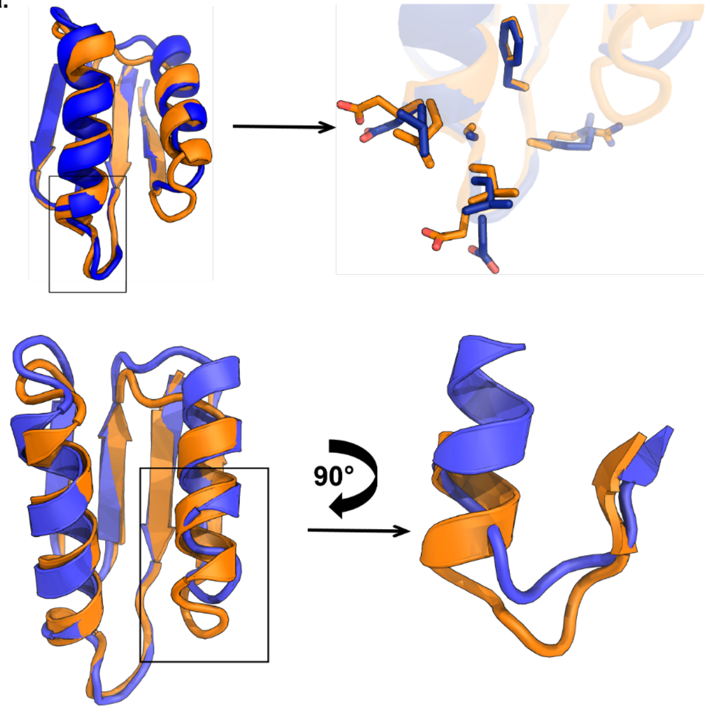

b.

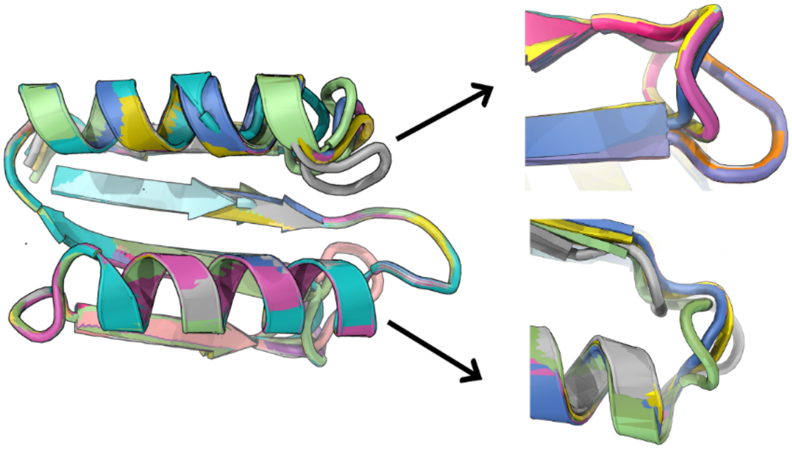

c)

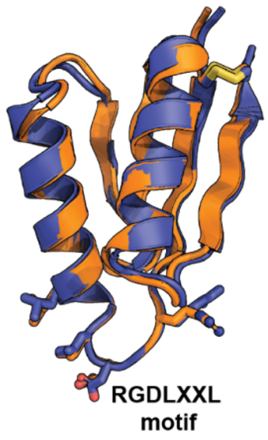

**Extended Data Fig. 1: Crystal structures of designed αvβ6 inhibitors from first and second round design strategies. a)** Crystal structure (blue) of the evolved variant from the first round of design superimposed onto the design model (orange). Although the first part of the crystal structure including the RGD loop (Top panel) overlaid well with the design model, there was rigid body movement of the C-terminal helix of the fold equivalent to one helical turn. **b)** For the second generation of designs, the crystal structure of the previous round was superimposed onto αvβ6 by aligning the RGD motif. Two loops were sampled for length and conformation and 16 designs were ordered in the second round. **c)** Superposition of the designed B6B8_BP_dslf (orange) and the crystal structure (blue) in the cartoon model. The designed disulfide bond and the

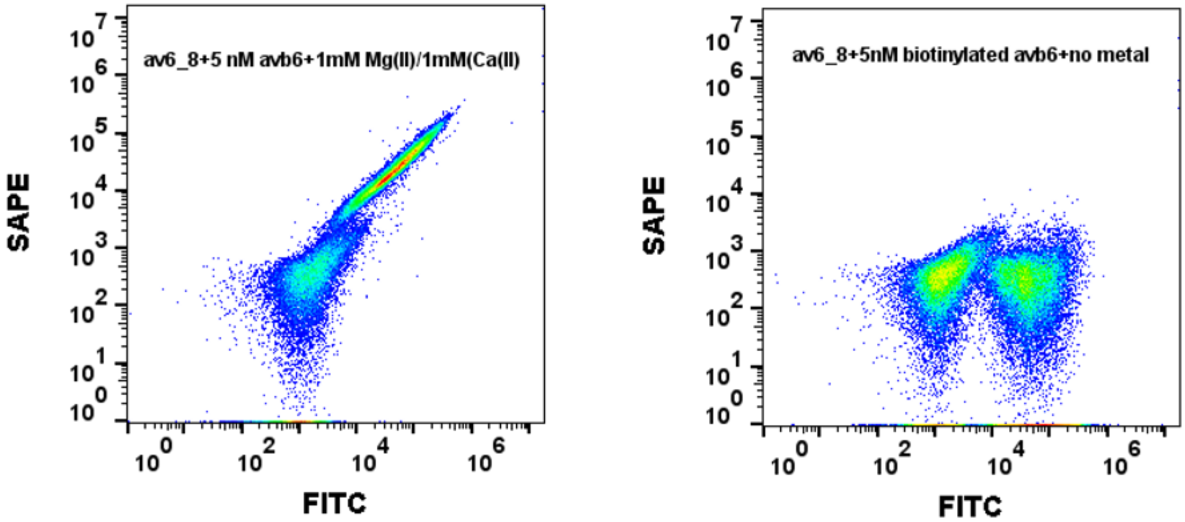

RGDLXXL motif is shown in stick representation.

**Extended Data Fig. 2: Metal-dependent binding of designed proteins to human αvβ6.** The designed protein shows metal dependent binding to αvβ6. In the absence of any metal, there is no detectable binding (left panel) as compared to in presence of 1mM Ca(II)/1mM Mg(II) (right panel).

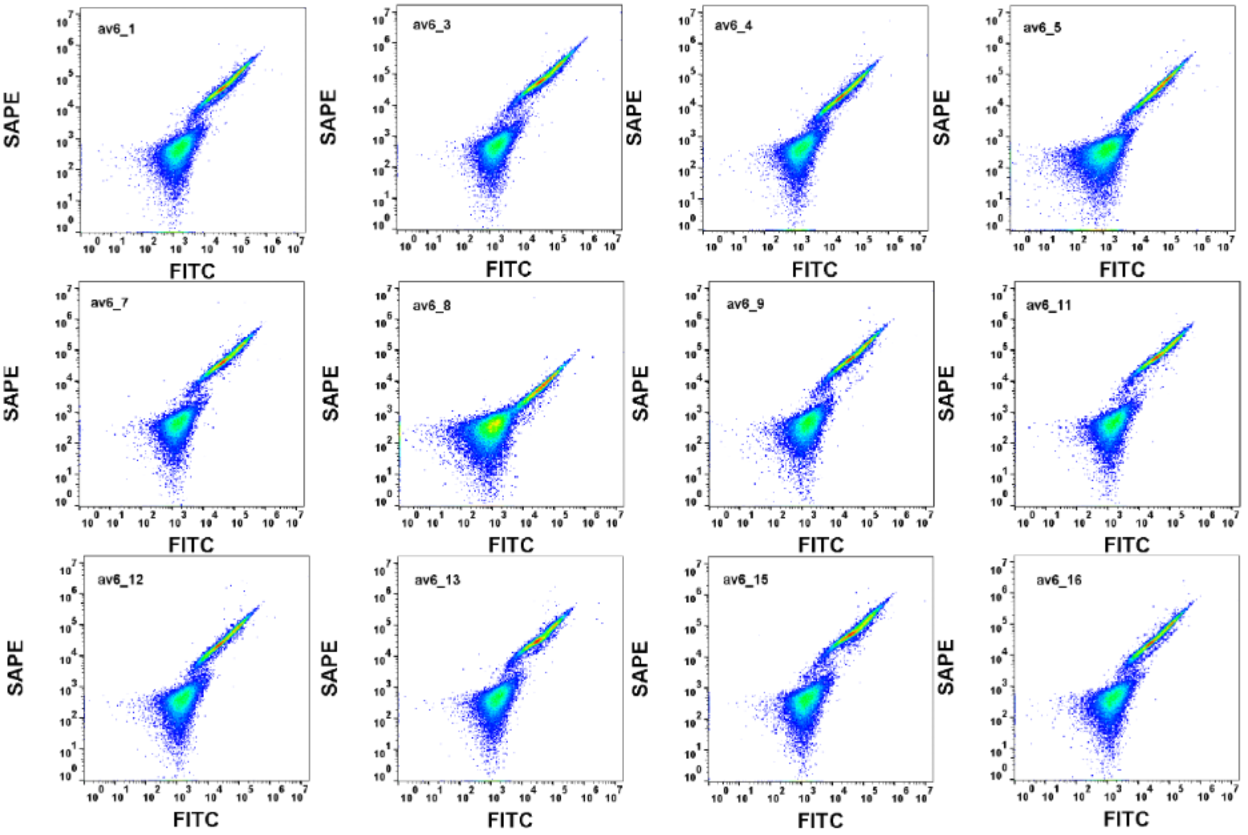

av6_1
av6_3
av6_4
av6_5
av6_7
av6_8
av6_9
av6_11
av6_12
av6_13
av6_15
av6_16
SAPE
FITC

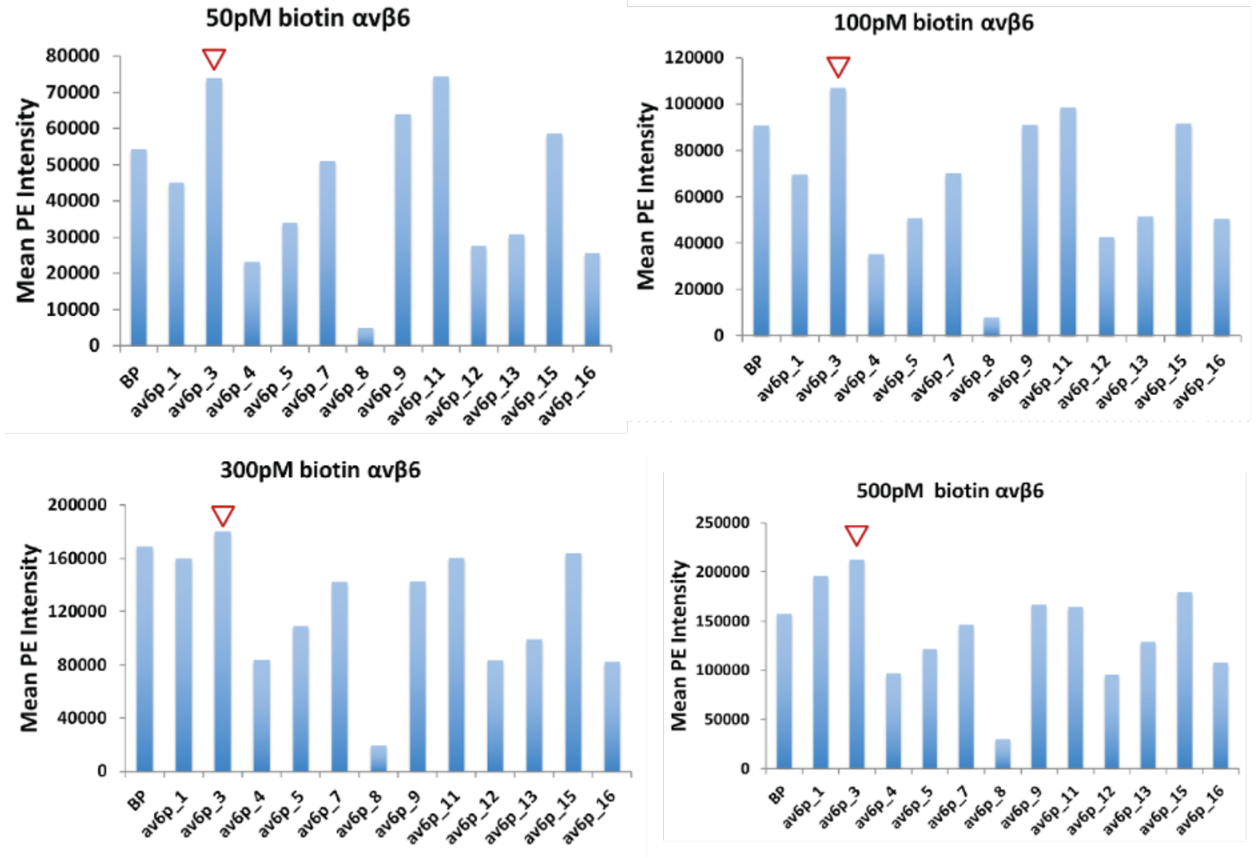

50pM biotin αvβ6
100pM biotin αvβ6
300pM biotin αvβ6
500pM biotin αvβ6
Mean PE Intensity
BP
av6p_1
av6p_3
av6p_4
av6p_5
av6p_7
av6p_8
av6p_9
av6p_11
av6p_12
av6p_13
av6p_15
av6p_16

**Extended Data Fig. 3: Flow cytometry screening of second-round designed protein binders using human αvβ6.** (Top Panel) Designs were expressed using yeast surface display technique and incubated with 50 pM of biotinylated human αvβ6. (Bottom Panel) Mean PE intensity of BP (binding protein) and all second round designs at 50 pM, 100 pM, 300 pM, and 500 pM of biotinylated integrin αvβ6. av6_3 shows the highest binding signal (PE fluorescence signal) across all concentrations tested.

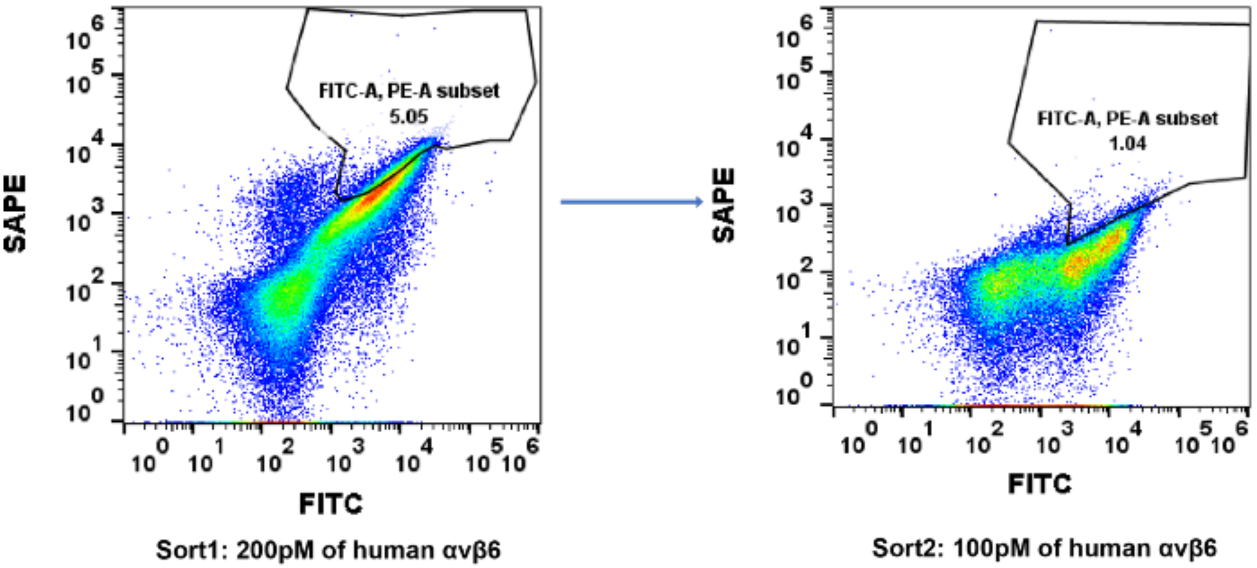

**Extended Data Fig. 4: Sorting scheme for the site-saturation mutagenesis (SSM) library of av6_3.** For the first round of selection, the library was incubated with 200 pM of biotinylated human αvβ6 and the top 5% of the binders were collected. For the second and final round of selection, 100 pM of biotinylated αvβ6 was used and the top ~1% of the binding population was collected.

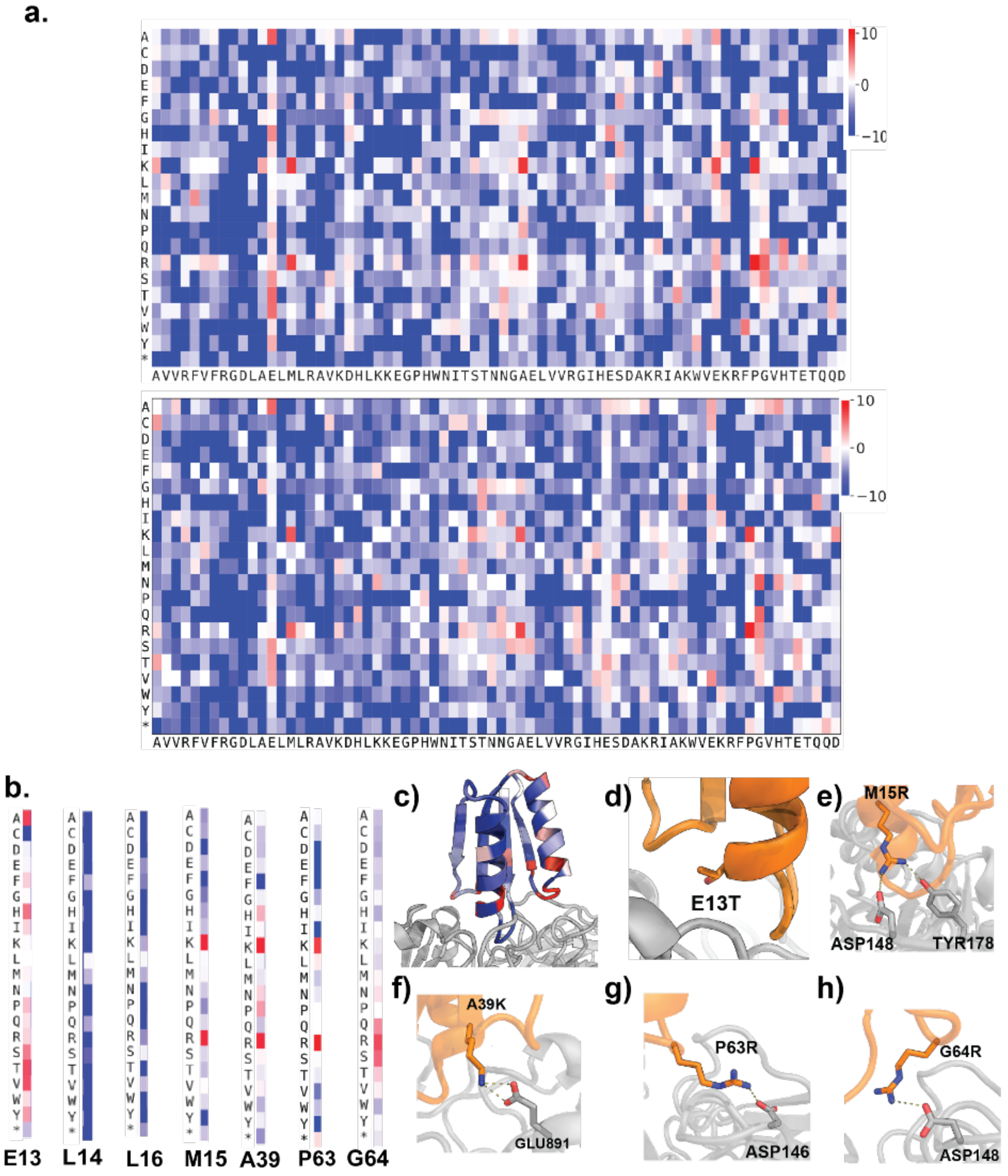

**Extended Data Fig. 5. Experimental optimization of av6_3 for binding and heatmap of key evolved residues.** a) Enrichment ratio of the evolved variants compared to the naive library after two rounds of sorting with increasing stringency (200 pM and 100 pM) from biological replicates. Beneficial mutations are colored in red and deleterious mutations are colored in blue. b) Heat map of key enriched and conserved residues derived from SSM optimization of the inhibitor. E13 prefers to be small polar residues or small hydrophobic residues. L14 and L16 of the RGDLXXL motif are highly conserved, demonstrating the importance of the LXXL motif. M15/A39/P63/G64 mutations are mainly charge-complementary to the receptor. c) The majority of the enriched substitutions increase charge complementarity to the receptors. d) E13 prefers to be small polar or hydrophobic. e) M15R/K is well within hydrogen bonding interaction range to D148 and Y178 of αvβ6. f) A39K substitution facing the β6 subunit likely introduces a salt bridge with Glu891. g,h) Two consecutive residues (P63 and G64) on Loop2 facing the αv subunit are enriched as positively charged Lys or Arg residue, which likely form salt bridges with two acidic residues D146 and D148 on the αv subunit of the receptor, respectively.

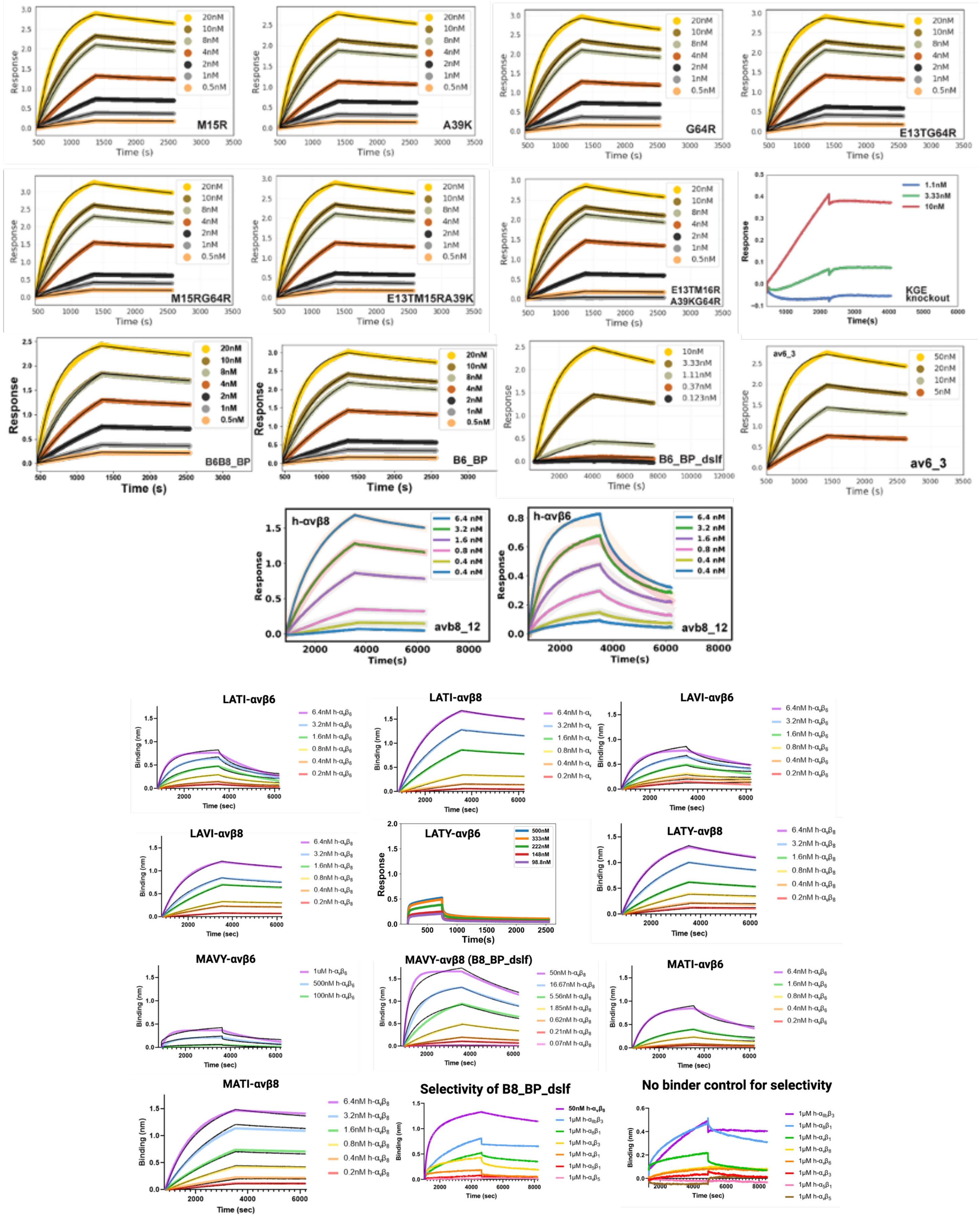
M15R
A39K
G64R
E13TG64R
M15RG64R
E13TM15RA39K
E13TM16R A39KG64R
KGE knockout
B6B8_BP
B6_BP
B6_BP_dslf
av6_3
h-αvβ8
h-αvβ6
avb8_12
LATI-avβ6
LATI-avβ8
LAVI-avβ6
LAVI-avβ8
LATY-avβ6
LATY-avβ8
MAVY-avβ6
MAVY-avβ8 (B8_BP_dslf)
MATI-avβ6
MATI-avβ8
Selectivity of B8_BP_dslf
No binder control for selectivity
Response
Time (s)
Binding (nm)
Time (sec)

**Extended Data Fig. 6: Bio-layer interferometry (BLI) binding of all purified mutants against titrations of human αvβ6, αvβ8, and other RGD integrins. (a)** See Table S1 for kinetic analysis values. All panels are binding against αvβ6 titrations, except the bottom left panel that shows binding of avb8_12 to αvβ8. **(b)** Binding for the B8_BP_dslf point mutants and selectivity for B8_BP_dslf for RGD integrins.

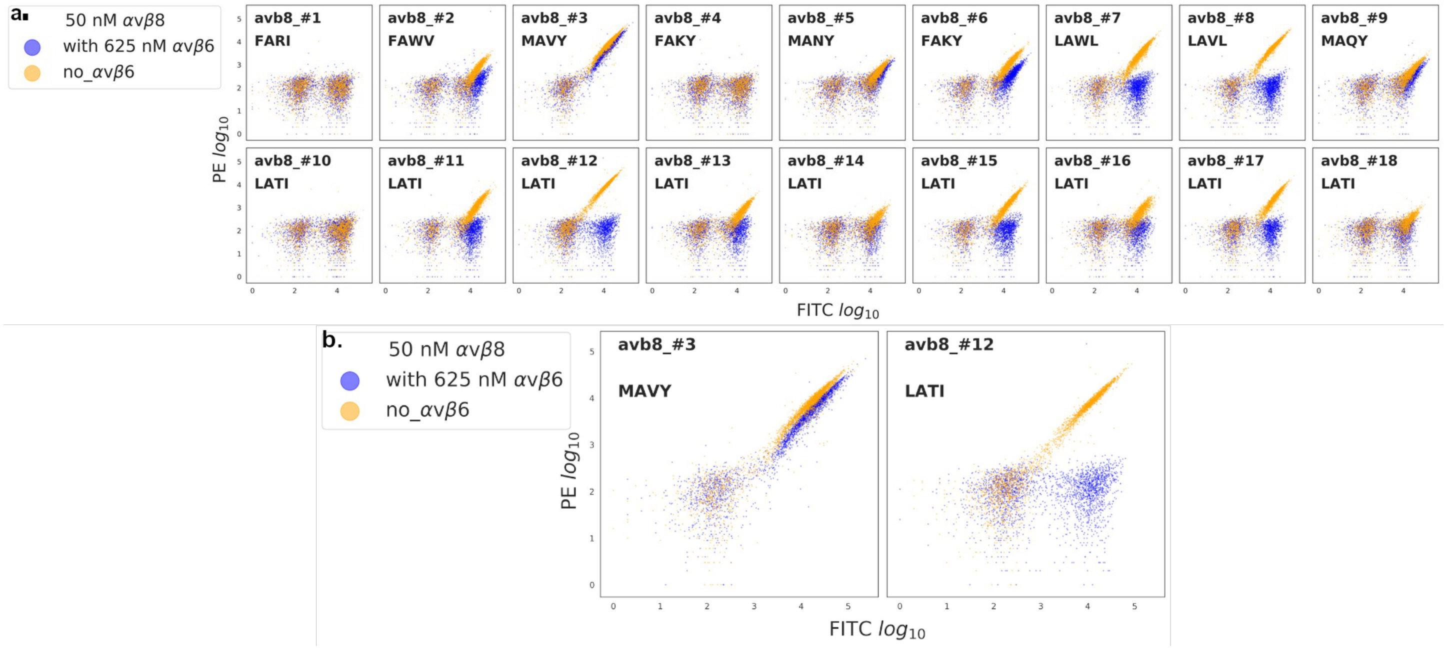

**Extended Data Fig. 7: Competition assay using yeast surface display technique to determine selectivity of designed αvβ8 binders. a)** Designs were expressed using yeast surface display technique and incubated with 50 nM of FLAG-tagged αvβ8 and 625nM of unlabelled αvβ6. Binding was detected by anti-FLAG-PE conjugated antibody from abcam (Product # ab72469). **b)** Effect of -RGDLXXL motif on selectivity of the designed binders. Mutation of -RGDMAVY motif (avb8_#3 or B8_BP_dslf) to -RGDLATI (avb8_#12) completely abrogates selectivity towards αvβ8.

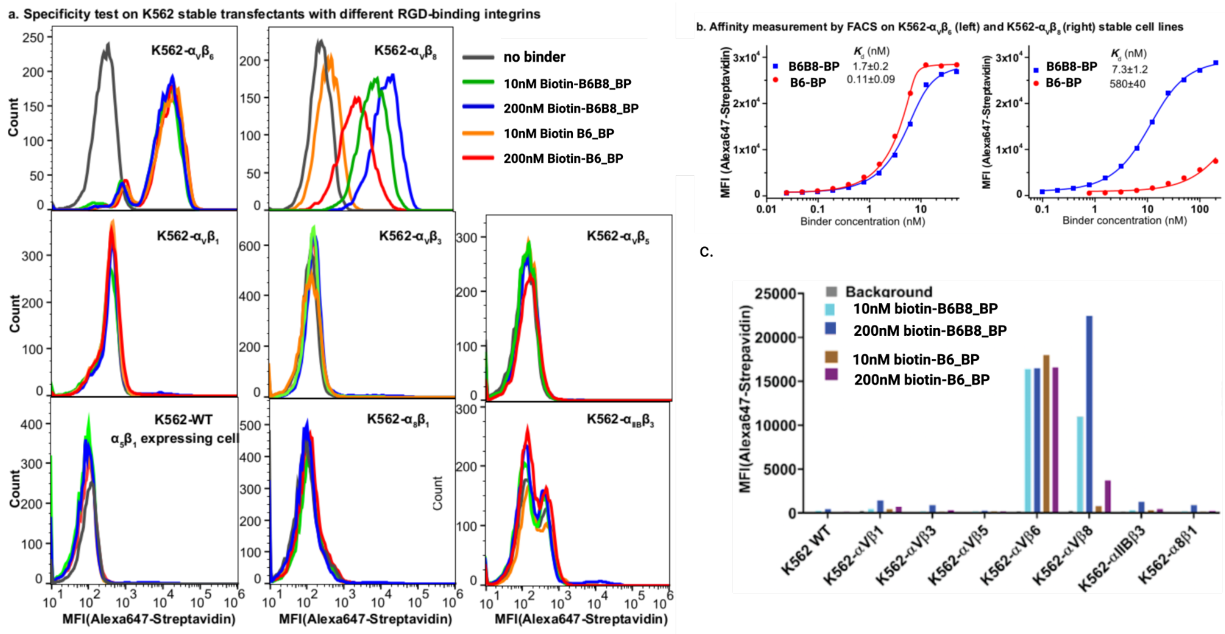

**Extended Data Fig. 8: Selectivity of designed αvβ6 inhibitors against other RGD-binding integrins. a)** Selectivity of B6B8_BP and B6_BP against other RGD-binding integrins: B6B8_BP/B6_BP does not bind to αvβ1, αvβ3, αvβ5, α5β1, α8β1, and αiibβ3 integrins up to 200 nM concentration. Assays were performed using biotinylated B6B8_BP/B6_BP via an N-term Avitag and K562 cells stably transfected with different integrins. All the incubation and washes were done in the assay buffer, L15 medium with 1% BSA. In detail, cells were incubated with biotinylated binders with indicated concentrations at 22℃ for 1h, followed by three washes and staining with Alexa-647 streptavidin for 20 minutes, followed by another three washes and subject to flow cytometry. **b)** Dose-dependent titration of B6B8_BP and B6_BP on K562 cells stably transfected with αvβ6 (left) and αvβ8 (right) with the same method described in panel a. B6B8_BP and B6_BP binds to αvβ6 with $K_d$ values of 1.7 (± 0.2) and 0.11 (± .09) nM, respectively, and binds to αvβ8 with $K_d$ values of 7.3 (± 1.2) and 580 (±40) nM, respectively. **c)** Selectivity of B6B8_BP and B6_BP against 8 -RGD binding integrins: K562 cells stably transfected with different integrins were incubated with biotinylated B6B8_BP and B6_BP followed by staining with Alexa-647 streptavidin. B6_BP binding is highly specific to αvβ6, whereas B6B8_BP binds to both αvβ6 and αvβ8. Both B6B8_BP and B6_BP show negligible binding up to 200 nM of concentration against other 6 integrins.

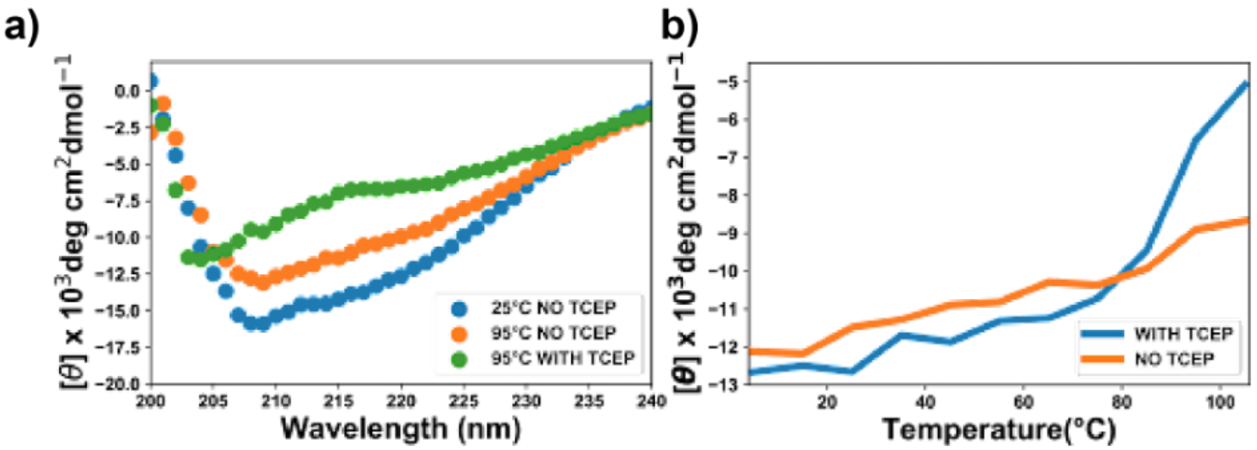

**Extended Data Fig. 9: Thermal stability of B6_BP_dslf under reducing and nonreducing conditions.** a) Circular dichroism (CD) wavelength scan of B6_BP_dslf at 25°C and 95°C in the presence of 1 mM reducing agent (TCEP). b)

Thermal melt of B6_BP_dslf monitored by CD spectra in the presence/absence of a reducing agent. B6_BP_dslf maintains its signature CD spectra consistent with a mixed alpha/beta fold under non-reducing conditions. Under reducing conditions, B6_BP_dslf melts ~90°C indicating hyper-thermostability is partially mediated by the engineered disulfide bond.

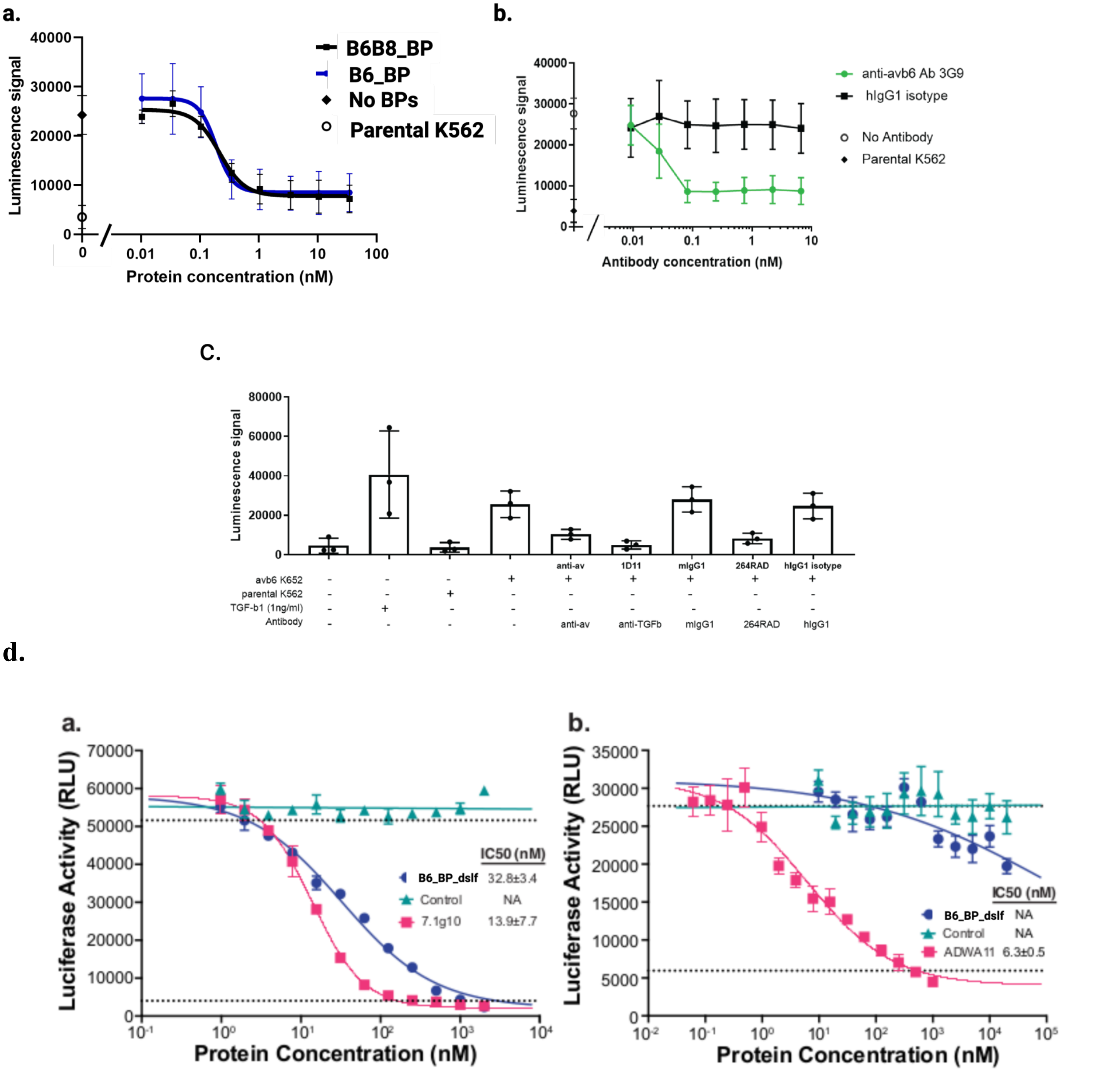

**Extended Data Fig. 10: TMLC and CAGA-reporter co-culture assays assessing αvβ6-mediated inhibition of TGF-β1.** a) TGF-β inhibition mediated by B6B8_BP and B6_BP in TMLC:αvβ6 K562 co-culture assay. Both B6B8_BP and B6_BP blocks αvβ6 mediated TGF-β activation with similar $IC_{50}$ values (216 pM and 188 pM, respectively). b) Positive and negative controls for αvβ6-mediated TGF-β activation in TMLC:αvβ6 K562 co-culture assay. An αvβ6-specific antibody 3G9 (known clinically as STX-100 and BG00011) can inhibit TGF-β activation in the TMLC assay, whereas the human IgG1 isotype control has no effect on αvβ6 mediated TGF-β activation. c) Additional controls for TMLC assay: Recombinant TGF-β induces a clear increase in luciferase activity, showing that the TMLCs can respond to active TGF-β.

Anti TGF-β, anti-αv, and an anti-αvβ6/β8 (264-RAD) are capable of blocking TGF-β activation in the TMLC assay. See methods section for detailed description for each antibody used in this assay. a,b) B6_BP_dslf selectively inhibits αvβ6-mediated TGF-β1 activation. αvβ6 (a) and αvβ8 (b) transfectants were co-incubated with CAGA-reporter cells and GARP/TGF-β1 transfectants and inhibitors. Control: irrelevant nanobody. 7.1g10: inhibitory antibody to integrin β6. ADWA11, an inhibitory antibody to integrin β8. Mean ± SD are plotted, and $IC_{50}$ values for each inhibitor are reported. Lower dashed lines represent integrin-independent TGF-β1 activation, which is 7.8% of the total activation with αvβ6 transfectants (upper dashed line in a) and 13.3% of the total activation with αvβ8 transfectants (upper dashed line in b).

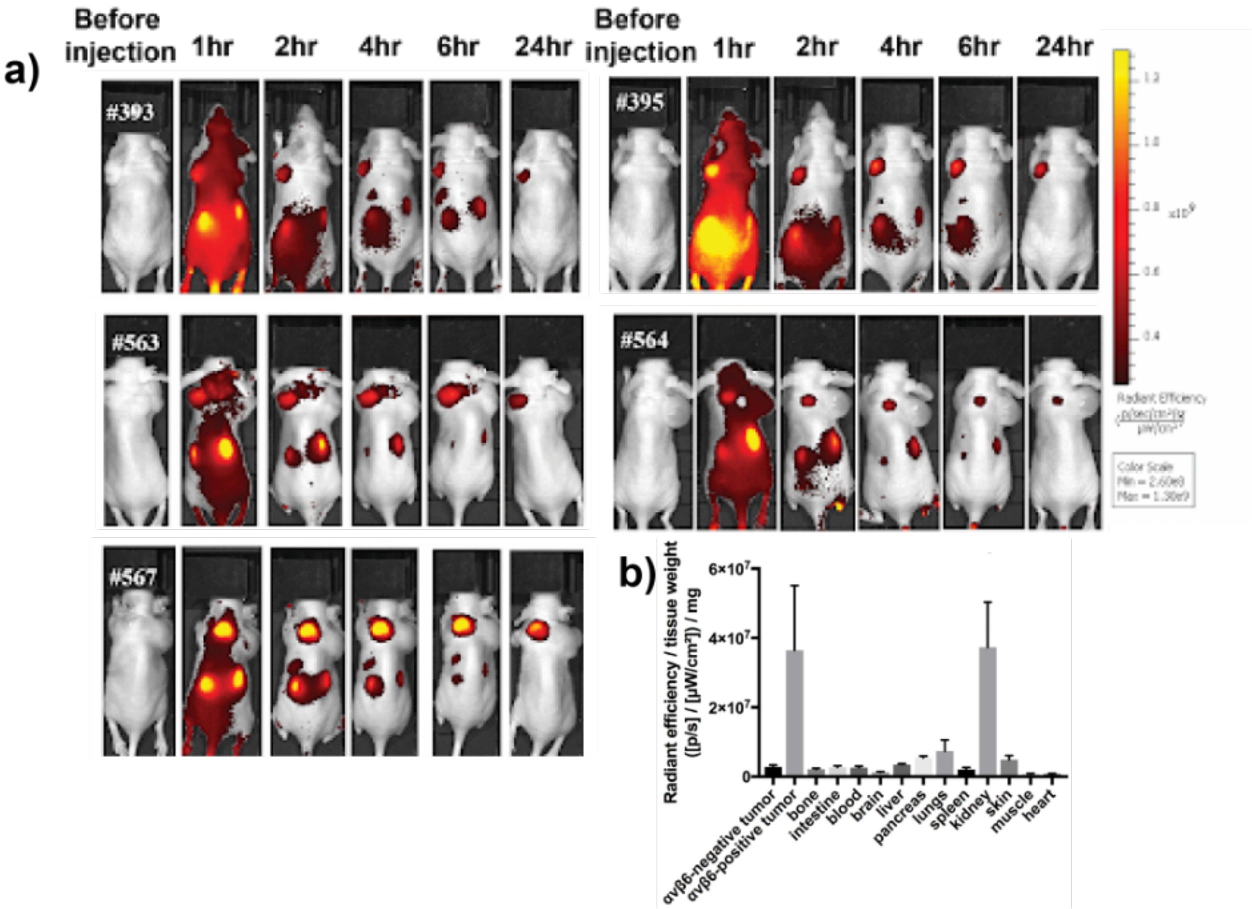

**Extended Data Fig. 11: *In vivo* imaging of αvβ6 (+) A431 tumors using fluorescently-labelled B6_BP.** a) Imaging αvβ6 (+) tumors *in vivo*; athymic nude mice (n=5) were injected with αvβ6 (+) A431 cells on the left shoulder and αvβ6 (-) HEK293T cells on the right shoulder. AlexaFluor-680-labelled B6_BP (AF680-B6_BP) was injected via tail vein to image the tumors over time as indicated. b) Semiquantitative *ex vivo* biodistribution assay of AF-680-B6_BP at 6 hours post-tail vein injection. B6_BP selectively accumulates in αvβ6 (+) tumors and primarily clears via glomerular filtration in the kidneys.

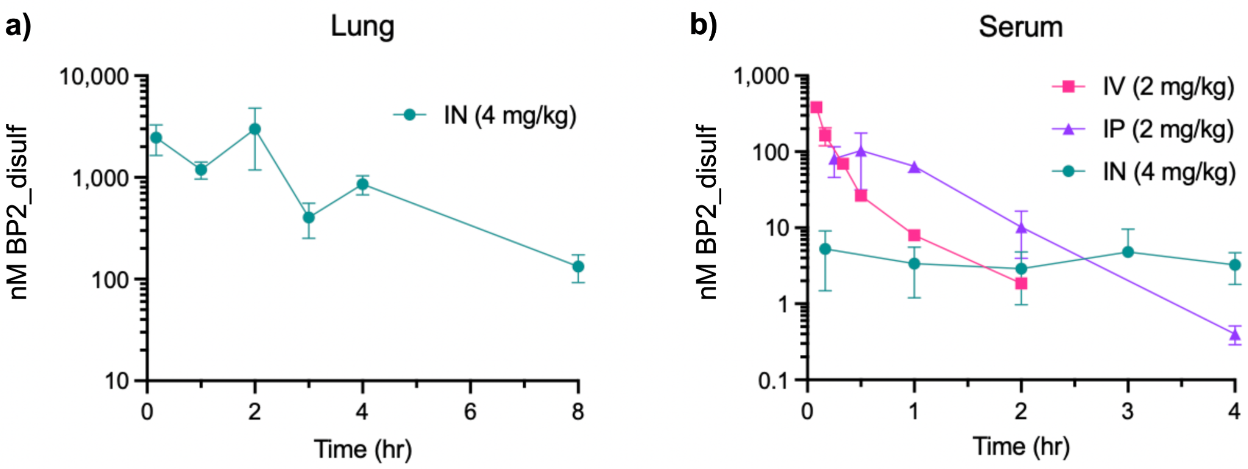

c)

| Biological sample | BP2_disulf dose (mg/kg) | Route of administration | Terminal elimination half-life ($t_{1/2,z}$) |
|---|---|---|---|
| Lung | 4 | Intranasal (IN) | 55.2 ± 15.6 min |
| Serum | 2 | Intravenous (IV) | 10.5 ± 0.8 min |
| Serum | 2 | Intraperitoneal (IP) | 11.2 ± 1.8 min |

**Extended Data Fig. 12: Lung and serum pharmacokinetics of B6_BP_dslf in healthy male C57BL/6 mice following a single intravenous or intraperitoneal 2 mg/kg dose, or intranasal 4 mg/kg dose.** To characterize the pharmacokinetics of B6_BP_dslf following different routes of administration, 8-12 week old male C57BL/6 mice were given 2 mg/kg B6_BP_dslf via the intravenous (IV) or intraperitoneal (IP) routes, or 4 mg/kg B6_BP_dslf via the intranasal (IN) route. B6_BP_dslf concentrations were quantified in (a) lung tissue and (b) serum samples collected at different time points post-administration using a sandwich enzyme linked immunosorbent assay (ELISA) method. (a,b) Following intranasal administration, serum concentrations were on average 484-fold lower than the lung concentrations over 4 h. (b) The area under the curve from 0 to 2 hours ($AUC_{0\text{-}2hr}$) following IP administration was ~2-fold higher than the $AUC_{0\text{-}2hr}$ from the IV route (7,569.7 ± 1,055.6 nM•min vs. 3,892.9 ± 925.4 nM•min). (c) Terminal elimination half-lifes ($t_{1/2,z}$) of B6_BP_dslf in lung and serum following different routes of administration. The elimination of B6_BP_dslf from the lungs within hours and low relative serum concentrations compared to lung concentrations following IN administration is consistent with the lung and serum pharmacokinetics observed for intranasal GSK3008348 in healthy C57BL/6 mice[1].

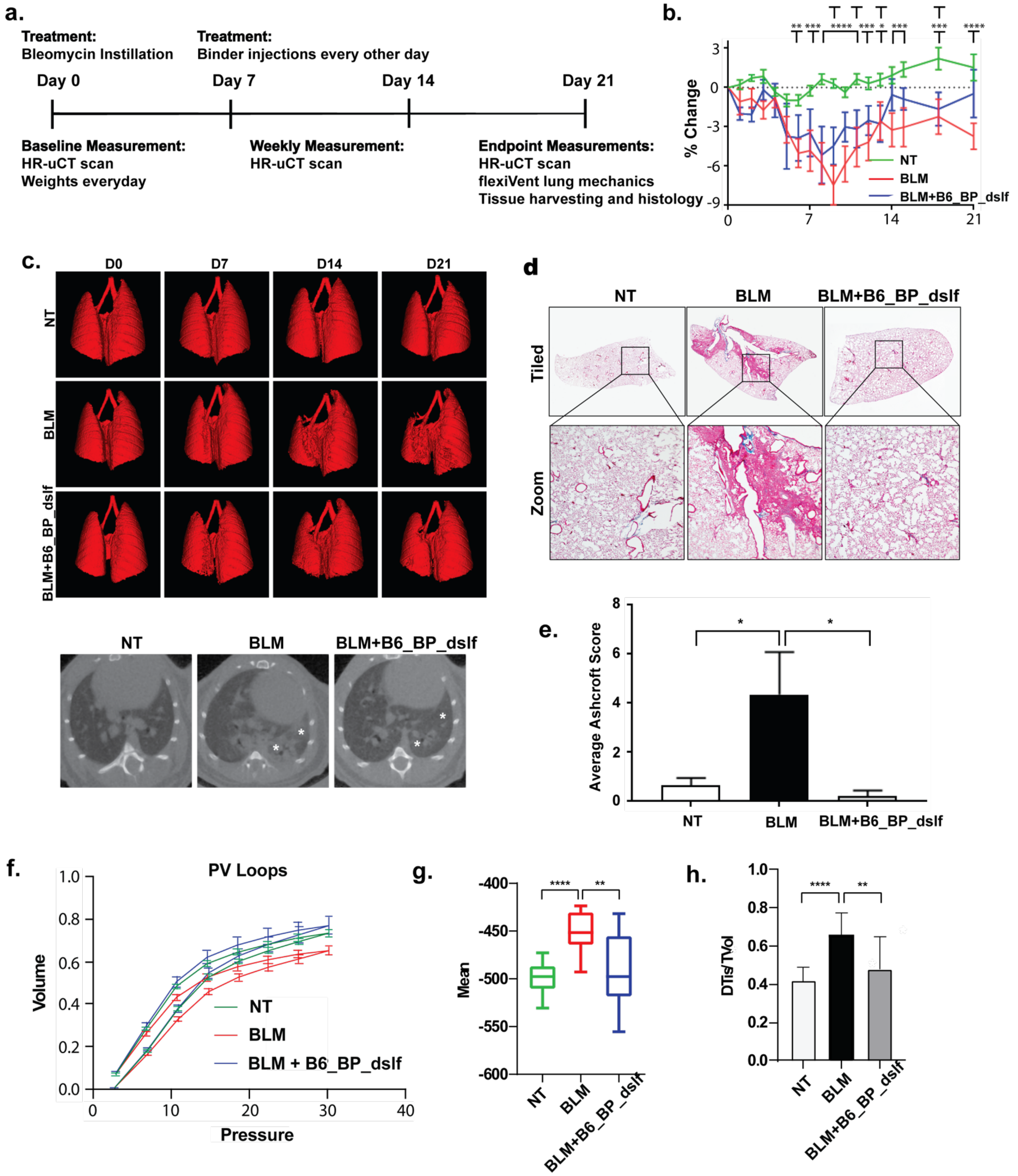

**Extended Data Fig. 13: High-resolution micro-computed tomography (HR-μCT) imaging, histopathology, and lung function following B6_BP_dslf intraperitoneal (IP) administration in the "mild" bleomycin-induced pulmonary fibrosis mouse model.** In the "mild" bleomycin mouse model, lidocaine was not used to relax the vocal cords. a) Treatment regimen for bleomycin-induced pulmonary fibrosis. Mice were intratracheally administered bleomycin at 1 U/kg body weight. Mice were injected intraperitoneally with B6_BP_dslf binder at 100 μg/kg every-other-day starting at day 7 post-bleomycin instillation, and ending on day 19, for a total of 7 treatments administered as compared to the healthy non-treated (NT) group. b) Weight changes for NT, BLM and BLM+B6_BP_dslf groups (NT n=24; BLM n=14; BLM+B6_BP_dslf n=5). Mice treated with B6_BP_dslf show a steady weight gain after day ~8 and no longer have a significant weight loss compared to NT by day 21, while mice receiving only bleomycin continue to have significant weight loss by day 21. c) Upper and lower panel: High-resolution micro-CT scans with asterisks at fibrotic areas in the lower lobes lung. HR-µCT of mice treated with B6_BP_dslf intraperitoneally shows more healthy tissue present, indicating that B6_BP_dslf halted the progression of fibrosis (Non-treated 611.1, BLM 429, B6_BP_dslf 542.2 mm$^3$). d) Masson-Trichrome staining with boxes indicating regions of higher magnification for NT, BLM and BLM+B6_BP_dslf treated mice. Fibrous thickening of alveolar space can be clearly seen in BLM treated mice as compared to the NT group (middle panel). e) Gradation of fibrotic burden: Average Ashcroft Scoring of histological slides for NT (0.66), BLM (4.347) and BLM+B6_BP_dslf (0.22) treated mice. B6_BP_dslf treatment changes the ashcroft score from 4.347 to 0.22 as compared to the BLM treated group. f) Representative pressure-volume loop curves (PV-loops) for NT, BLM and B6_BP_dslf groups. In all the measurements, B6_BP-disulf treatment improves lung function and nears the NT group as compared to BLM treated mice only. g,h) Mean intensities of micro-CT scan (g) and quantification of lung density from micro-CT scans (h).

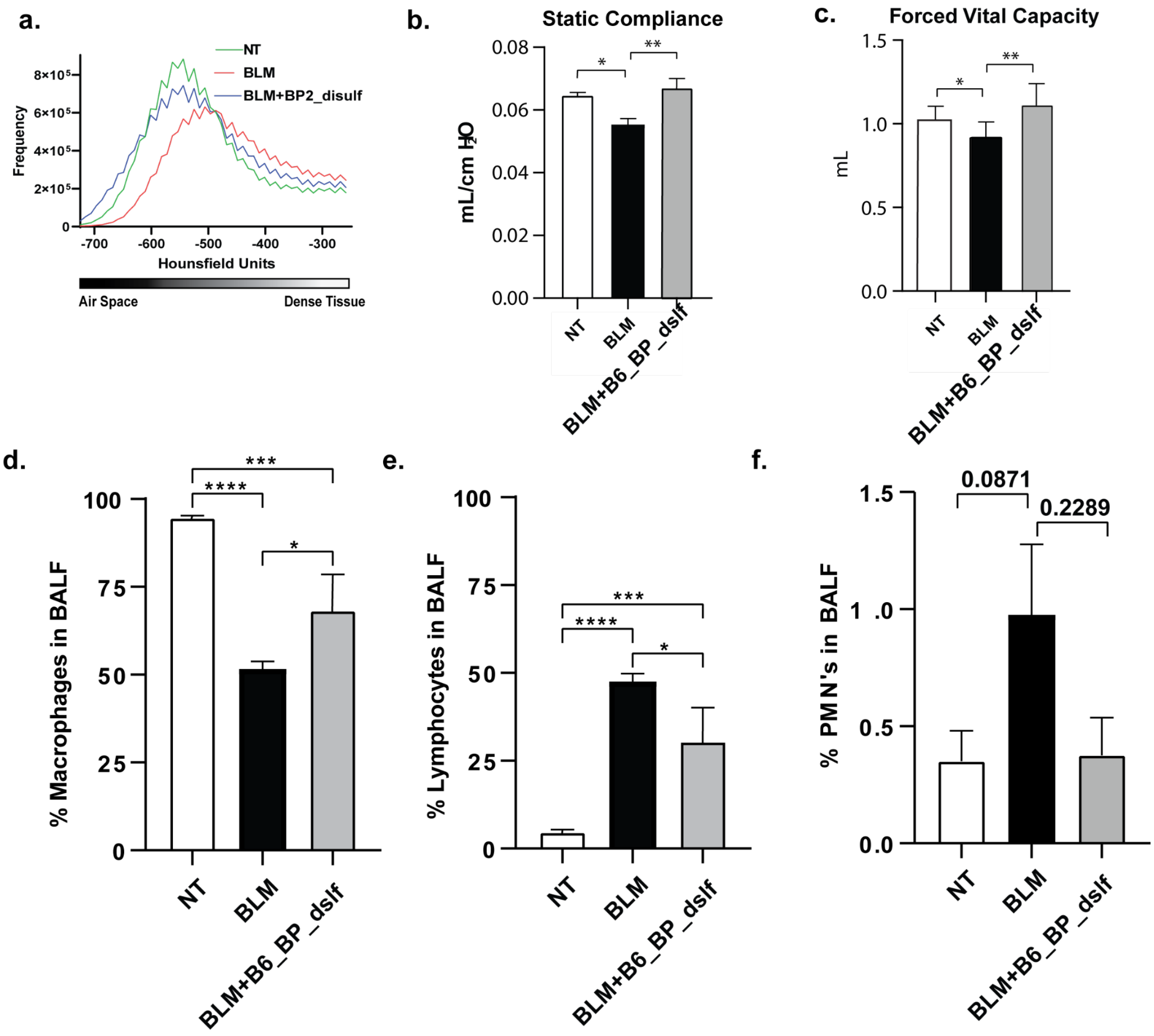

**Extended Data Fig. 14: Computed tomography (CT) imaging frequency of intensities, lung function, and cellular responses following B6_BP_dslf intraperitoneal (IP) administration in the "less severe" bleomycin-induced pulmonary fibrosis mouse model.** In the "less severe" bleomycin mouse model, lidocaine was not used to relax the vocal cords. a) Frequency of intensities of CT scans in Hounsfield Units for healthy non-treated (NT) mice, bleomycin-injured mice (BLM), and BLM+B6_BP_dslf treatment via intraperitoneal (IP) administration of 100 µg/kg (NT n=29; BLM n=14; BLM+B6_BP_dslf n=6). The distribution of the tissue density shows a shift to the right for bleomycin-injured mouse scan images indicating an increase in dense tissue compared to NT and B6_BP_dslf treated groups of mice; overall the

B6_BP_dslf treated and NT distributions are comparable. b, c) Measurement of lung mechanics on Day 21 for NT, BLM and BLM+B6_BP_dslf. Both static compliance (Cst, b) and forced vital capacity (FVC, c) improves and nears NT groups following B6_BP_dslf treatment as compared to the BLM only group. d,e,f) Differential counts of cellular response 21 days post bleomycin administration: the non-treated lungs contained predominantly macrophages (~94.77%, d), with relatively few lymphocytes (~4.77%, e) and polymorphonucleocytes (PMNs0.35%, f) present. Bleomycin treatment significantly increases inflammatory leukocytes in BALF, shifting the composition to 51.66% macrophages (d), 47.53% lymphocytes (e) and 0.97% PMNs (f). B6_BP_dslf treatment shifts this composition back to 68.4% macrophages (d), 30.5% lymphocytes (e), and 0.375% PMNs (f) nearing the BALF profile of the NT group. * $p < 0.05$. Error bars indicate standard error of the mean.

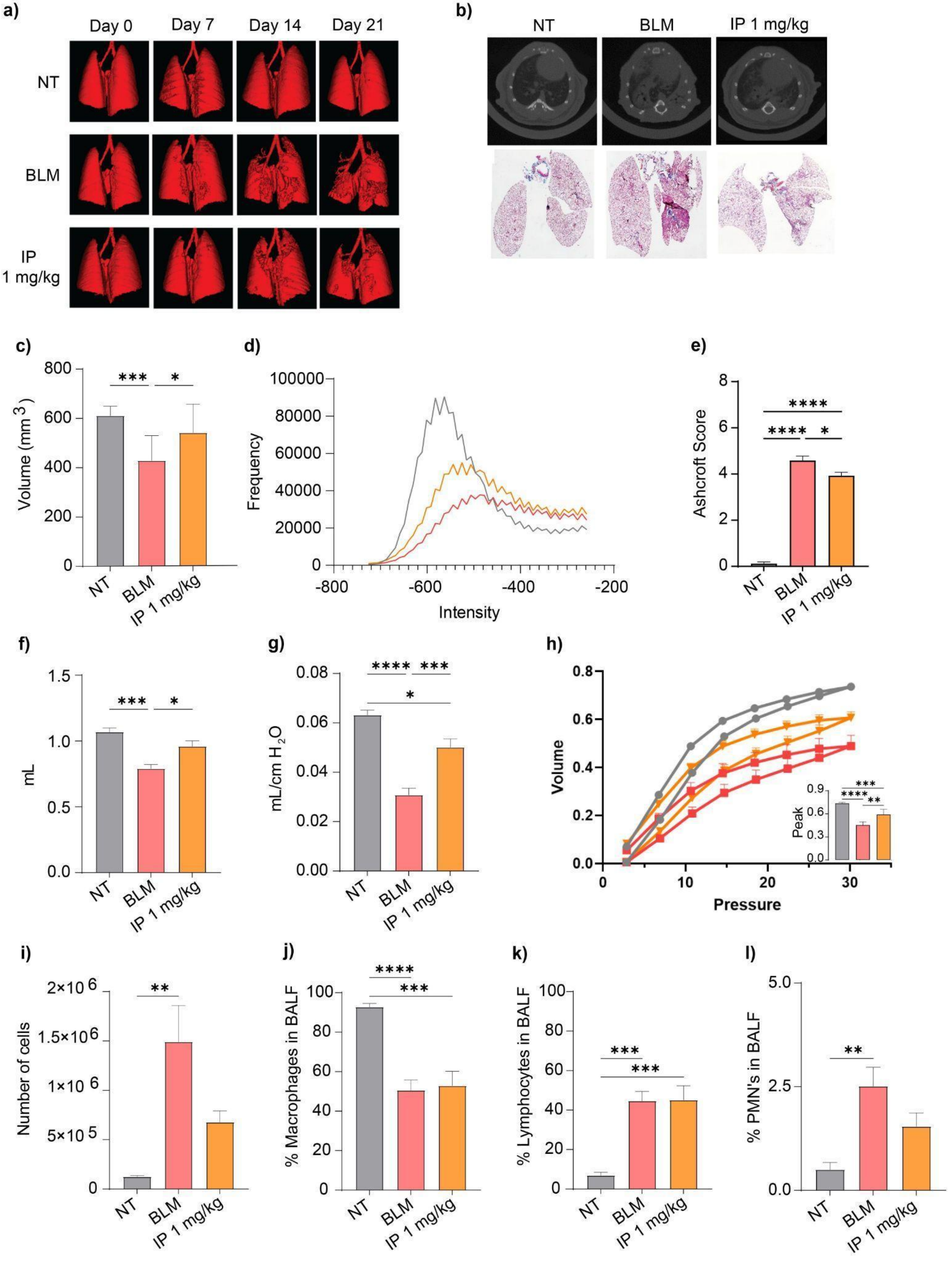
a)
Day 0
Day 7
Day 14
Day 21
NT
BLM
IP
1 mg/kg
b)
NT
BLM
IP 1 mg/kg
c)
Volume (mm 3 )
NT
BLM
IP 1 mg/kg
d)
Frequency
Intensity
e)
Ashcroft Score
NT
BLM
IP 1 mg/kg
f)
mL
NT
BLM
IP 1 mg/kg
g)
mL/cm H2O
NT
BLM
IP 1 mg/kg
h)
Volume
Pressure
Peak
i)
Number of cells
NT
BLM
IP 1 mg/kg
j)
% Macrophages in BALF
NT
BLM
IP 1 mg/kg
k)
% Lymphocytes in BALF
NT
BLM
IP 1 mg/kg
l)
% PMN's in BALF
NT
BLM
IP 1 mg/kg

**Extended Data Fig. 15: High-resolution micro-computed tomography (HR-µCT) imaging and lung function following B6_BP_dslf intraperitoneal (IP) administration in the "more severe" bleomycin-induced pulmonary fibrosis mouse model.** In the "more severe" bleomycin mouse model, lidocaine was used to relax the vocal cords and more fully insert the intratracheal tube and allow a more consistent and deeper pulmonary distribution of bleomycin (1 U/kg bleomycin intratracheally instilled at Day 0, every-other-day 1 mg/kg IP injections starting at Day 7). a) Longitudinal three-dimensional renderings of HR-µCT scans show the development of fibrosis. b) Upper panel: Representative HR-µCT scans at Day 21 with lower panel: representative whole lung stained with Masson Trichrome for NT, BLM, and B6_BP_dslf 1 mg/kg groups harvested at Day 21. c) Quantification of lung volumes from HR-µCT scans on Day 21 (mean ± SD, NT n=9, BLM n=10, B6_BP_dslf IP 1 mg/kg n=8). d) Histograms of the frequency of intensities from HR-µCT scans on Day 21 show an increase of healthy tissue and shift to intensities more like the healthy non-treated lungs following B6_BP_dslf injections as compared to BLM group, e) Average Ashcroft Score from Masson Trichrome sections is reduced after B6_BP_dslf treatment (mean ± SD, NT n=9, BLM n=3, B6_BP_dslf IP 1 mg/kg n =3). f) Forced Vital Capacity (mean ± SEM, NT n=9, BLM n=10, B6_BP_dslf IP 1 mg/kg n=9). g) Static Compliance (mean ± SEM, NT n=10, BLM n=12, B6_BP_dslf IP 1 mg/kg n=9). h) Pressure-Volume Loops measured by the SCIREQ flexiVent FX with peak volumes in the inset graph show a rescue from the restrictive nature of BLM-induced fibrosis. All lung function parameters were measured at Day 21 (mean ± SEM, NT n= , BLM n= , B6_BP_dslf IP 1 mg/kg n= ). i) Total BALF cell count at Day 21 (mean ± SD, NT n=8, BLM n=11, B6_BP_dslf IP 1 mg/kg n=7). Differential cell counts j) Macrophages (mean ± SD, NT n=6, BLM n=9, B6_BP_dslf IP 1 mg/kg n=7). k) Lymphocytes (mean ± SD, NT n=6, BLM n=9 B6_BP_dslf IP 1 mg/kg n=7). and l) PMNs (mean ± SD, NT n=6, BLM n=7, B6_BP_dslf IP 1 mg/kg n=6).

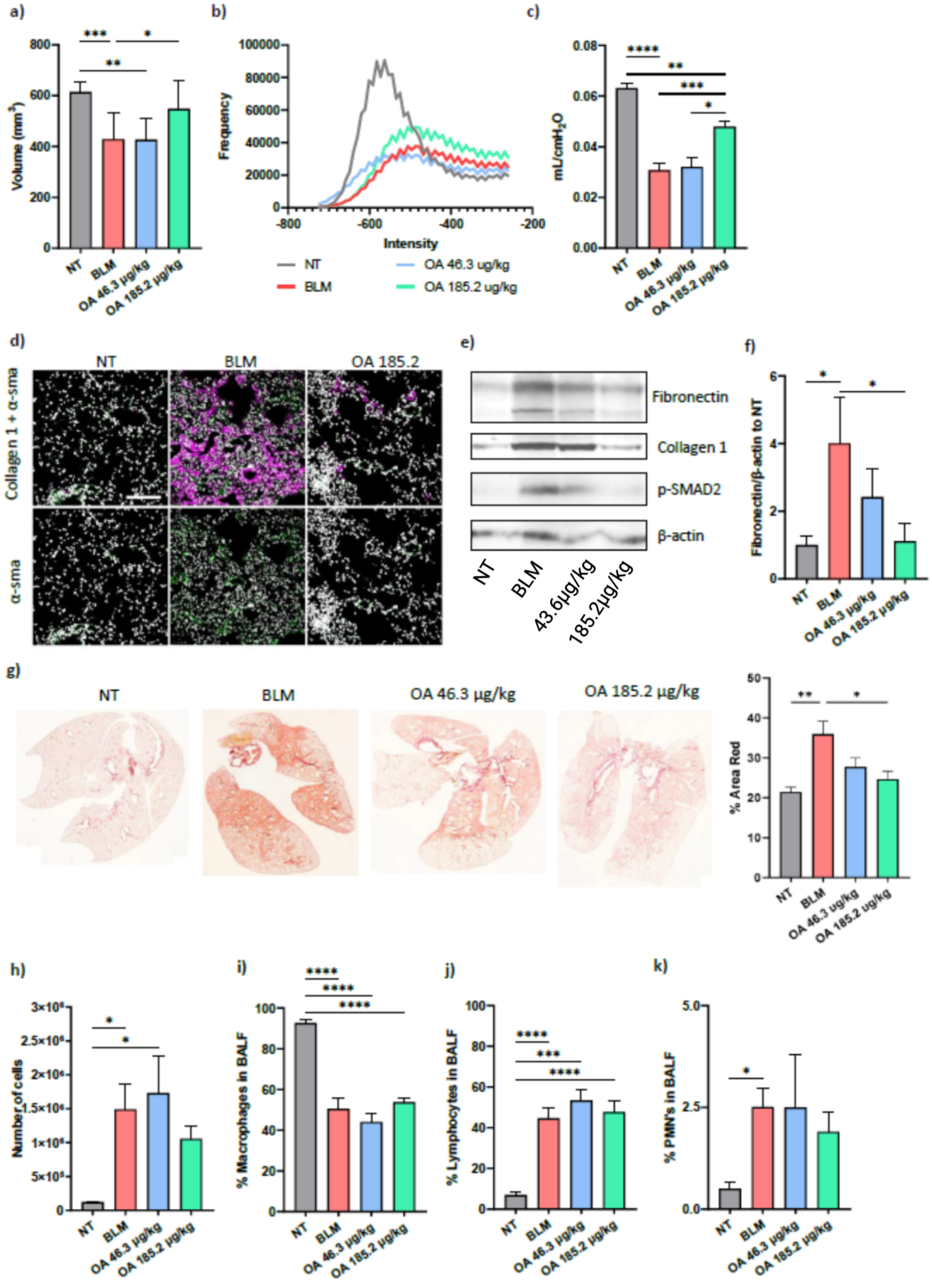

a)
Volume (mm³)
NT
BLM
OA 46.3 μg/kg
OA 185.2 μg/kg
b)
Frequency
Intensity
NT
BLM
OA 46.3 ug/kg
OA 185.2 ug/kg
c)
mL/cmH₂O
d)
NT
BLM
OA 185.2
Collagen 1 + α-sma
α-sma
e)
Fibronectin
Collagen 1
p-SMAD2
β-actin
NT
BLM
43.6μg/kg
185.2μg/kg
f)
Fibronectin/β-actin to NT
g)
NT
BLM
OA 46.3 μg/kg
OA 185.2 μg/kg
% Area Red
h)
Number of cells
i)
% Macrophages in BALF
j)
% Lymphocytes in BALF
k)
% PMN's in BALF

**Extended Data Fig. 16: Lung mechanics and cellular responses following B6_BP_dslf oropharyngeal administration (OA) in the "more severe" bleomycin-induced pulmonary fibrosis mouse model.** In the "more severe" bleomycin mouse model, lidocaine was used to relax the vocal cords and more fully insert the intratracheal tube and allow a more consistent and deeper pulmonary distribution of bleomycin (1 U/kg bleomycin intratracheally instilled at Day 0, every-other-day OA starting at Day 7). a) Quantification of HR-uCT volumes on Day 21 (mean ± SD, NT n=8, BLM n=10, B6_BP_dslf 46.3 ug/kg n=6, B6_BP_dslf 185.2 ug/kg n=9). b) Histograms of the frequency of intensities from HR-uCT scans on Day 21. c) Static Compliance as measured by the SCIREQ flexiVent FX at Day 21. (NT n=10, BLM n=12, B6_BP_dslf 46.3 ug/kg n=4, B6_BP_dslf 185.2 ug/kg n=6). d) Immunofluorescence histological staining of collagen type-I and α-SMA for NT, BLM and OA B6_BP_dslf 185.2 ug/kg group show a reduction in these two fibrotic markers with the B6_BP_dslf treated group. e) Representative mages from western blot analysis for pro-fibrotic markers: Fibronectin, Collagen 1, p-SMAD2 with B-actin loading control. Quantification of Collagen 1 and p-SMAD are represented in Fig 4e, f. f) Quantification of Fibronectin western blot (mean ± SD, NT n=3, BLM n=3, B6_BP_dslf 46.3 ug/kg n=3, B6_BP_dslf 185.2 ug/kg n=3). g) Representative Sirius Red images and quantification of percentage red area. h) Total BALF Cell Counts from recovered BALF fluid on Day 21 (mean ± SD, NT n=8, BLM n=11, B6_BP_dslf 46.3 ug/kg n=3, B6_BP_dslf 185.2 ug/kg n=7). Differential cell counts i) Macrophage (mean ± SD, NT n=6, BLM n=9, B6_BP_dslf 46.3 ug/kg n=3, B6_BP_dslf 185.2 ug/kg n=6). j) Lymphocyte (mean ± SD, NT n=6, BLM n=9, B6_BP_dslf 46.3 ug/kg n=3, B6_BP_dslf 185.2 ug/kg n=7), and k) PMNs (mean ± SD, NT n=6, BLM n=7, B6_BP_dslf 46.3 ug/kg n=3, B6_BP_dslf 185.2 ug/kg n=6).

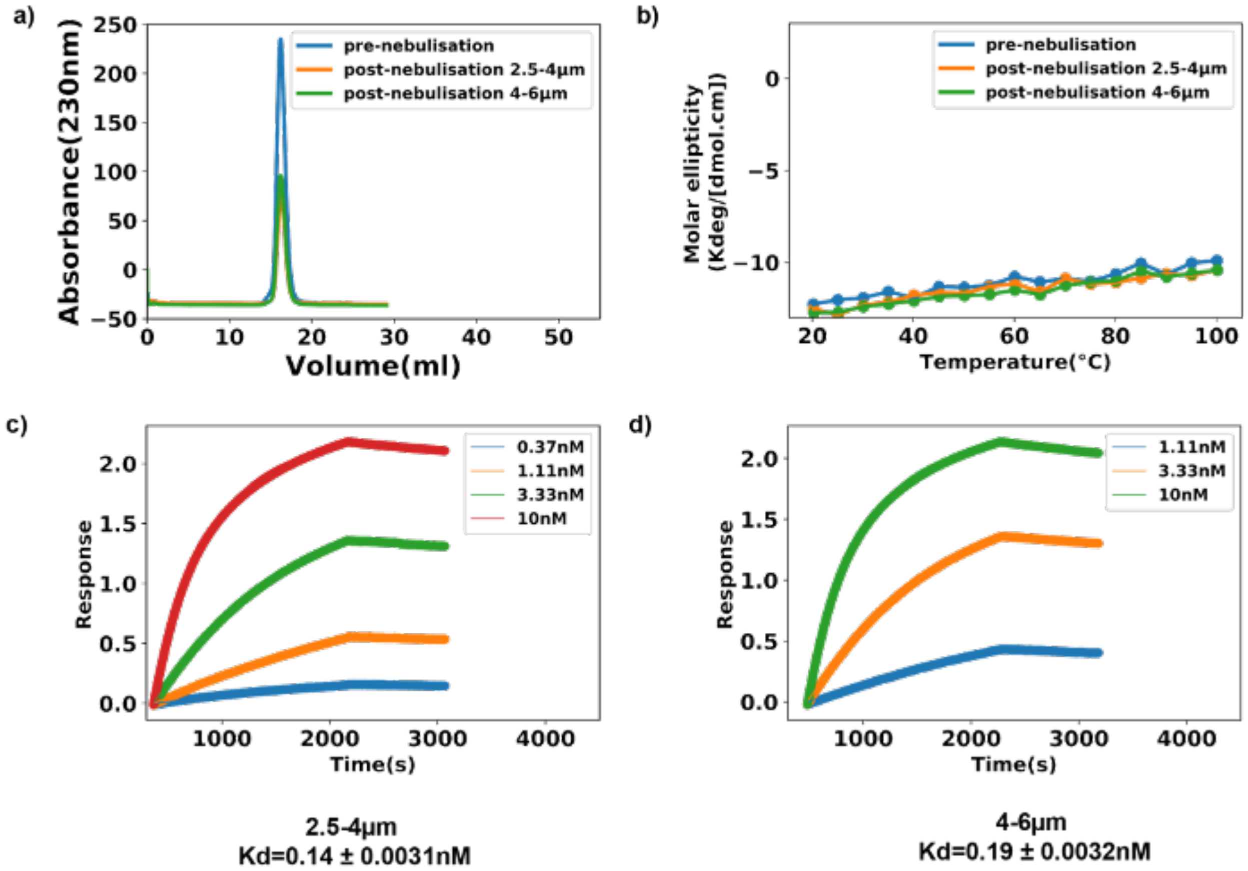

**Extended Data Fig. 17: Stability of B6_BP_dslf following nebulization.** a) Size-exclusion chromatography pre- and post-nebulization. b) Thermal melt of B6_BP_dslf pre- and post-nebulization using circular dichroism spectroscopy. The protein remains highly thermostable following nebulization. c,d) BLI binding experiment using aerosolized B6_BP_dslf and human αvβ6 integrin. B6_BP_dslf binds to αvβ6 with similar affinity post-nebulization for both particle sizes.

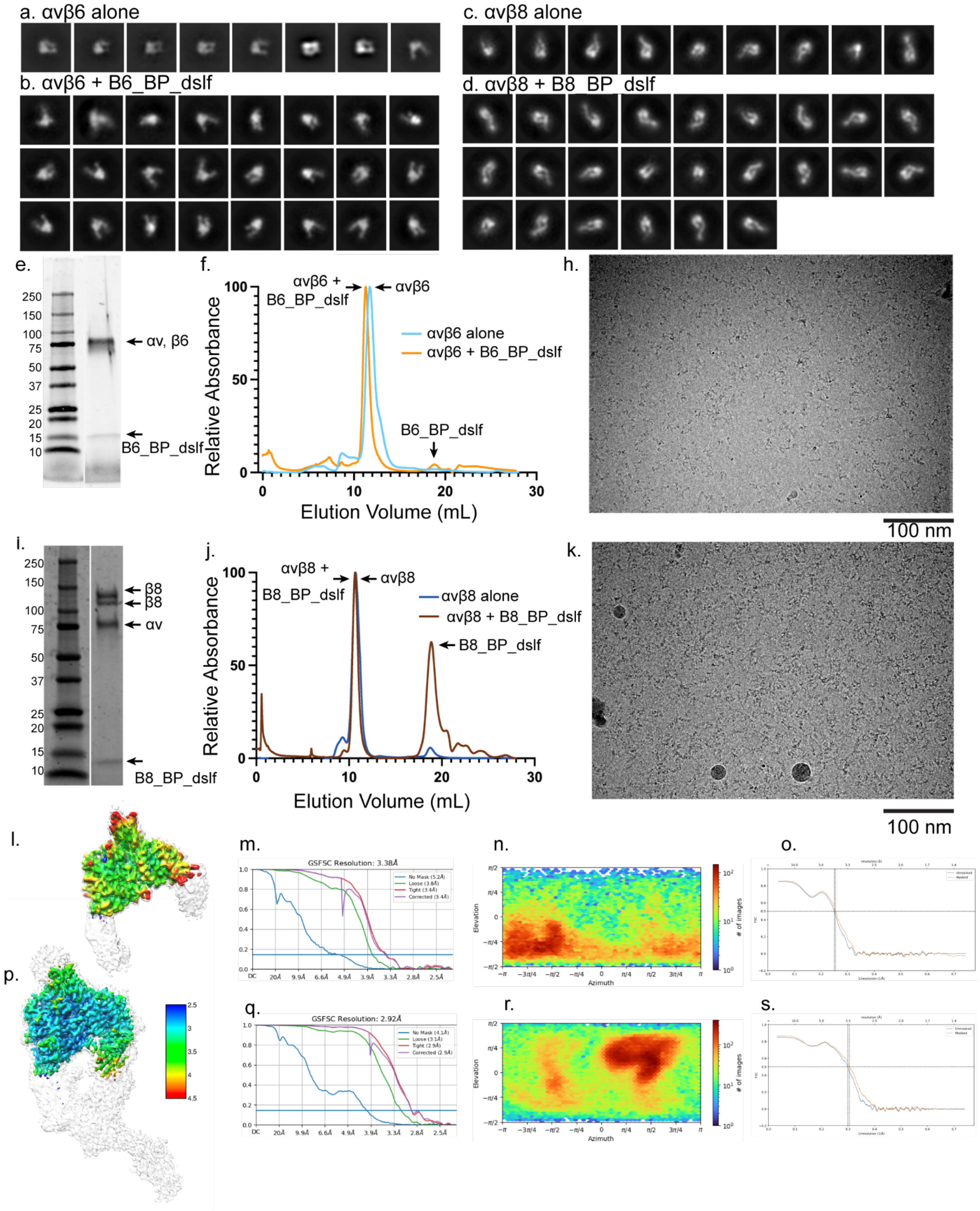

**Extended Data Fig. 18: Negative stain EM class averages, SDS-PAGE, SEC, and CryoEM micrographs and statistics of αvβ6 and αvβ8 integrin alone and in complex with respective selective minibinders B6_BP_dslf and B8_BP_dslf. a-d)** Negative stain EM class averages of (a) αvβ6 and (c) αvβ8 alone, and complexes of (b) αvβ6 + B6_BP_dslf and (d)

αvβ8 + B8_BP_dslf. **e, i)** SDS-PAGE of complexes of (e) αvβ6 + B6_BP_dslf and (i) αvβ8 + B8_BP_dslf. **f, j)** Size-exclusion chromatography of (f) αvβ6 alone and in complex with B6_BP_dslf and (j) αvβ8 alone and in complex with B8_BP_dslf using a Superdex 200 Increase 10/300 SEC column (Cytiva). **h, k)** Representative motion corrected micrographs of (h) αvβ6 + B6_BP_dslf and (k) αvβ8 + B8_BP_dslf particles suspended in vitreous ice.

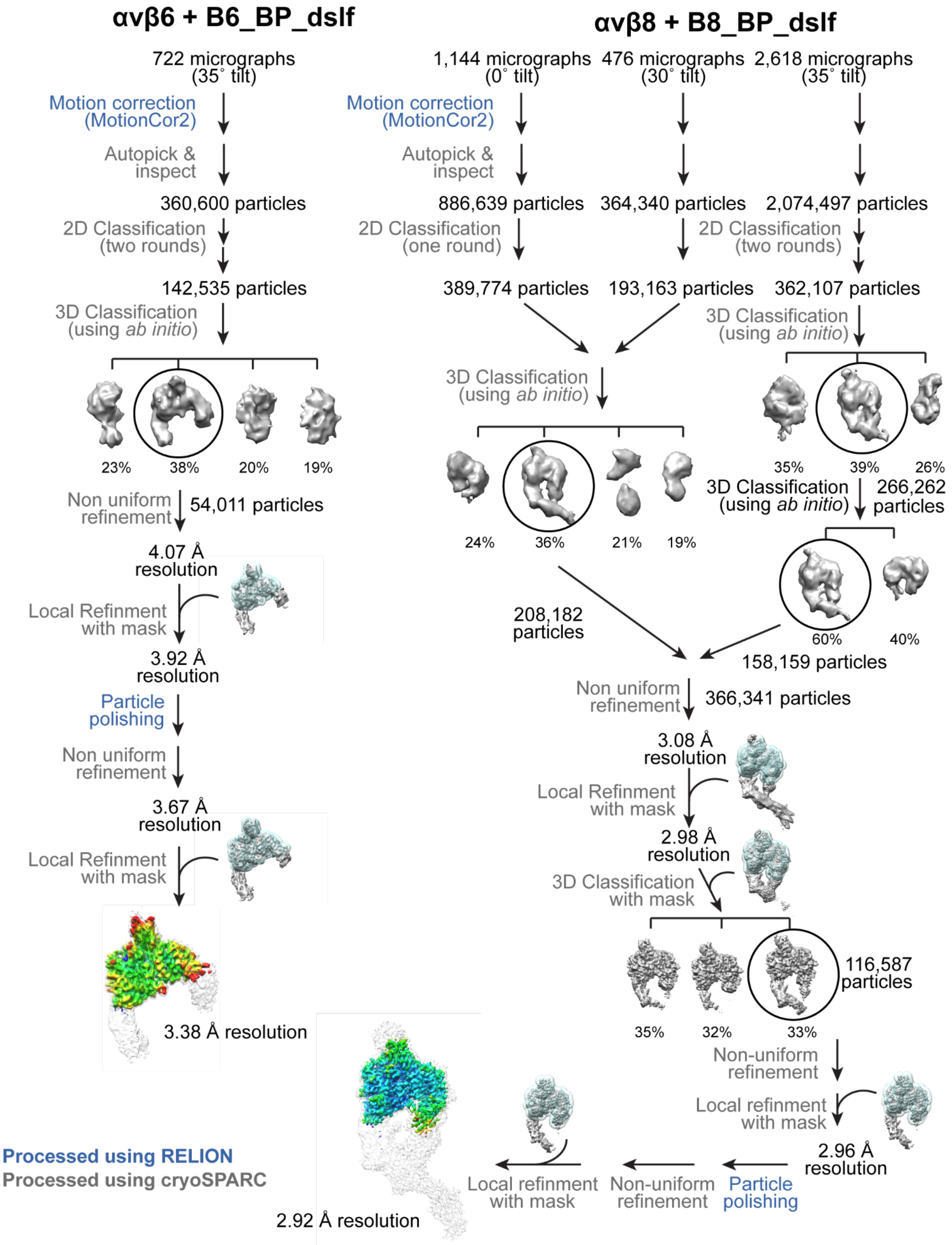

**Extended Data Fig. 19: Processing schematic for cryoEM of αvβ6 + B6_BP_dslf and αvβ8 + B8_BP_dslf complexes.** A schematic flowchart showing the classification scheme of the αvβ6 + B6_BP_dslf and αvβ8 + B8_BP_dslf complexes. Particle numbers at each step and for each class are indicated. All maps are colored on the same scale, as indicated, based on local resolution estimates.

**Extended Data Tables**

**Extended Data Table 1: Kinetic analysis of BLI binding of purified mutants against titrations of biotinylated human αvβ6 or αvβ8.**

| Mutant | Receptor | $K_{on}$ ($M^{-1}s^{-1}$) | $K_{dis}$ ($s^{-1}$) | $K_D$ (nM) |
|---|---|---|---|---|
| **E13T (B6B8_BP)** | h-αvβ6 | $1.68x10^5 \pm 1.60x10^2$ | $5.60x10^{-5} \pm 2.26x10^{-7}$ | 0.33 ± 0.0014 |
| **M15R** | h-αvβ6 | $1.75x10^5 \pm 1.26x10^2$ | $6.65x10^{-5} \pm 1.74x10^{-7}$ | 0.38 ± 0.0010 |
| **A39K** | h-αvβ6 | $1.27x10^5 \pm 2.91x10^1$ | $8.16x10^{-5} \pm 1.23x10^{-7}$ | 0.64 ± 0.0010 |
| **G64R** | h-αvβ6 | $1.41x10^5 \pm 1.89x10^2$ | $1.34x10^{-4} \pm 3.08x10^{-7}$ | 0.95 ± 0.0025 |
| **E13TG64R** | h-αvβ6 | $1.12x10^5 \pm 8.77x10^1$ | $8.87x10^{-5} \pm 1.60x10^{-7}$ | 0.79 ± 0.0016 |
| **M15RG64R** | h-αvβ6 | $1.04x10^5 \pm 1.72x10^2$ | $7.71x10^{-5} \pm 3.19x10^{-7}$ | 0.74 ± 0.0033 |
| **A39KG64R (B6_BP)** | h-αvβ6 | $9.92x10^4 \pm 9.83x10^1$ | $8.82x10^{-5} \pm 1.85x10^{-7}$ | 0.89 ± 0.0021 |
| **E13TM15RA39K** | h-αvβ6 | $8.43x10^4 \pm 1.32x10^2$ | $9.10x10^{-5} \pm 2.62x10^{-7}$ | 1.08 ± 0.0035 |
| **E13TM15RA39KG64R** | h-αvβ6 | $1.04x10^5 \pm 8.88x10^1$ | $8.31x10^{-5} \pm 1.63x10^{-7}$ | 0.79 ± 0.0017 |
| **av6_3** | h-αvβ6 | $8.16x10^4 \pm 1.37x10^2$ | $9.67x10^{-5} \pm 4.43x10^{-7}$ | 1.19 ± 0.0058 |
| **B6_BP_dslf** | h-αvβ6 | $7.44x10^4 \pm 5.5x10^1$ | $3.08x10^{-5} \pm 1.00x10^{-7}$ | 0.50 ± 0.0010 |
| **B8_BP_dslf** | h-αvβ8 | $8.03x10^4 \pm 1.70x10^2$ | $2.39x10^{-4} \pm 7.83x10^{-7}$ | 1.97 ± 0.0012 |
| **avb8_12** | h-αvβ8 | $1.19x10^5 \pm 1.22x10^3$ | $3.97x10^{-5} \pm 1.00x10^{-7}$ | 0.33 ± 0.0010 |
| **avb8_12** | h-αvβ6 | $3.47x10^5 \pm 1.07x10^3$ | $3.92x10^{-4} \pm 5.33x10^{-7}$ | 1.13 ± 0.0038 |

**Extended Data Table 2: Affinity and selectivity comparison of B6_BP against other leading reported αvβ6 inhibitors.**

| Integrin | B6_BP Affinity (nM)[1] | Fold Selectivity | GSK3008348 (IC50 nM)[2] | Fold Selectivity | A20FMDV2 (nM)[3] | Fold selectivity | Knottin R01 (nM)[4,*] | Fold selectivity | 264-RAD (nM) | Fold selectivity |
|---|---|---|---|---|---|---|---|---|---|---|
| αvβ6 | 0.11 | | 0.012 | | 3 | | 3.6 | | .01 | |
| αvβ1 | >200 | >1818 | 2.6 | 190 | Not reported | N/A | Not reported | Not reported | Not reported | Not reported |
| αvβ3 | >200 | >1818 | 37.4 | 3375 | >10000 | >3333 | >10 | >2.77 | Does not bind | N/A |
| αvβ5 | >200 | >1818 | 11.7 | 1035 | >100000 | >33330 | >10 | >2.77 | Does not bind | N/A |
| αvβ8 | 580 | >5000 | 2.1 | 182 | Not reported | Not reported | Not reported | Not reported | Binds (Kd not reported) | |
| α5β1 | >200 | >1818 | 97.2 | 6911 | >10000 | >33330 | >10 | >2.77 | Does not bind | N/A |
| α8β1 | >200 | >1818 | 0.3 | 26 | Not reported | Not reported | Not reported | Not reported | Not reported | Not reported |
| αiibβ3 | >200 | >1818 | 10000 | 917814 | Not reported | Not reported | Not reported | Not reported | Not reported | Not reported |

1: Measured using stable K562 cell lines overexpressing corresponding integrins.

2: Measured by competition assay using radio-labelled ligands.

3: Measured by competitive ELISA

4: Measured by integrin protein capture on immobilized knottins

*: This variant has been subjected to directed evolution in subsequent publications.

**Extended Data Table 3. Predicted and observed interactions between integrins αvβ6/αvβ8 and designed minibinders.**

| **αvβ6 - B6_BP_dslf *predicted* interactions by design** (confirmed interactions shown in **bold**) | | | | **αvβ6 - B6_BP_dslf *observed* interactions in cryoEM** (predicted interactions shown in **bold**) | | | |
|---|---|---|---|---|---|---|---|
| **αv** | **β6** | **minibinder** | **interaction type** | **αv** | **β8** | **minibinder** | **interaction type** |
| 150 | | 10 | hydrogen | 213 | | 10 | hydrogen |
| 177 | | 10 | hydrogen | **218** | | **10** | **hydrogen** |
| 178 | | 10 | hydrogen | **218** | | **10** | **salt** |
| **218** | | **10** | **hydrogen** | | **126** | **12** | **hydrogen** |
| 150 | | 10 | salt | | 127 | 12 | hydrogen |
| **218** | | **10** | **salt** | | **218** | **12** | **hydrogen** |
| | **126** | **12** | **hydrogen** | | **219** | **12** | **hydrogen** |

| | | | | | | | |
|---|---|---|---|---|---|---|---|
| | **219** | **12** | **hydrogen** | | **129** | **19** | **hydrogen** |
| | **218** | **12** | **hydrogen** | | **316** | **41** | **hydrogen** |
| | **129** | **19** | **hydrogen** | | **182** | **63** | **hydrogen** |
| | 185 | 19 | hydrogen | | **183** | **63** | **hydrogen** |
| | 254 | 39 | hydrogen | | **129** | **19** | **salt** |
| | **316** | **41** | **hydrogen** | | **316** | **41** | **salt** |
| | **182** | **63** | **hydrogen** | | | | |
| | **183** | **63** | **hydrogen** | | | | |
| | **129** | **19** | **salt** | | | | |
| | **316** | **41** | **salt** | | | | |

| **αvβ8 - B8_BP_dslf *predicted* interactions by design (confirmed interactions shown in bold)** | | | | **αvβ8 - B8_BP_dslf *observed* interactions in (predicted interactions shown in bold)** | | | |
|---|---|---|---|---|---|---|---|
| **αv** | **β6** | **minibinder** | **interaction type** | **αv** | **β8** | **minibinder** | **interaction type** |
| 150 | | 10 | hydrogen | **218** | | **10** | **hydrogen** |
| **218** | | **10** | **hydrogen** | **215** | | **41** | **hydrogen** |
| **215** | | **41** | **hydrogen** | 119 | | 66 | hydrogen |
| 119 | | 63 | hydrogen | 178 | | 67 | hydrogen |
| 150 | | 10 | salt | **218** | | **10** | **salt** |
| **218** | | **10** | **salt** | | 114 | 12 | hydrogen |
| | 98 | 12 | hydrogen | | 115 | 12 | hydrogen |
| | 99 | 12 | hydrogen | | 116 | 12 | hydrogen |
| | 189 | 12 | hydrogen | | 207 | 12 | hydrogen |
| | 285 | 40 | hydrogen | | 208 | 12 | hydrogen |
| | 285 | 40 | salt | | 115 | 16 | hydrogen |
| | | | | | 304 | 40 | hydrogen |
| | | | | | 304 | 40 | salt |

**Supplementary Table 1. Statistics of X-ray diffraction and structure refinement.** [1]

**Data collection and refinement statistics (molecular replacement)**

| | **BP (PDB: 7LMV)** | **B6B8_BP_dslf (PDB: 7LMX)** |
|---|---|---|
| **Data collection** | | |
| Space group | *$P3_1$* | *$P2_12_12_1$* |
| Cell dimensions | | |
| *a*, *b*, *c* (Å) | 92.27, 92.27, 82.42 | 40.69, 64.1, 80.99 |
| α, β, γ (°) | 90, 90, 120 | 90, 90, 90 |
| Resolution (Å) | 40.26 - 1.90 (1.96 -1.90)* | 50.00 - 1.80 (1.86 - 1.80)* |
| $R_{merge}$ | 0.093 (0.613) | 0.087 (1.283) |
| $I / \sigma I$ | 10.4 (2.4) | 13.5 (2.5) |
| Completeness (%) | 99.9 (99.8) | 98.9 (100.0) |
| Redundancy | 5.6 (4.5) | 6.0 (6.3) |
| **Refinement** | | |
| Resolution (Å) | 40.26 - 1.90 (1.96 - 1.90) | 26.20 - 1.80 (1.86 - 1.80) |
| No. reflections | 61653 (6206) | 20060 (1978) |
| $R_{work}$ / $R_{free}$ | | |
| No. atoms | | |

| | | |
|---|---|---|
| Protein | 6444 | 1777 |
| Ligand/ion | 0 | 0 |
| Water | 334 | 91 |
| *B*-factors | | |
| Protein | 36.56 | 42.99 |
| Ligand/ion | 0 | 0 |
| Water | 34.80 | 44.23 |
| R.m.s. deviations | | |
| Bond lengths (Å) | 0.006 | 0.004 |
| Bond angles (°) | 0.560 | 0.606 |

*Single Crystal used for each data collection. *Values in parentheses are for highest-resolution shell.

**Data collection, phasing and refinement statistics (MIR)**

| | Crystal 1 name | Crystal 2 name |
|---|---|---|
| **Data collection** | | |
| Space group | | |
| Cell dimensions | | |
| *a*, *b*, *c* (Å) | | |
| $\alpha$, $\beta$, $\gamma$ (°) | | |
| Resolution (Å) | ##(high res shell) * | |
| $R_{sym}$ or $R_{merge}$ | ##(high res shell) | |
| $I / \sigma I$ | ##(high res shell) | |
| Completeness (%) | ##(high res shell) | |
| Redundancy | ##(high res shell) | |
| **Refinement** | | |
| Resolution (Å) | | |
| No. reflections | | |
| $R_{work}$ / $R_{free}$ | | |
| No. atoms | | |
| Protein | | |
| Ligand/ion | | |
| Water | | |
| *B*-factors | | |
| Protein | | |
| Ligand/ion | | |
| Water | | |
| R.m.s deviations | | |
| Bond lengths (Å) | | |
| Bond angles (°) | | |

*Number of xtals for each structure should be noted in footnote. *Values in parentheses are for highest-resolution shell.

[AU: Equations defining various *R*-values are standard and hence are no longer defined in the footnotes.] [AU: Phasing data should be reported in Methods section.]
[AU: Ramachandran statistics should be in Methods section at end of Refinement subsection.]
[AU: Wavelength of data collection, temperature and beamline should all be in Methods section.]

**Data collection, phasing and refinement statistics for MAD (SeMet) structures**

| | Native | | Crystal 1 name | | | Crystal 2 name | |
|---|---|---|---|---|---|---|---|
| **Data collection** | | | | | | | |
| Space group | | | common # | | | common # | |
| Cell dimensions | | | | | | | |
| *a*, *b*, *c* (Å) | | | common # | | | common # | |
| α, β, γ (°) | | | common # | | | common # | |
| | | *Peak* | *Inflection* | *Remote* | *Peak* | *Inflection* | *Remote* |
| Wavelength | | # | # | # | # | # | # |
| Resolution (Å) | | # | # | # | # | # | # |
| $R_{sym}$ or $R_{merge}$ | | # | # | # | # | # | # |
| $I / \sigma I$ | | # | # | # | # | # | # |
| Completeness (%) | | # | # | # | # | # | # |
| Redundancy | | # | # | # | # | # | # |
| **Refinement** | | | | | | | |
| Resolution (Å) | | | common # | | | common # | |
| No. reflections | | | | | | | |
| $R_{work}$ / $R_{free}$ | | | | | | | |
| No. atoms | | | | | | | |
| Protein | | | | | | | |
| Ligand/ion | | | | | | | |
| Water | | | | | | | |
| *B*-factors | | | | | | | |
| Protein | | | | | | | |
| Ligand/ion | | | | | | | |
| Water | | | | | | | |
| R.m.s deviations | | | | | | | |
| Bond lengths (Å) | | | | | | | |
| Bond angles (°) | | | | | | | |

*Number of xtals for each structure should be noted in footnote. *Values in parentheses are for highest-resolution shell.

[AU: Equations defining various *R*-values are standard and hence are no longer defined in the footnotes.]
[AU: Phasing data should be reported in Methods section.]
[AU: Ramachandran statistics should be in Methods section at end of Refinement subsection.]
[AU: Wavelength of data collection, temperature and beamline should all be in Methods section.]

**Supplementary Table 2. Exclusion criteria for bleomycin-induced pulmonary fibrosis.**

| **Treatment Group** (Extended Data Figs. 13, 14) | **Exclusion Criteria A[1]** | **Exclusion Criteria B** | **Exclusion Criteria C** | **Included in study** |
|---|---|---|---|---|

| | | | | |
|---|---|---|---|---|
| NT | 0 | 0 | 0 | 29 |
| BLM | 20 | 0 | 0 | 14 |
| BLM + IP | 18 | 0 | 0 | 6 |
| **Treatment Group** (Figure 4, Extended Data Figs. 15, 16) | | | | |
| BLM | 1 | 5 | 8 | 10 |
| BLM + IP | 0 | 5 | 10 | 9 |
| Inhalations[2] | 0 | 16 | 12 | 9 |

1. These criteria are as follows: Exclusion Criteria A: no development of fibrosis, Exclusion Criteria B: culled because of morbid weight loss, and Exclusion Criteria C: unexpected deaths from experimental procedures
2. n=4 43.6 µg/kg, n=5 185.2 µg/kg

**Supplementary Table 3: Sequences of the all designs and evolved variants reported in this paper**

| Construct | Sequence |
|---|---|
| Design 2 | AEVRFVFRGDLTELMLRAVKDHLKKEGPHWNITSRGNELEVRGSHESDAKRIQKEFPSVQSTTQA |
| BP | AVVRFVFRGDLAELMLRAVKDHLKKEGPHWNITSRGNELVVRGIHESDAKRIQKEFPSVQSTIQA |
| av6_1 | AVVRFVFRGDLAELMLRAVKDHLKKEGPHWNITSTNNGAELVVRGIHESDAKRIARTVEKLTNGKSQSLVLT |
| av6_2 | AVVRFVFRGDLAELMLRAVKDHLKKEGPHWNITSTNNGAELVVRGIHESDAKRIANWAKTYSPGGKESYTIP |
| av6_3 | AVVRFVFRGDLAELMLRAVKDHLKKEGPHWNITSTNNGAELVVRGIHESDAKRIAKWVEKRFPGVHTETQQD |
| av6_4 | AVVRFVFRGDLAELMLRAVKDHLKKEGPHWNITSTNNGAELVVRGIHESDAKRIAKWARLKFPGTDTRIEVR |
| av6_5 | AVVRFVFRGDLAELMLRAVKDHLKKEGPHWNITSDESGFELVVRGIHESDAKRIARTVEKLTNGKSQSLVLT |
| av6_6 | AVVRFVFRGDLAELMLRAVKDHLKKEGPHWNITSDESGFELVVRGIHESDAKRIANWAKTYSPGGKESYTIP |

| | |
|---|---|
| av6_7 | AVVRFVFRGDLAELMLRAVKDHLKKEGPHWNITSDESGFELVVRGIHESDAKRIAKWVEKRFP GVHTETQQD |
| av6_8 | AVVRFVFRGDLAELMLRAVKDHLKKEGPHWNITSDESGFELVVRGIHESDAKRIAKWARLKFP GTDTRIEVR |
| av6p_9 | AVVRFVFRGDLAELMLRAVKDHLKKEGPHWNITSDTSKGAELVVRGIHESDAKRIARTVEKLT NGKSQSLVL |
| av6_10 | AVVRFVFRGDLAELMLRAVKDHLKKEGPHWNITSDTSKGAELVVRGIHESDAKRIANWAKTYS PGGKESYTI |
| av6_11 | AVVRFVFRGDLAELMLRAVKDHLKKEGPHWNITSDTSKGAELVVRGIHESDAKRIAKWVEKRF PGVHTETQQ |
| av6_12 | AVVRFVFRGDLAELMLRAVKDHLKKEGPHWNITSDTSKGAELVVRGIHESDAKRIAKWARLKF PGTDTRIEV |
| av6_13 | AVVRFVFRGDLAELMLRAVKDHLKKEGPHWNITSVESSQVELVVRGIHESDAKRIARTVEKLT NGKSQSLVL |
| av6_14 | AVVRFVFRGDLAELMLRAVKDHLKKEGPHWNITSVESSQVELVVRGIHESDAKRIANWAKTYS PGGKESYTI |
| av6_15 | AVVRFVFRGDLAELMLRAVKDHLKKEGPHWNITSVESSQVELVVRGIHESDAKRIAKWVEKRF PGVHTETQQ |
| av6_16 | AVVRFVFRGDLAELMLRAVKDHLKKEGPHWNITSVESSQVELVVRGIHESDAKRIAKWARLKF PGTDTRIEV |
| E13T(B6B 8_BP) | AVVRFVFRGDLATLMLRAVKDHLKKEGPHWNITSTNNGAELVVRGIHESDAKRIAKWVEKRFP GVHTETQQD |
| M15R | AVVRFVFRGDLAELRLRAVKDHLKKEGPHWNITSTNNGAELVVRGIHESDAKRIAKWVEKRFP GVHTETQQD |
| A39K | AVVRFVFRGDLAELMLRAVKDHLKKEGPHWNITSTNNGKELVVRGIHESDAKRIAKWVEKRFP GVHTETQQD |
| P63R | AVVRFVFRGDLAELMLRAVKDHLKKEGPHWNITSTNNGAELVVRGIHESDAKRIAKWVEKRF RGVHTETQQD |
| G64R | AVVRFVFRGDLAELMLRAVKDHLKKEGPHWNITSTNNGAELVVRGIHESDAKRIAKWVEKRFP RVHTETQQD |
| M15RG64 R | AVVRFVFRGDLAELRLRAVKDHLKKEGPHWNITSTNNGAELVVRGIHESDAKRIAKWVEKRFP RVHTETQQD |
| A39KG64 R (B6_BP) | AVVRFVFRGDLAELMLRAVKDHLKKEGPHWNITSTNNGKELVVRGIHESDAKRIAKWVEKRFP RVHTETQQD |
| E13TG64 R | AVVRFVFRGDLATLMLRAVKDHLKKEGPHWNITSTNNGAELVVRGIHESDAKRIAKWVEKRFP RVHTETQQD |

| | |
|---|---|
| E13TM15 RA39K | AVVRFVFRGDLATLRLRAVKDHLKKEGPHWNITSTNNGKELVVRGIHESDAKRIAKWVEKRFPGVHTETQQD |
| E13TM15 RA39KG64R | AVVRFVFRGDLATLRLRAVKDHLKKEGPHWNITSTNNGKELVVRGIHESDAKRIAKWVEKRFPRVHTETQQD |
| E13T_disulf(B6B8_BP_dslf) | TKCVVRFVFRGDLATLMLRAVKDHLKKEGPHWNITSTNNGAELVVRGIHESDAKRIAKWVEKRFPGVHTETQCD |
| A39KG64R_disulf (B6_BP_dslf) | TKCVVRFVFRGDLAELMLRAVKDHLKKEGPHWNITSTNNGKELVVRGIHESDAKRIAKWVEKRFPRVHTETQCD |
| avb8_1 | TKCVVRFEFRGDFARIMLKAVKDHLKKEGPHWNITEVNTQWPQLVVRGIHESDAKRIAKWVESRIPYIRVETQCD |
| avb8_2 | TKCVVRFIFRGDFAWVALRAVKDHLKKEGPHWNITETNQDYMQLVVRGIHESDAKRIAKWVENRFPYARVETQCD |
| Avb8_3 (B8_BP_dslf) | TKCVVRFNFRGDMAVYALKAVKDHLKKEGPHWNITTHNDDEVYLVVRGIHESDAKRIAKWVESTIPGISVETQCD |
| avb8_4 | TKCVVRFIFRGDFAKYMLRAVKDHLKKEGPHWNITETHTQYTQLVVRGIHESDAKRIAKWVQSRFPWIKVETQCD |
| avb8_5 | TKCVVRFIFRGDMANYALKAVKDHLKKEGPHWNITTQEEPNQYLVVRGIHESDAKRIAKWVEKRFPFAGVETQCD |
| avb8_6 | TKCVVRFVFRGDFAKYMLKAVKDHLKKEGPHWNITEGESEENQLVVRGIHESDAKRIAKWVQTRFPYIRVETQCD |
| avb8_7 | TKCVVRFEFRGDLAWLALKAVKDHLKKEGPHWNITDVNTDLIQLVVRGIHESDAKRIAKWVESTIPWVRVETQCD |
| avb8_8 | TKCVVRFLFRGDLAVLALRAVKDHLKKEGPHWNITDANDESYSLVVRGIHESDAKRIAKWVQSRIPYVRAETQCD |
| avb8_9 | TKCVVRFIFRGDMAQYALKAVKDHLKKEGPHWNITTQEEPNQYLVVRGIHESDAKRIAKWVEKRFPFAGVETQCD |
| avb8_10 | TKCVVRFEFRGDLATIMLKAVKDHLKKEGPHWNITEVNTQWPQLVVRGIHESDAKRIAKWVESRIPYIRVETQCD |

| | |
|---|---|
| avb8_11 | TKCVVRFIFRGDLATIALRAVKDHLKKEGPHWNITETNQDYMQLVVRGIHESDAKRIAKWVEN RFPYARVETQCD |
| avb8_12 | TKCVVRFNFRGDLATIALKAVKDHLKKEGPHWNITTHNDDEVYLVVRGIHESDAKRIAKWVES TIPGISVETQCD |
| avb8_13 | TKCVVRFIFRGDLATIMLRAVKDHLKKEGPHWNITETHTQYTQLVVRGIHESDAKRIAKWVQS RFPWIKVETQCD |
| avb8_14 | TKCVVRFIFRGDLATIALKAVKDHLKKEGPHWNITTQEEPNQYLVVRGIHESDAKRIAKWVEKR FPFAGVETQCD |
| avb8_15 | TKCVVRFVFRGDLATIMLKAVKDHLKKEGPHWNITEGESEENQLVVRGIHESDAKRIAKWVQT RFPYIRVETQCD |
| avb8_16 | TKCVVRFEFRGDLATIALKAVKDHLKKEGPHWNITDVNTDLIQLVVRGIHESDAKRIAKWVEST IPWVRVETQCD |
| avb8_17 | TKCVVRFLFRGDLATIALRAVKDHLKKEGPHWNITDANDESYSLVVRGIHESDAKRIAKWVQS RIPYVRAETQCD |
| avb8_18 | TKCVVRFIFRGDLATIALKAVKDHLKKEGPHWNITTQEEPNQYLVVRGIHESDAKRIAKWVEKR FPFAGVETQCD |

**Supplementary Table 4: SSM forward primers.**

| | |
|---|---|
| av6_3.fasta_1 | AGGGTCGGCTAGCCATATGNNKGCTGTAGTTAGGTTCGTATTCAG |
| av6_3.fasta_2 | GGGTCGGCTAGCCATATGATGNNKGTAGTTAGGTTCGTATTCAGAGGT |
| av6_3.fasta_3 | TCGGCTAGCCATATGATGGCTNNKGTTAGGTTCGTATTCAGAGGTGA |
| av6_3.fasta_4 | GGCTAGCCATATGATGGCTGTANNKAGGTTCGTATTCAGAGGTGAC |
| av6_3.fasta_5 | GCTAGCCATATGATGGCTGTAGTTNNKTTCGTATTCAGAGGTGACTTGG |
| av6_3.fasta_6 | AGCCATATGATGGCTGTAGTTAGGNNKGTATTCAGAGGTGACTTGGCA |
| av6_3.fasta_7 | CCATATGATGGCTGTAGTTAGGTTCNNKTTCAGAGGTGACTTGGCAG |
| av6_3.fasta_8 | TGATGGCTGTAGTTAGGTTCGTANNKAGAGGTGACTTGGCAGAATTG |
| av6_3.fasta_9 | ATGGCTGTAGTTAGGTTCGTATTCNNKGGTGACTTGGCAGAATTGATG |
| av6_3.fasta_10 | GGCTGTAGTTAGGTTCGTATTCAGANNKGACTTGGCAGAATTGATGTTGA |
| av6_3.fasta_11 | TGTAGTTAGGTTCGTATTCAGAGGTNNKTTGGCAGAATTGATGTTGAGGG |
| av6_3.fasta_12 | TAGGTTCGTATTCAGAGGTGACNNKGCAGAATTGATGTTGAGGGC |
| av6_3.fasta_13 | GGTTCGTATTCAGAGGTGACTTGNNKGAATTGATGTTGAGGGCAGTTAA |

| | |
|---|---|
| av6_3.fasta_14 | CGTATTCAGAGGTGACTTGGCANNKTTGATGTTGAGGGCAGTTAAGG |
| av6_3.fasta_15 | TCAGAGGTGACTTGGCAGAANNKATGTTGAGGGCAGTTAAGGAC |
| av6_3.fasta_16 | CAGAGGTGACTTGGCAGAATTGNNKTTGAGGGCAGTTAAGGACCA |
| av6_3.fasta_17 | AGGTGACTTGGCAGAATTGATGNNKAGGGCAGTTAAGGACCACT |
| av6_3.fasta_18 | GTGACTTGGCAGAATTGATGTTGNNKGCAGTTAAGGACCACTTGAAG |
| av6_3.fasta_19 | ACTTGGCAGAATTGATGTTGAGGNNKGTTAAGGACCACTTGAAGAAGGA |
| av6_3.fasta_20 | GGCAGAATTGATGTTGAGGGCANNKAAGGACCACTTGAAGAAGGAAG |
| av6_3.fasta_21 | CAGAATTGATGTTGAGGGCAGTTNNKGACCACTTGAAGAAGGAAGGT |
| av6_3.fasta_22 | GAATTGATGTTGAGGGCAGTTAAGNNKCACTTGAAGAAGGAAGGTCCA |
| av6_3.fasta_23 | ATGTTGAGGGCAGTTAAGGACNNKTTGAAGAAGGAAGGTCCACATT |
| av6_3.fasta_24 | TGAGGGCAGTTAAGGACCACNNKAAGAAGGAAGGTCCACATTGG |
| av6_3.fasta_25 | AGGGCAGTTAAGGACCACTTGNNKAAGGAAGGTCCACATTGGAAC |
| av6_3.fasta_26 | GGCAGTTAAGGACCACTTGAAGNNKGAAGGTCCACATTGGAACATTAC |
| av6_3.fasta_27 | GCAGTTAAGGACCACTTGAAGAAGNNKGGTCCACATTGGAACATTACTTC |
| av6_3.fasta_28 | GTTAAGGACCACTTGAAGAAGGAANNKCCACATTGGAACATTACTTCAACT |
| av6_3.fasta_29 | GGACCACTTGAAGAAGGAAGGTNNKCATTGGAACATTACTTCAACTAACAA<br>C |
| av6_3.fasta_30 | CCACTTGAAGAAGGAAGGTCCANNKTGGAACATTACTTCAACTAACAACG |
| av6_3.fasta_31 | ACTTGAAGAAGGAAGGTCCACATNNKAACATTACTTCAACTAACAACGGAG |
| av6_3.fasta_32 | AAGAAGGAAGGTCCACATTGGNNKATTACTTCAACTAACAACGGAGC |
| av6_3.fasta_33 | AAGGAAGGTCCACATTGGAACNNKACTTCAACTAACAACGGAGCT |
| av6_3.fasta_34 | AGGAAGGTCCACATTGGAACATTNNKTCAACTAACAACGGAGCTGA |
| av6_3.fasta_35 | AAGGTCCACATTGGAACATTACTNNKACTAACAACGGAGCTGAATTGG |
| av6_3.fasta_36 | GGTCCACATTGGAACATTACTTCANNKAACAACGGAGCTGAATTGGT |
| av6_3.fasta_37 | TCCACATTGGAACATTACTTCAACTNNKAACGGAGCTGAATTGGTTGT |
| av6_3.fasta_38 | CCACATTGGAACATTACTTCAACTAACNNKGGAGCTGAATTGGTTGTAAGG |
| av6_3.fasta_39 | ACATTGGAACATTACTTCAACTAACAACNNKGCTGAATTGGTTGTAAGGGG<br>A |
| av6_3.fasta_40 | GGAACATTACTTCAACTAACAACGGANNKGAATTGGTTGTAAGGGGAATTC<br>A |
| av6_3.fasta_41 | TACTTCAACTAACAACGGAGCTNNKTTGGTTGTAAGGGGAATTCATGA |
| av6_3.fasta_42 | TTCAACTAACAACGGAGCTGAANNKGTTGTAAGGGGAATTCATGAGTC |
| av6_3.fasta_43 | CAACTAACAACGGAGCTGAATTGNNKGTAAGGGGAATTCATGAGTCAGA |

| | |
|---|---|
| av6_3.fasta_44 | AACAACGGAGCTGAATTGGTTNNKAGGGGAATTCATGAGTCAGAC |
| av6_3.fasta_45 | AACGGAGCTGAATTGGTTGTANNKGGAATTCATGAGTCAGACGCA |
| av6_3.fasta_46 | CGGAGCTGAATTGGTTGTAAGGNNKATTCATGAGTCAGACGCAAAGA |
| av6_3.fasta_47 | AGCTGAATTGGTTGTAAGGGGANNKCATGAGTCAGACGCAAAGAGA |
| av6_3.fasta_48 | GCTGAATTGGTTGTAAGGGGAATTNNKGAGTCAGACGCAAAGAGAATTG |
| av6_3.fasta_49 | TGAATTGGTTGTAAGGGGAATTCATNNKTCAGACGCAAAGAGAATTGC |
| av6_3.fasta_50 | TTGGTTGTAAGGGGAATTCATGAGNNKGACGCAAAGAGAATTGCTAAGT |
| av6_3.fasta_51 | GTTGTAAGGGGAATTCATGAGTCANNKGCAAAGAGAATTGCTAAGTGGG |
| av6_3.fasta_52 | AAGGGGAATTCATGAGTCAGACNNKAAGAGAATTGCTAAGTGGGTTGA |
| av6_3.fasta_53 | GGGAATTCATGAGTCAGACGCANNKAGAATTGCTAAGTGGGTTGAGA |
| av6_3.fasta_54 | GAATTCATGAGTCAGACGCAAAGNNKATTGCTAAGTGGGTTGAGAAGA |
| av6_3.fasta_55 | TCATGAGTCAGACGCAAAGAGANNKGCTAAGTGGGTTGAGAAGAGG |
| av6_3.fasta_56 | ATGAGTCAGACGCAAAGAGAATTNNKAAGTGGGTTGAGAAGAGGTTT |
| av6_3.fasta_57 | TCAGACGCAAAGAGAATTGCTNNKTGGGTTGAGAAGAGGTTTCC |
| av6_3.fasta_58 | CAGACGCAAAGAGAATTGCTAAGNNKGTTGAGAAGAGGTTTCCAGGA |
| av6_3.fasta_59 | ACGCAAAGAGAATTGCTAAGTGGNNKGAGAAGAGGTTTCCAGGAGTT |
| av6_3.fasta_60 | GCAAAGAGAATTGCTAAGTGGGTTNNKAAGAGGTTTCCAGGAGTTCATAC |
| av6_3.fasta_61 | AGAGAATTGCTAAGTGGGTTGAGNNKAGGTTTCCAGGAGTTCATACTG |
| av6_3.fasta_62 | AGAATTGCTAAGTGGGTTGAGAAGNNKTTTCCAGGAGTTCATACTGAAACA |
| av6_3.fasta_63 | TGCTAAGTGGGTTGAGAAGAGGNNKCCAGGAGTTCATACTGAAACACA |
| av6_3.fasta_64 | TAAGTGGGTTGAGAAGAGGTTTNNKGGAGTTCATACTGAAACACAGC |
| av6_3.fasta_65 | TGGGTTGAGAAGAGGTTTCCANNKGTTCATACTGAAACACAGCAAGA |
| av6_3.fasta_66 | GGTTGAGAAGAGGTTTCCAGGANNKCATACTGAAACACAGCAAGATCT |
| av6_3.fasta_67 | TGAGAAGAGGTTTCCAGGAGTTNNKACTGAAACACAGCAAGATCTCG |
| av6_3.fasta_68 | AGAAGAGGTTTCCAGGAGTTCATNNKGAAACACAGCAAGATCTCGAG |
| av6_3.fasta_69 | AGAGGTTTCCAGGAGTTCATACTNNKACACAGCAAGATCTCGAGG |
| av6_3.fasta_70 | AGGTTTCCAGGAGTTCATACTGAANNKCAGCAAGATCTCGAGGGAG |
| av6_3.fasta_71 | TTTCCAGGAGTTCATACTGAAACANNKCAAGATCTCGAGGGAGGCG |
| av6_3.fasta_72 | CCAGGAGTTCATACTGAAACACAGNNKGATCTCGAGGGAGGCGG |
| av6_3.fasta_73 | GGAGTTCATACTGAAACACAGCAANNKCTCGAGGGAGGCGGAT |

**Supplementary Table 5: SSM reverse primers.**

| | |
|---|---|
| av6_3.fasta_1R | CATATGGCTAGCCGACCCT |
| av6_3.fasta_2R | CATCATATGGCTAGCCGACCC |
| av6_3.fasta_3R | AGCCATCATATGGCTAGCCGA |
| av6_3.fasta_4R | TACAGCCATCATATGGCTAGCC |
| av6_3.fasta_5R | AACTACAGCCATCATATGGCTAGC |
| av6_3.fasta_6R | CCTAACTACAGCCATCATATGGCT |
| av6_3.fasta_7R | GAACCTAACTACAGCCATCATATGG |
| av6_3.fasta_8R | TACGAACCTAACTACAGCCATCA |
| av6_3.fasta_9R | GAATACGAACCTAACTACAGCCAT |
| av6_3.fasta_10R | TCTGAATACGAACCTAACTACAGCC |
| av6_3.fasta_11R | ACCTCTGAATACGAACCTAACTACA |
| av6_3.fasta_12R | GTCACCTCTGAATACGAACCTA |
| av6_3.fasta_13R | CAAGTCACCTCTGAATACGAACC |
| av6_3.fasta_14R | TGCCAAGTCACCTCTGAATACG |
| av6_3.fasta_15R | TTCTGCCAAGTCACCTCTGA |
| av6_3.fasta_16R | CAATTCTGCCAAGTCACCTCTG |
| av6_3.fasta_17R | CATCAATTCTGCCAAGTCACCT |
| av6_3.fasta_18R | CAACATCAATTCTGCCAAGTCAC |
| av6_3.fasta_19R | CCTCAACATCAATTCTGCCAAGT |
| av6_3.fasta_20R | TGCCCTCAACATCAATTCTGCC |
| av6_3.fasta_21R | AACTGCCCTCAACATCAATTCTG |
| av6_3.fasta_22R | CTTAACTGCCCTCAACATCAATTC |
| av6_3.fasta_23R | GTCCTTAACTGCCCTCAACAT |
| av6_3.fasta_24R | GTGGTCCTTAACTGCCCTCA |
| av6_3.fasta_25R | CAAGTGGTCCTTAACTGCCCT |
| av6_3.fasta_26R | CTTCAAGTGGTCCTTAACTGCC |
| av6_3.fasta_27R | CTTCTTCAAGTGGTCCTTAACTGC |
| av6_3.fasta_28R | TTCCTTCTTCAAGTGGTCCTTAAC |
| av6_3.fasta_29R | ACCTTCCTTCTTCAAGTGGTCC |
| av6_3.fasta_30R | TGGACCTTCCTTCTTCAAGTGG |
| av6_3.fasta_31R | ATGTGGACCTTCCTTCTTCAAGT |

| | |
|---|---|
| av6_3.fasta_32R | CCAATGTGGACCTTCCTTCTT |
| av6_3.fasta_33R | GTTCCAATGTGGACCTTCCTT |
| av6_3.fasta_34R | AATGTTCCAATGTGGACCTTCCT |
| av6_3.fasta_35R | AGTAATGTTCCAATGTGGACCTT |
| av6_3.fasta_36R | TGAAGTAATGTTCCAATGTGGACC |
| av6_3.fasta_37R | AGTTGAAGTAATGTTCCAATGTGGA |
| av6_3.fasta_38R | GTTAGTTGAAGTAATGTTCCAATGTGG |
| av6_3.fasta_39R | GTTGTTAGTTGAAGTAATGTTCCAATGT |
| av6_3.fasta_40R | TCCGTTGTTAGTTGAAGTAATGTTCC |
| av6_3.fasta_41R | AGCTCCGTTGTTAGTTGAAGTA |
| av6_3.fasta_42R | TTCAGCTCCGTTGTTAGTTGAA |
| av6_3.fasta_43R | CAATTCAGCTCCGTTGTTAGTTG |
| av6_3.fasta_44R | AACCAATTCAGCTCCGTTGTT |
| av6_3.fasta_45R | TACAACCAATTCAGCTCCGTT |
| av6_3.fasta_46R | CCTTACAACCAATTCAGCTCCG |
| av6_3.fasta_47R | TCCCCTTACAACCAATTCAGCT |
| av6_3.fasta_48R | AATTCCCCTTACAACCAATTCAGC |
| av6_3.fasta_49R | ATGAATTCCCCTTACAACCAATTCA |
| av6_3.fasta_50R | CTCATGAATTCCCCTTACAACCAA |
| av6_3.fasta_51R | TGACTCATGAATTCCCCTTACAAC |
| av6_3.fasta_52R | GTCTGACTCATGAATTCCCCTT |
| av6_3.fasta_53R | TGCGTCTGACTCATGAATTCCC |
| av6_3.fasta_54R | CTTTGCGTCTGACTCATGAATTC |
| av6_3.fasta_55R | TCTCTTTGCGTCTGACTCATGA |
| av6_3.fasta_56R | AATTCTCTTTGCGTCTGACTCAT |
| av6_3.fasta_57R | AGCAATTCTCTTTGCGTCTGA |
| av6_3.fasta_58R | CTTAGCAATTCTCTTTGCGTCTG |
| av6_3.fasta_59R | CCACTTAGCAATTCTCTTTGCGT |
| av6_3.fasta_60R | AACCCACTTAGCAATTCTCTTTGC |
| av6_3.fasta_61R | CTCAACCCACTTAGCAATTCTCT |
| av6_3.fasta_62R | CTTCTCAACCCACTTAGCAATTCT |
| av6_3.fasta_63R | CCTCTTCTCAACCCACTTAGCA |

| | |
|---|---|
| av6_3.fasta_64R | AAACCTCTTCTCAACCCACTTA |
| av6_3.fasta_65R | TGGAAACCTCTTCTCAACCCA |
| av6_3.fasta_66R | TCCTGGAAACCTCTTCTCAACC |
| av6_3.fasta_67R | AACTCCTGGAAACCTCTTCTCA |
| av6_3.fasta_68R | ATGAACTCCTGGAAACCTCTTCT |
| av6_3.fasta_69R | AGTATGAACTCCTGGAAACCTCT |
| av6_3.fasta_70R | TTCAGTATGAACTCCTGGAAACCT |
| av6_3.fasta_71R | TGTTTCAGTATGAACTCCTGGAAA |
| av6_3.fasta_72R | CTGTGTTTCAGTATGAACTCCTGG |
| av6_3.fasta_73R | TTGCTGTGTTTCAGTATGAACTCC |

**Supplementary Table 6: Integrin αvβ6 and αvβ8 headpiece and ectodomain amino acid sequences.**

>Human αv headpiece (with N-term signal peptide; **M430GC**; and C-term 3C cleavage site, **ACID coil**, and 6xH tag)
MAFPPRRRLRLGPRGLPLLLSGLLLPLCRAFNLDVDSPAEYSGPEGSYFGFAVDFFVPSASSRMFLLVGAPKANTTQPGIV
EGGQVLKCDWSSTRRCQPIEFDATGNRDYAKDDPLEFKSHQWFGASVRSKQDKILACAPLYHWRTEMKQEREPVGTCFLQD
GTKTVEYAPCRSQDIDADGQGFCQGGFSIDFTKADRVLLGGPGSFYWQGQLISDQVAEIVSKYDPNVYSIKYNNQLATRTA
QAIFDDSYLGYSVAVGDFNGDGIDDFVSGVPRAARTLGMVYIYDGKNMSSLYNFTGEQMAAYFGFSVAATDINGDDYADVF
IGAPLFMDRGSDGKLQEVGQVSVSLQRASGDFQTTKLNGFEVFARFGSAIAPLGDLDQDGFNDIAIAAPYGGEDKKGIVYI
FNGRSTGLNAVPSQILEGQWAARS**GC**PPSFGYSMKGATDIDKNGYPDLIVGAFGVDRAILYRARPVITVNAGLEVYPSILN
QDNKTCSLPGTALKVSCFNVRFCLKADGKGVLPRKLNFQVELLLDKLKQKGAIRRALFLYSRSPSHSKNMTISRGGLMQCE
ELIAYLRDESEFRDKLTPITIFMEYRLDYRTAADTTGLQPILNQFTPANISRQAHILLTGGLEVLFQGPGEN**AQLEKELQA
LEKENAQLEWELQALEKELAQ**GGSHHHHHH

>Human β6 headpiece (with N-term signal peptide; **I287C**; and C-term 3C cleavage site, **BASE coil**, and 6xH tag)
MAFPPRRRLRLGPRGLPLLLSGLLLPLCRAHVQGGCALGGAETCEDCLLIGPQCAWCAQENFTHPSGVGERCDTPANLLAK
GCQLNFIENPVSQVEILKNKPLSVGRQKNSSDIVQIAPQSLILKLRPGGAQTLQVHVRQTEDYPVDLYYLMDLSASMDDDL
NTIKELGSRLSKEMSKLTSNFRLGFGSFVEKPVSPFVKTTPEEIANPCSSIPYFCLPTFGFKHILPLTNDAERFNEIVKNQ
KISANIDTPEGGFDAIMQAAVCKEKIGWRNDSLHLLVFVSDADSHFGMDSKLAGIV**C**PNDGLCHLDSKNEYSMSTVLEYPT
IGQLIDKLVQNNVLLIFAVTQEQVHLYENYAKLIPGATVGLLQKDSGNILQLIISAYEELRSEVELEVLGDTEGLNLSFTA
ICNNGTLFQHQKKCSHMKVGDTASFSVTVNIPHCERRSRHIIIKPVGLGDALELLVSPECNCDCQKEVEVNSSKCHHGNGS
FQCGVCACHPGHMGPRCESGGLEVLFQGPSGG**AQLKKKLQALKKKNAQLKWKLQALKKKLAQ**GGSHHHHHH

>Human β8 headpiece (with N-term signal peptide; **V301C**; and C-term 3C cleavage site, **BASE coil**, and 6xH tag)
MAFPPRRRLRLGPRGLPLLLSGLLLPLCRAEDNRCASSNAASCARCLALGPECGWCVQEDFISGGSRSERCDIVSNLISKG
CSVDSIEYPSVHVIIPTENEINTQVTPGEVSIQLRPGAEANFMLKVHPLKKYPVDLYYLVDVSASMHNNIEKLNSVGNDLS
RKMAFFSRDFRLGFGSYVDKTVSPYISIHPERIHNQCSDYNLDCMPPHGYIHVLSLTENITEFEKAVHRQKISGNIDTPEG
GFDAMLQAAVCESHIGWRKEAKRLLLVMTDQTSHLALDSKLAGIV**C**PNDGNCHLKNNVYVKSTTMEHPSLGQLSEKLIDNN
INVIFAVQGKQFHWYKDLLPLLPGTIAGEIESKAANLNNLVVEAYQKLISEVKVQVENQVQGIYFNITAICPDGSRKPGME
GCRNVTSNDEVLFNVTVTMKKCDVTGGKNYAIIKPIGFNETAKIHIHRNCSCQCEDNRGPKGKCVDETFLDSKCFQCDENK
SGGLEVLFQGPSGG**AQLKKKLQALKKKNAQLKWKLQALKKKLAQ**GGSHHHHHH

>Human αv ectodomain (with N-term signal peptide; **M430GC**; and C-term 3C cleavage site, **ACID coil**, and 6xH tag)

MAFPPRRRLRLGPRGLPLLLSGLLLPLCRAFNLDVDSPAEYSGPEGSYFGFAVDFFVPSASSRMFLLVGAPKANTTQPGIV
EGGQVLKCDWSSTRRCQPIEFDATGNRDYAKDDPLEFKSHQWFGASVRSKQDKILACAPLYHWRTEMKQEREPVGTCFLQD
GTKTVEYAPCRSQDIDADGQGFCQGGFSIDFTKADRVLLGGPGSFYWQGQLISDQVAEIVSKYDPNVYSIKYNNQLATRTA
QAIFDDSYLGYSVAVGDFNGDGIDDFVSGVPRAARTLGMVYIYDGKNMSSLYNFTGEQMAAYFGFSVAATDINGDDYADVF
IGAPLFMDRGSDGKLQEVGQVSVSLQRASGDFQTTKLNGFEVFARFGSAIAPLGDLDQDGFNDIAIAAPYGGEDKKGIVYI
FNGRSTGLNAVPSQILEGQWAARS**GC**PPSFGYSMKGATDIDKNGYPDLIVGAFGVDRAILYRARPVITVNAGLEVYPSILN
QDNKTCSLPGTALKVSCFNVRFCLKADGKGVLPRKLNFQVELLLDKLKQKGAIRRALFLYSRSPSHSKNMTISRGGLMQCE
ELIAYLRDESEFRDKLTPITIFMEYRLDYRTAADTTGLQPILNQFTPANISRQAHILLDCGEDNVCKPKLEVSVDSDQKKI
YIGDDNPLTLIVKAQNQGEGAYEAELIVSIPLQADFIGVVRNNEALARLSCAFKTENQTRQVVCDLGNPMKAGTQLLAGLR
FSVHQQSEMDTSVKFDLQIQSSNLFDKVSPVVSHKVDLAVLAAVEIRGVSSPDHVFLPIPNWEHKENPETEEDVGPVVQHI
YELRNNGPSSFSKAMLHLQWPYKYNNNTLLYILHYDIDGPMNCTSDMEINPLRIKISSLQTTEKNDTVAGQGERDHLITKR
DLALSEGDIHTLGCGVAQCLKIVCQVGRLDRGKSAILYVKSLLWTETFMNKENQNHSYSLKSSASFNVIEFPYKNLPIEDI
TNSTLVTTNVTWGIQPAPMTGGLEVLFQGPGEN**AQLEKELQALEKENAQLEWELQALEKELAQ**GGSHHHHHH

>Human β6 ectodomain (with N-term signal peptide; **I287C**; and C-term 3C cleavage site, **BASE coil**, and 6xH tag)
MAFPPRRRLRLGPRGLPLLLSGLLLPLCRAHVQGGCALGGAETCEDCLLIGPQCAWCAQENFTHPSGVGERCDTPANLLAK
GCQLNFIENPVSQVEILKNKPLSVGRQKNSSDIVQIAPQSLILKLRPGGAQTLQVHVRQTEDYPVDLYYLMDLSASMDDDL
NTIKELGSRLSKEMSKLTSNFRLGFGSFVEKPVSPFVKTTPEEIANPCSSIPYFCLPTFGFKHILPLTNDAERFNEIVKNQ
KISANIDTPEGGFDAIMQAAVCKEKIGWRNDSLHLLVFVSDADSHFGMDSKLAGIV**C**PNDGLCHLDSKNEYSMSTVLEYPT
IGQLIDKLVQNNVLLIFAVTQEQVHLYENYAKLIPGATVGLLQKDSGNILQLIISAYEELRSEVELEVLGDTEGLNLSFTA
ICNNGTLFQHQKKCSHMKVGDTASFSVTVNIPHCERRSRHIIIKPVGLGDALELLVSPECNCDCQKEVEVNSSKCHHGNGS
FQCGVCACHPGHMGPRCECGEDMLSTDSCKEAPDHPSCSGRGDCYCGQCICHLSPYGNIYGPYCQCDNFSCVRHKGLLCGG
NGDCDCGECVCRSGWTGEYCNCTTSTDSCVSEDGVLCSGRGDCVCGKCVCTNPGASGPTCERCPTCGDPCNSKRSCIECHL
SAAGQAREECVDKCKLAGATISEEEDFSKDGSVSCSLQGENECLITFLITTDNEGKTIIHSINEKDCPKPPNSGGLEVLFQ
GPSGG**AQLKKKLQALKKKNAQLKWKLQALKKKLAQ**GGSHHHHHH

>Human β8 ectodomain (with N-term signal peptide; **V301C**; and C-term 3C cleavage site, **BASE coil**, and 6xH tag)
MAFPPRRRLRLGPRGLPLLLSGLLLPLCRAEDNRCASSNAASCARCLALGPECGWCVQEDFISGGSRSERCDIVSNLISKG
CSVDSIEYPSVHVIIPTENEINTQVTPGEVSIQLRPGAEANFMLKVHPLKKYPVDLYYLVDVSASMHNNIEKLNSVGNDLS
RKMAFFSRDFRLGFGSYVDKTVSPYISIHPERIHNQCSDYNLDCMPPHGYIHVLSLTENITEFEKAVHRQKISGNIDTPEG
GFDAMLQAAVCESHIGWRKEAKRLLLVMTDQTSHLALDSKLAGIV**C**PNDGNCHLKNNVYVKSTTMEHPSLGQLSEKLIDNN
INVIFAVQGKQFHWYKDLLPLLPGTIAGEIESKAANLNNLVVEAYQKLISEVKVQVENQVQGIYFNITAICPDGSRKPGME
GCRNVTSNDEVLFNVTVTMKKCDVTGGKNYAIIKPIGFNETAKIHIHRNCSCQCEDNRGPKGKCVDETFLDSKCFQCDENK
CHFDEDQFSSESCKSHKDQPVCSGRGVCVCGKCSCHKIKLGKVYGKYCEKDDFSCPYHHGNLCAGHGECEAGRCQCFSGWE
GDRCQCPSAAAQHCVNSKGQVCSGRGTCVCGRCECTDPRSIGRFCEHCPTCYTACKENWNCMQCLHPHNLSQAILDQCKTS
CALMEQQHYVDQTSECFSSPSSGGLEVLFQGPSGG**AQLKKKLQALKKKNAQLKWKLQALKKKLAQ**GGSHHHHHH

**Supplementary Table 7: CryoEM data collection, refinement, and validation statistics.**

| | **αvβ8 + B8_BP_dslf (EMDB-xxxx) (PDB xxxx)** | **αvβ6 + B6_BP_dslf (EMDB-xxxx) (PDB xxxx)** |
|---|---|---|
| **Data collection and processing** | | |
| Magnification | 44563 (36000 nominal) | 44563 (36000 nominal) |
| Voltage (kV) | 200 | 200 |
| Electron exposure (e–/Å$^2$) | 50 | 50 |
| Defocus range (μm) | 1.0-1.8 (nominal) | 1.0-1.8 (nominal) |
| Pixel size (Å) | 1.122 | 1.122 |
| Symmetry imposed | C1 | C1 |

| | | |
|---|---|---|
| Initial particle images (no.) | 1,273,469 | 360,600 |
| Final particle images (no.) | 134,395 | 54,011 |
| Map resolution (Å) | 2.9 | 3.4 |
| FSC threshold | .143 | .143 |
| Map resolution range (Å) | 2.6-7.2 | 3.2-7.0 |
| **Refinement** | | |
| Initial model used (PDB code) | 6UJA (AB) | 5FFO (EF) |
| Model resolution (Å) | 2.9 | 3.4 |
| FSC threshold | .143 | .143 |
| Model resolution range (Å) | 2.6-7.2 | 3.2-7.0 |
| Map sharpening *B* factor ($Å^2$) | -51 | -51 |
| Model composition | | |
| Non-hydrogen atoms | 7050 | 5945 |
| Protein residues | 878 | 754 |
| Ligands | 0 | 0 |
| *B* factors ($Å^2$) | | |
| Protein | 30 | 79 |
| Ligand | n/a | n/a |
| R.m.s. deviations | | |
| Bond lengths (Å) | 0.013 | 0.013 |
| Bond angles (°) | 1.260 | 1.565 |
| Validation | | |
| MolProbity score | 0.99 | 0.85 |
| Clashscore | 0.72 | 0.34 |
| Poor rotamers (%) | 0.41 | 0.16 |
| Ramachandran plot | | |
| Favored (%) | 96 | 97 |
| Allowed (%) | 4 | 3 |
| Disallowed (%) | 0 | 0 |

### Materials and Methods

#### Computational techniques

Overview of the design protocol has been discussed in the main text. Detailed step by step protocol and all associated codes have been deposited on github (https://github.com/aroy10/avb6-publication) and available from the main authors upon request. XMLs for designing the disulfide bond discussed in the main text have also been deposited in the same repository.

Structures were assembled from fragments following rules for constructing ideal proteins,[25] sampling different alpha helix, beta sheet, and loop lengths, while constraining torsion angles in the region corresponding to the RGD peptide to those observed in the co-crystal structure using Rosetta. The resulting idealized ferredoxin fold structures were docked in complex with the αvβ6 integrin by superposition on the binding loop, and the amino acids at the binding surface were optimized for low energy interactions with the target. We solved the crystal structure of the best binder at 1.8 Å from the first round of design following a round of optimization using error prone PCR. The crystal structure fits the designed model well except for a rigid body translation of the C-terminal helix along the helical axis by one helical turn (~3.4 Å; Extended Data Fig. 1a).

In a second round of design we docked the crystal structure of the Round 1 designed binder onto αvβ6 by superimposing on the RGD loop. We identified two loop regions in the design close to the integrin and sampled a range of lengths and conformations for the two loops, and selected 16 designs with loops predicted to make selective interactions with the integrin for experimental testing (Extended Data Figs. 1b, 2, 3). Variant av6_3 with highest affinity towards αvβ6 was further subjected to site-saturation mutagenesis (SSM) (Extended Data Figs. 4, 5) with increasing stringency. Five substitutions at the interface which primarily increase charge complementarity were enriched (Extended Data Fig. 5d-h). αvβ6 not only recognises the RGD loop but also an amphipathic helix formed by the LXXL motif that interacts only with the β6 subunit,[1] and substitutions in the region of the design corresponding to this motif were strongly disfavored for β6 binding (Extended Data Fig. 5b). We expressed and purified 9 enriched variants and measured binding for αvβ6 integrin using biolayer interferometry (BLI) measurements; all had subnanomolar binding affinity (the original av6_3 binds to αvβ6 with a $K_d$ of 1.2 (±0.006) nM; Extended Data Fig. 6, Extended Data Table 1). Two high affinity variants were selected for further characterization: B6B8_BP (av6_3_E13T) with a single substitution and B6_BP (av6_3_A39KG64R) with two substitutions introducing positive charges complementing negative charges in both subunits of the αvβ6 integrin (Extended Data Figs. 5, 6).

**Yeast display**

Standard yeast surface display techniques were used to screen designs for binding and directed evolution. Genes encoding the designs were cloned into petcon2 in frame with N-term aga2 and C-term myc tag. Surface expression of myc was detected using FITC conjugated chicken anti-C-myc (Immunology Consultants Laboratory, Inc) and binding was detected using biotinylated human αvβ6 and stained with phycoerythrin conjugated streptavidin (Life technologies) for FACS. αvβ6 was chemically biotinylated to the lysine residues using EZ-Link Sulfo-NHS-LC-Biotin and biotinylation kits following manufacturer protocol. Excess biotin was removed from the mixture by dialyzing against a buffer containing no biotin. Two different buffers were used for the binding and washing steps for yeast display; Binding Buffer: 20 mM Tris, 150 mM NaCl, pH=8.0, 1% BSA, 1 mM $Ca^{2+}$ and 1 mM $Mg^{2+}$, Wash Buffer: 20 mM Tris, 150 mM NaCl, pH=8.0, 0.5% BSA, 1 mM $Ca^{2+}$ and 1 mM $Mg^{2+}$.

The SSM library was generated by using mutagenic primers (see below for sequences, Table S6 and S7) for each position following a previously described protocol[2,3]. The resulting library was transformed into yeast using electroporation in duplicates (biological replicate). The sorting was performed in two rounds: The library was first treated with 4 µM Trypsin and 0.8 µM chymotrypsin[4] for 45 secs followed by labeling with 200 pM of biotinylated αvβ6 and top 5% of the binders were collected. For the second and final round of selection, 100 pM of biotinylated αvβ6 was used and the top 1% of the binding population was selected (Extended Data Fig. 4). DNA was extracted from pre and post sorted pools and barcoded. Enrichment ratios were calculated after sequencing the pools using Illumina.

**Protein minibinder expression and purification**

Genes encoding protein variants were ordered as gblock gene fragments from IDT and cloned in pet29b in between NdeI/XhoI restriction sites with a C-term Histag, or directly ordered from IDT already cloned into pet29b. All the mutant variants of the proteins were expressed in BL21(DE3*) using Studier autoinduction technique in standard shake flasks at 37℃ for 24hrs. Cells were harvested and resuspended in 20 mM Tris, 250 mM NaCl, 20 mM Imidazole (lysis buffer). Cells were lysed using microfluidizer and cell debris was separated by centrifuging at 24000g for 45 mins. Soluble proteins were first purified using standard Ni-NTA affinity columns followed by size exclusion chromatography (S75 10/300 Increase) on a GE-Akta pure FPLC system. Peak corresponding to the monomeric protein was collected and further verified by mass spectrometry. For bleomycin induced PF models, protein was subjected to further purification to achieve endotoxin level <5 EU/ml.

**Integrin αvβ6 and αvβ8 DNA constructs**

Wild-type human integrin αvβ6 headpiece[5] and αvβ6 and αvβ8 ectodomains[6,7] have been described. Here, the αvβ6 and αvβ8 headpiece and ectodomain constructs were synthesized by GenScript into the pCMV/R vector with an N-terminal signal peptide and C-terminal Human Rhinovirus (HRV) 3C protease cleavage site, ACID coiled coil in the αv (UniProt: P06756) subunit or a BASE coiled coil in the β6 (UniProt: P18564) and β8 (UniProt: P26012) subunits, and hexa-histidine tag. The αv, β6, and β8 subunits each had a cysteine mutation (M430GC, I287C, and V301C, respectively) to generate a disulfide bond to prevent α/β subunit dissociation following 3C cleavage. The Gly inserted prior to residue 430 in the M430GC mutation in the αv subunit was previously reported[8]. Plasmids were transformed into the NEB 5-alpha strain of *E. coli* (New England Biolabs) for subsequent DNA extraction from bacterial culture (Qiagen Plasmid Plus Maxi Kit) to obtain plasmid for transient transfection into Expi293F cells. The amino acid sequences for the αvβ6 and αvβ8 headpieces and ectodomains are listed in Supplementary Table 6.

**Secreted integrin αvβ6 and αvβ8 expression and purification**

For integrin headpieces, 800 mL cultures of Expi293F cells were grown in suspension to a density of $3.0\times10^6$ cells per mL and transiently transfected using PEI-MAX (Polyscience) and cultivated for 5 days in Expi293F expression medium (Life Technologies) at 37°C, 70% humidity, 8% CO2, and rotating at 150 rpm. Supernatants were clarified by centrifugation (5 min at 4000 rcf), PDADMAC solution was added to a final concentration of 0.0375% (Sigma Aldrich, #409014), and a final spin was performed (5 min at 4000 rcf). Clarified supernatant was supplemented with 1 M Tris-HCl pH 8.0 to a final concentration of 45 mM and 5 M NaCl to a final concentration of ~310 mM. His-tagged integrins were purified from clarified supernatants via a batch bind method where Ni Sepharose excel resin (Cytiva) was added to the treated supernatants and allowed to incubate overnight at 4°C with gentle shaking. Resin was isolated using 0.2 μm vacuum filtration and transferred to a gravity column, where it was washed with 20 mM Tris pH 8.0, 300 mM NaCl, and protein was eluted with 3 column volumes of 20 mM Tris pH 8.0, 300 mM NaCl, 300 mM imidazole. Eluted protein was concentrated in 50K MWCO centrifugal filters (Millipore), sterile filtered (0.22 μm), and applied to a Superdex 200 Increase 10/300 SEC column (Cytiva) using 20 mM Tris pH 8.0, 150 mM NaCl, 5% glycerol buffer on an AKTA Pure25 FPLC system (Cytiva). SDS-PAGE was used to assess purity and proper integrin dimerization.

For αvβ6 integrin headpiece and αvβ8 integrin ectodomain used in negative-stain or cryoEM analysis, the GenScript plasmids (described above) containing either the αv and β6 headpieces (αvβ6) or the αv and β8 ectodomains (αvβ8) were co-transfected into ExpiCHO cells (ThermoFisher) and grown per the manufacturer's 'Max Titer' recommendations. In brief, cells were grown in suspension at 37˚C, 8% $CO_2$, and ~90% humidity. Cultures were co-transfected with plasmids encoding an alpha and beta subunit with Expifectamine CHO. One day post-transfection cells were supplemented with Enhancer and Feed, and cultures were then moved to a 32˚C, 5% $CO_2$ incubator. Five days post-transfection, supernatant was harvested and clarified via centrifugation before affinity purification using a 5 mL HisTrap FF Crude column (Cytiva). Eluted protein was pooled, concentrated and purified via gel filtration chromatography using a Superdex 200 Increase 10/300 SEC column (Cytiva) that had been equilibrated with 20 mM Tris-HCl pH=7.4, 150 mM NaCl, 1 mM $MgCl_2$, and 1 mM $CaCl_2$. Peak fractions were pooled, concentrated, and incubated overnight at 4˚C with 1:20 3C PreScission protease to cleave the ACID-BASE coils. The following day glycerol (10% v/v) was added and samples were snap-frozen and stored at -80˚C.

**Synthesis of PLN-74809**

PLN-74809 was identified as Compound 5 in a patent application from Pliant Therapeutics[9] and was synthesized to >97.5% HPLC purity by WuXi STA (Shanghai, China).

**Biotinylation of designed proteins**

To generate mono-biotinylated proteins, avi-tag sequence (GLNDIFEAQKIEWHE) was introduced to the N-term of the proteins. Proteins were biotinylated either by co-transforming protein of interest along with pBirA, a vector encoding *E.coli* biotin ligase for *in vivo* biotinylation or using purified protein and an *in vitro* biotinylation kit from Avity using manufacturer's protocol. Biotinylation was further confirmed via mass spec.

**Structural analysis of designed proteins**

For determining the crystal structure of B6B8_BP_dslf, we expressed B6B8_BP_dslf with a N- terminal TEV cleavable histag. After protein expression and purification, B6B8_BP_dslf was treated with (1/100) dilution of stock TEV protease and incubated overnight at room temperature dialyzing against TBS. Following the completion of the cleavage (as monitored via SDS-page gel), proteins were run over a second gravity Ni-NTA column to separate cut his-tag and his-tagged-TEV from cleaved protein.

Following the his-tag cleavage, protein was concentrated to ~50mg/ml and subjected to crystallization trials. Both Binding protein and B6B8_BP_dslf were crystallized by vapor diffusion at 24°C by mixing with an equal volume of reservoir solution: 0.2 M $KNO_3$, 20% PEG3350 (Binding protein) and 0.2 M tripotassium citrate, 20% PEG3350 (B6B8_BP_dslf). Crystals were briefly cryo-soaked in a reservoir solution containing 15% PEG200 and flash-frozen in liquid nitrogen. Diffraction data were collected at the GM/CA beam line of Advanced Photon Source (APS) at -173°C using a MAR225 CCD detector and processed using XDS.

The diffraction data for binding protein were originally scaled to $P6_122$ space group with large Patterson peaks 1/3 and 2/3 along the c axis indicating two translational NCS molecules along the c axis. A solution was found using molecular replacement with the designed model. Autobuild was able to rebuild most sequences in the model, but R and Rfree were still very high, at 44%/47% with reasonably good electron density maps. Data were then re-scaled to the $P3_1$ space group with 12 molecules per asymmetric unit and refined with tetrahedral twinning with three twin laws: -k, -h, -l; k, h, -l; and -h, -k, l. AUTOBUILD was used to build one-third of the sequence and was used several times in the first few of many iterative steps of manual building in COOT[10] and refinement with PHENIX and RefMAC. MolProbity[11] was used to validate the final structure.

**Negative-stain EM sample preparation**

The integrin-minibinder complexes were formed using a 1:2 integrin to minibinder ratio, incubated at room temperature for 60 min, and diluted to a final concentration of 10 µg/mL in 20 mM Tris-HCl pH=7.4, 150 mM NaCl, 1 mM $MgCl_2$, and 1 mM $CaCl_2$. For both experiments, 3 µL of sample was applied to a glow-discharged 400 mesh copper glider grid that had been covered with a thin layer of continuous amorphous carbon. The specimens were stained with a solution containing 2% (wt/vol) uranyl formate as previously described (PMCID: PMC389902).

**Negative-stain EM data acquisition and processing**

Data were acquired using a Thermo Fisher Scientific Talos L120C transmission electron microscope operating at 200 kV and recorded on a 4k × 4k Thermo Fisher Scientific Ceta camera at a nominal magnification of 92,000× with a pixel size of 0.158 nm. Leginon[12] was used to collect 296 (αvβ6) or 337 (αvβ8) micrographs at a nominal range of 1.8–2.2 µm under focus and a dose of approximately 50 $e^-/Å^2$.

Experimental data were processed using cryoSPARC[13] and CTFFIND4[14] within the cryoSPARC wrapper. Initially, 34,794 (αvβ6) or 36,710 (αvβ8) particles were picked using an unbiased blob picker and subjected to three rounds of reference-free 2D alignment and classification to remove false positive particle images. The final particle counts contributing to 2D class averages were 22,096 (αvβ6) and 24,532 (αvβ8).

**CryoEM sample preparation**

The integrin-minibinder complexes were formed using a 1:2 integrin to minibinder ratio, incubated at room temperature for 60 min, subjected to size exclusion chromatography, and concentrated to 2-3 mg/mL. From there, complexes were diluted to a final concentration of 0.90 mg/mL (αvβ6) or 0.96 mg/mL (αvβ8) in 20 mM Tris-HCl pH=7.4, 150 mM NaCl, 1 mM $MgCl_2$, and 1 mM $CaCl_2$. For cryoEM grid preparation, both Quantafoil and UltrAufoil grids were glow-discharged for 60s at 15 mA. Just prior to sample application, 10% CHAPS detergent was added to each complex up to a final concentration of 0.025%. From there, 3 µL of each complex was added to each grid. Both complexes were frozen with a Thermo Fisher Scientific Vitrobot Mark IV using a 4s blot time (αvβ6) or 4s and 10s blot times (αvβ8). All grids were frozen with 100% humidity at 4°C and plunge-frozen in liquid ethane cooled by liquid nitrogen.

**CryoEM data acquisition and processing**

Four datasets (one for αvβ6, three for αvβ8) were acquired on a Thermo Fisher Scientific Glacios cryo-transmission electron microscope operating at 200 kV and recorded with a Gatan K3 Direct Detection Camera. Automated data collection was carried out using the SerialEM software.(Mastronarde 2005) Ninty-nine frame movies were recorded in super-resolution mode with a super-resolution pixel size of 0.561 Å/px and a nominal magnification of 36kx. Each dataset was collected in a single session with a nominal defocus range of 1.0–1.8 µm under focus and a dose of approximately 50 $e^-/Å^2$. For the αvβ6 complex, one dataset of 2,152 micrographs was collected from an UltrAufoil grid with the stage tilted to 35°. For the αvβ8 complex, three datasets were collected from a Quantafoil grid with a stage tilt of 0˚ (1,460 micrographs) or a singleUltrAufoil grid with the stage tilted to of 30° (570 micrographs) or 35° (3,174 micrographs).

Dose fractionated super-resolution image stacks were motion corrected and binned 2x2 by Fourier cropping using MotionCor2 (**PMID: 28250466**). Motion corrected stacks were then processed using cryoSPARC[13] and CTFFIND4[14] within the cryoSPARC wrapper. Initially, 360,600 (αvβ6) or 3,325,476 (αvβ8) particles were picked using the unbiased blob picker in cryoSPARC and subjected to multiple rounds of reference-free 2D and 3D alignment and classification as outlined in Extended Data Figure 19. . The particle counts contributing to the final 3D structures into which the models were built are 54,011 (αvβ6) and 116,587 (αvβ8). Images showing EM maps were generated using UCSF Chimera (PMID: 15264254), UCSF ChimeraX (PMID: 32881101).

**Model Building**

The initial atomic models used for the αvβ6 - minibinder complex were the αvβ6 headpiece (PDB: 5FFO, chains E and F) and the designed B6_BP_dslf model. The initial models used for the αvβ8 - minibinder complex were αvβ8 (PDB: 6UJA) and the designed B8_BP2_dslf model. Initial models were fit into their respective cryoEM density using UCSF Chimera (PMID: 15264254) and manually adjusted in COOT (PMID: 20383002). Glycans were manually added using COOT. Models (including glycans) were refined and relaxed using Rosetta. Modeling was aided by using cryoEM maps that were focused on specific regions, using sharpened and unsharpened maps. All maps used for modeling have been deposited.

**Biophysical characterization of designed proteins**

Protein secondary structure and thermal stability were measured using the JASCO-1500 CD instrument. For wavelength scan, 10-15 µM of protein in TBS (20 mM Tris, 50 mM NaCl, pH=8.0) was used. The CD spectra were measured from 240-195 nm with a scan rate of 100 nm/min. For thermal melt experiments, signal intensity at 222 nm was monitored as a function of temperature (4°C-95°C) with a temperature gradient of 2°C/min. The sample was held at the specified temperature for at least 5 sec before the measurement. To investigate the role of the engineered disulfide bond on stability, 1 mM TCEP was added to the protein to measure thermal stability under reducing conditions.

**Nebulization of B6_BP_dslf**

For stability studies, 6 mL protein solution was added to the reservoir of an Aeroneb nebulizer (Kent Scientific item #AG-AL7000 and either item #AG-AL1000 [4.0-6.0 um VMD) or item #AG-AL1100 [2.5-4.0 um VMD]). The solution was nebulized for approximately 10 minutes while collecting any condensate in a 50 mL Falcon tube placed in a dry ice and ethanol bath (dry ice and ethanol facilitate deposition of the aerosol). Condensate in the Falcon tube and any residual volume remaining in the nebulizer reservoir were both collected and analyzed for stability and activity. Procedure was subsequently repeated using 6 mL of a 1:10 protein solution diluted in PBS.

**Biolayer interferometry for determining binding kinetics of minibinder proteins**

Data was collected on an Octet RED96 (Forte Bio) and processed using the instrument's software. His Tagged and avi-tagged protein binders were immobilized on Ni-NTA and streptavidin sensor tips. The tips were then dipped into wells containing different concentrations of αvβ6 and αvβ8. Association and dissociation steps were recorded for 900s and 1200s respectively. An empty sensor with no loaded binding protein was included to discard any non-specific binding of αvβ6 to the octet tip.

**Fluorescent labeling of designed binders**

For *in vitro* binding assay and *in vivo* imaging experiments, designed binders were labeled with Alexa Fluor™ 488 C5 Maleimide and Alexa Fluor™ 680 C2 Maleimide (Thermo Fisher Scientific), respectively, via a C-term single cysteine variant. In a typical labeling experiment, 50-200 µM of proteins were reduced with 1 mM TCEP for 30 minutes at room temperature. 3-5 molar excess of the maleimides were added to the protein solution and tumbled at room temperature overnight. The reaction mixture was then purified on a S75 Increase 10/300 column (GE healthcare) to separate free dye from the labeled proteins. Fluorophore conjugation was further confirmed by mass spectrometry.

***In vitro* and *in vivo* binding assays using fluorescently-labeled binders**

Epidermoid cancer cells (A431) and human embryonic kidney 293T cells (HEK 293T) were purchased from American Type Culture Collection (ATCC) and grown in Dulbecco's Modified Eagle Medium (DMEM, Gibco) supplemented with 10% fetal bovine serum (FBS, Gibco) in a humidified atmosphere with 5% CO2 at 37°C. Binding assays were performed on A431 carcinoma cells. A431 cells were dissociated from culture flasks with enzyme-free cell dissociation buffer (Gibco). Varying concentrations of AG-AF488 and E13T-AF488 were incubated with $5x10^4$ A431 cells in 1X TBS with 0.1% BSA, 1 mM $Ca^{2+}$, and 1 mM $Mg^{2+}$ (BTBS) rotating in suspension for 5 hours at 4°C. Sufficient incubation volumes were used to avoid >5% ligand depletion. After incubation, cells were washed with BTBS and analyzed by flow cytometry on a Accuri C6 instrument (BD Biosciences) or Invitrogen Attune NxT (Invitrogen), and data were quantified using FlowJo software (TreeStar). $K_d$ values were determined by fitting the data to a one site– specific binding curve using Prism 7 (GraphPad Software).

For *in vivo* imaging experiments; approximately 1.5 x $10^7$ A431 cells and 1.5 x $10^7$ HEK 293T cells were suspended in 25 µL DMEM supplemented with 10% FBS along with 25 µL Matrigel Basement Membrane Matrix (Corning, #354234) and injected into the left and right shoulder, respectively, of 6-8 week old female athymic nude mice (Jackson Laboratories, NU/J #002019, homozygous for Foxn1nu). Mice were imaged when tumors reached 5-10 mm in diameter. Tumor Bearing mice were injected via the tail vein with 1.5 nmol AF680-labeled proteins in 100 µL of 20 mM Tris buffer (pH 7.4) and 150 mM NaCl, and imaged with a IVIS Lumina Series III system (PerkinElmer) at the indicated time points. The AF680 fluorophore was excited at 615-665 nm and emission was analyzed at 695-770 nm. In each image, a mouse injected with PBS alone was included as a negative control to allow measurement of background signals for data processing.

**Inhibition of αvβ6-mediated TGF-β activation by B6_BP_dslf using a TMLC co-culture assay**

TGF-β activation was measured using transformed mink lung epithelial cells (TMLC) stably transfected with part of the plasminogen activated inhibitor 1 (PAI-1) reporter conjugated to a luciferase reporter (from Professor Daniel Rifkin (New York University, NY, USA)[15]. TMLCs were plated in 96 well plates (15,000 cells/well) in DMEM (Gibco 41966) supplemented with 1% FBS, 10 U/ml Penicillin G and 10 ug/ml Streptomycin G sulphate and allowed to adhere for 3 hours. K562 cells stably transfected with αvβ6[16] were incubated with B6B8_BP, B6_BP, or media alone ('no BP' control) or a range of control antibodies for 15 minutes to allow binding to occur, and then co-cultured with the TMLCs overnight (final concentration 60,000 αvβ6 K562 cells/well). TMLCs incubated alone or co-cultured with parental K562 cells were included as negative controls. Recombinant TGF-β1 (1 ng/ml R&D Systems) was included as a positive control for the PAI-1 luciferase reporter response to active TGF-β1, and a number of antibodies were included in the TMLC:αvβ6 K562 co-culture as additional controls. Anti-TGFβ1, 2, 3 mIgG1 clone 1D11 (20 ug/ml) was included to confirm the effect of complete inhibition of TGF-β1 activity in this assay, along with the corresponding mouse IgG1 isotype control (both antibodies R & D Systems). A titration of anti-avβ6 3G9 hIgG1 monoclonal antibody (1.4-1000 ng/ml, known clinically as STX-100, formerly known as 6.3G9[7]) was included as a positive control for inhibition of avβ6 in this assay along with a human IgG1 isotype control. Additional antibodies were included at single (high) concentrations to indicate the effect of maximal inhibition with each antibody. 264RAD (40 µg/ml) was included as this is a neutralising antibody to both avβ6 and avβ8.[16] An anti-αv (CD51) antibody (20 mg/ml, Enzo Life Sciences) was included to confirm the effect of inhibiting multiple integrins containing the αvsubunit. Unless indicated otherwise, antibodies were manufactured by AstraZeneca, UK. After 18-20 hours, the supernatants were removed and cells were washed in PBS and then lysed in 100 µl of reporter lysis buffer (1x) (Promega E397A) followed by a freeze thaw cycle to ensure complete lysis. Luciferase activity was quantified using the Luciferase assay system (Promega E1501) according to the manufacturer's instructions. 80 µl of lysate was transferred to white-walled luminescence plates (Perkin Elmer), followed by addition of 100 µl of Luciferase assay reagent and the luminescence signal read on a luminometer (Envision, Perkin Elmer).

**Competitive inhibition of αvβ6- and αvβ8-mediated TGF-β activation by B6_BP_dslf using a CAGA co-culture assay**
TGF-β activation was measured using HEK293 cells stably transfected with a reporter construct consisting of 12 repeats of the CAGA TGF-β responsive element upstream of the luciferase gene (from Professor Tom Thompson (University of Cincinnati, Cincinnati, OH, USA)[(Cash et al. 2012)]. Expi293F cells were transfected with αvβ6, αvβ8, or empty vector, or co-transfected with GARP and TGF-β1 with an N-terminal FLAG tag. After two days in FreeStyle culture medium, cells were harvested for the co-culture assay. Cell surface expression of αvβ6 and αvβ8 was confirmed by staining with integrin β6 subunit-specific 7.1g10 antibody[7] and the integrin β8 subunit-specific ADWA11 antibody[18], respectively, followed by secondary fluorescent detection in FACS using AlexaFluor647 goat anti-mouse (Invitrogen A21235). Cell surface expression of GARP/TGF-β1 complexes was confirmed by direct fluorescent detection of APC-labelled anti-FLAG (clone L5, BioLegend) 637308) staining in FACS. CAGA-reporter cells (15,000 cells) were plated with αvβ6 or αvβ8 EXPi293F transient transfectants (15,000 cells) in 96 well plates (30,000 cells/well) in FreeStyle (Gibco). Expi293F cells transiently co-transfected with GARP and TGF-β1 (5,000 cells) and empty vector transfected cells (10,000) were mixed with 2-fold serial dilutions of B6_BP_dslf, irrelevant nanobody, inhibitory 7.1g10 antibody (integrin β6 subunit specific)[7], or inhibitory ADWA11 antibody (integrin β8 subunit specific)[18] in FreeStyle media and then immediately co-cultured with the CAGA-reporter cells and integrin transfectants overnight (for a final total amount of 45,000 cells/well in 100 µl of media). Serial dilutions were prepared in Freestyle media supplemented with 0.1% BSA to prevent protein loss due to adherence to plastic. CAGA-reporter cells co-cultured with GARP/TGF-β1 co-transfectants and empty vector (mock) transfectants were included to determine the background level of integrin-independent TGF-β1 activation, and CAGA-reporter cells co-cultured with GARP/TGF-β1 co-transfectants and αvβ6 or αvβ8 transfectants were used to determine the total level of TGF-β1 activation in the presence of either integrin. After 18-20 hours, the supernatants were removed and cells were incubated in 50 µl of reporter lysis buffer (1x) (Promega E397A) on ice for 30 minutes. Luciferase activity was quantified using the Luciferase assay system (Promega E1501) according to the manufacturer's instructions. 40 µl of lysate was transferred to white-walled luminescence plates (Perkin Elmer), followed by addition of 40 µl of Luciferase assay reagent and the luminescence signal read on a luminometer (Biotek Synergy H1).

**Human integrin αvβ6 and αvβ8 Latency Associated Peptide-1 ($hLAP_1$)-binding assays**
Recombinant human integrin αvβ6 or αvβ8 headpieces at 1.0 µg/mL in TBS (25 mM Tris pH 8.0, 150 mM NaCl) were incubated overnight at 4°C on 96-well Nunc MaxiSorp plates (ThermoFisher, #442404) (100 µL/well). Plates were blocked with 200 µL/well of blocking buffer (2% [w/v] bovine serum albumin [BSA] in TBS) and incubated for 1 h at 37°C. Plates were washed 3× in binding/wash buffer (25 mM Tris pH 8.0, 150 mM NaCl, 1 mM $CaCl_2$, 1 mM $MgCl_2$, 1 mM $MnCl_2$, 0.1% [w/v] BSA, 0.02% [v/v] Tween20) using a robotic plate washer (BioTek). Serial dilutions of B6_BP_dslf, B6B8_BP_dslf, B8_BP_dslf, and PLN-74809 starting at 100 nM and diluting 3-fold seven times and 1 µg/mL (for αvβ6) or 0.25 µg/mL (for αvβ8) recombinant human $LAP_1$ (R&D Systems, #246-LP-025/CF) in binding/wash buffer were added (100 µL/well) and incubated for 2 h at room temperature. After washing plates 3× in binding/wash buffer, plates were incubated for 1 h at room temperature with 0.5 µg/mL biotinylated mouse IgG2a anti-$hLAP_1$ antibody in binding/wash buffer (R&D Systems, clone #27240, #BAM2462) (100 µL/well). Following washing plates 3× in binding/wash buffer, plates were incubated with peroxidase-conjugated streptavidin (Vector Labs, #SA-5014-1) in binding/wash buffer for 30 min at room temperature (100 µL/well). Following a final 3× plate wash in binding/wash buffer, 100 µL TMB (3,3′,5′,5-tetramethylbenzidine) substrate (SeraCare, #5120-0083) was added to each well, and then TMB was immediately quenched with 1 N HCl (100 µL/well). Absorbance at 450 nm was immediately collected for each well on an Agilent BioTek Epoch 2 plate reader. Data were plotted in Prism 9 (GraphPad Software, San Diego, CA) to determine the 50% inhibitory concentration ($IC_{50}$) using a four-parameter logistic regression model.

**Animals**

For the BLM induced PF model, all mice were housed in Brigham Young University's pathogen-free facility and all experimentation was done in accordance with the protocol approved by the IACUC of Brigham Young University. These protocols conform to both institutional and national guidelines and regulations. Mice were randomly assigned to the treatment groups before any experimental assessments were made. Each cage had a combination of treatment groups to control for cage variation.

For pharmacokinetic analysis of B6_BP_dslf following different routes of administration, male C57BL/6 mice (8-12 weeks old) were obtained from Jackson Laboratory (Strain# 000664, Bar Harbor, Maine) and maintained at the Comparative Medicine Facility at the University of Washington, Seattle, WA, accredited by the American Association for the Accreditation of Laboratory Animal Care International (AAALAC). Animal procedures were performed under the protocol #4470-01 approved by the Institutional Animal Care and Use Committee (IACUC) of University of Washington, Seattle, WA.

**Pharmacokinetic analysis of B6_BP_dslf in healthy mice**

Pharmacokinetics of B6_BP_dslf via the intravenous (IV), intraperitoneal (IP), and intranasal (IN) routes was investigated in healthy male C57BL/6 mice (N=5-7 per time point, body weight 19-22 g) housed in standard holding cages and maintained in a controlled environment with free access to food and water in the Comparative Medicine Facility at the University of Washington, Seattle, WA. B6_BP_dslf, dissolved in 25 mM Tris pH 8.0, 150 mM NaCl, was dosed under isoflurane anesthesia by the IV (via retro-orbital injection) and IP routes at 2 mg/kg and via the IN route at 4 mg/kg. For IV and IP routes, a fixed volume of 50 µL was used. For IN administration, mice were lightly scruffed to create a vertical line from the nose to the lung and a fixed volume of 20 µL (10 µL/nostril) was dripped into the nostrils and allowed to be inhaled. Animals were returned to their home cage and propped up on their back to recover. Lungs were perfused with ~15 mL TBS via the right heart ventricle and harvested and blotted dry on a time course of 5 min to 48 hours after B6_BP_dslf administration; lung weight was recorded and lung was flash-frozen in liquid nitrogen and stored at -80°C until homogenization. Blood samples were also obtained over the same time course via sampling by terminal cardiac puncture. Serum was isolated from hematocrit via centrifugation at 2,000 × *g* for 10 min, and stored at -80°C until use. Non-compartmental methods were used to obtain estimates of the terminal elimination half-life ($t_{1/2,z}$) for each animal and mean values were obtained by averaging the individual parameters.

**Bleomycin-induced pulmonary fibrosis**

12 week-old C57BL/6 male mice (Jackson Laboratories, Bar Harbor, ME) were intratracheally instilled with a single dose of 1 U/kg bleomycin sulfate in 0.9% sterile saline (50 µl/animal). Mice treated with the intraperitoneal injection were anesthetized with an intraperitoneal injection of ketamine-xylazine (100 and 10 mg/kg body weight) in 0.9% sterile saline. An IV Catheter (BD Insyte shielded IV catheter, 22 ga x 38 mm) was inserted into the trachea. Bleomycin was then administered through the catheter. Using this method, we observed limited fibrotic development (i.e., "mild" bleomycin model), and so for the inhalation study, we changed our method as advised by our IACUC committee to use isoflurane to anesthetize and added the application of approximately 10 µL of 2% lidocaine to the laryngeal area using the tip of a feeding tube (20 ga x 38mm, Instech Lab Inc.) to inhibit spontaneous closing of the vocal cords. This improved the development of fibrosis, and we refer to this as the "severe" bleomycin model. To ensure the study only included mice which developed fibrosis the following criteria were applied to all mice treated with bleomycin. Exclusion criteria A (no development of fibrosis): for inclusion in the study mice had to have a weight loss of 3% or greater by day 7 from day 0 and noticeable fibrotic regions on the micro-CT scans. Exclusion criteria B (morbid weight loss): if any mice lost 30% or more of their body weight from day 0, they were euthanized and removed from analysis. Exclusion criteria C: mice that died from unexpected complications of experimental procedures or treatments. Exclusion criteria A was in place prior to the beginning

of the study. Exclusion criteria B was instituted based on Gilhodes et al., 2017. Mouse numbers used in experiments as a result of these exclusion criteria are listed in Supplementary Table 2.

**B6_BP_dslf binder treatments**

C57BL/6 mice were intraperitoneally injected with B6_BP_dslf binder every other day starting at day 7 from bleomycin instillation and ending on day 19 for a total of 7 treatments administered. Mice were injected with B6_BP_dslf with a dose of 100 µg/kg or 1 mg/kg body weight with a volume of 10 µl/g body weight per injection. For the inhalation study, mice were anesthetized using 5% isoflurane and hung semi-recumbently by their top incisors and their tongue retracted to visualize the vocal cords. B6_BP_dslf was delivered every other day with an oropharyngeal addition of 25 to 50 µL with drug concentrations of 43.6 and 185.2 µg/kg. The B6_BP_dslf was pipetted into the oral cavity and was audibly inhaled into the lungs.

**flexiVent lung mechanics assessment**

On day 21 from bleomycin instillation, lung mechanics measurements were performed as described in Gilhodes et al and Devos et al[19,20]. These lung mechanics measurements were performed using the flexiVent FX system (SCIREQ Inc., Montreal Qx, Canada). The instrument was equipped with a FX1 module and a Negative Pressure-Driven Forced Expiration (NPFE) extension for mice run by flexiWare 8.0 software. C57BL/6 mice were anesthetized using an intraperitoneal injection of ketamine-xylazine (100 and 10 mg/kg body weight) in 0.9% sterile saline. Once mice were observed to be in a surgical plane of anesthesia, the trachea was exposed to insert a 22-gauge metal cannula. The mice were then attached to the flexiVent and received an intraperitoneal injection of 0.8 mg/kg body weight pancuronium bromide to prevent spontaneous breathing. Mice were ventilated with a tidal volume of 10 mL/kg with a frequency of 150 breaths/min and an end-expiratory pressure of 3 $cmH_2O$. The baseline was recorded, and the following scripts were run three times: Deep Inflation, Snapshot-150, Quick Prime, Negative Pressure-Driven Forced Expiration (NPFE). After the scripts were completed, mice were euthanized.

**High-resolution micro-computed tomography (HR-µCT) scans**

Mice were induced with 1.5-2.5% isoflurane and placed into the Quantum GX Micro-CT scanner (Perkin Elmer, Waltham, MA) attached to a nose cone to continue the delivery of anesthetic. Micro-CT images of the lungs were acquired using the following parameters: 90 kV, 88 uA, acquisition FOV 36 mm, Al 0.5 mm + Cu 0.06 mm filter, acquisition time 4 minutes under High-Resolution conditions. Scans were calibrated for Hounsfield units (HU) and analyzed using AccuCT™ Advanced Imaging (Perkin Elmer, Waltham, MA). A semi-automatic segmentation process was used to isolate the left and right lobes of the lung. By performing HU histogram analysis of frequency of intensities for a sample of the scan image, the threshold for lung tissue was determined. Using these thresholds, the lungs were segmented. The frequency of intensities was determined for the segmented lung. Captured lung volume, dense tissue volume, and mean intensity were also calculated. Dense tissue was defined as anything above the intensity where non-treated mice and bleomycin-treated mice intersected on the frequency of intensities histogram.

**Masson trichrome staining**

The left lobe of the lung was inflated with 4% paraformaldehyde at 25 cm $H_2O$ of pressure. Lungs were stored in PFA overnight and then washed with PBS and dehydrated using a series of ethanol washes and then processed and paraffinized overnight using the Shandon Citadel 1000 tissue processor. After the tissues were paraffinized, they were embedded into paraffin blocks and sectioned into 7 µm slices using a Microm 325 microtome. Tissue was then deparaffinized as described previously (IHC Deparaffinization Protocol, Abcam). Masson trichrome staining was performed according to manufacturer's instructions (Sigma Aldrich, HT15-1KT).

**Ashcroft scoring of masson trichrome-stained lung slices**

After Masson trichrome staining was performed, slides were scored using a modified Ashcroft scoring applied to the whole lung divided into 10x fields where sections were given a score of 0-8 per section and the total score is defined by scores summed and divided by the number of sections[21].

**Sirius Red Staining and Quantification**

**Bronchoalveolar lavage fluid (BALF) collection and cell counts**

BALF was collected by intubating the trachea with a 22-gauge cannula. 800 µL of PBS was instilled and recovered from the lungs via syringe following three repetitive plunges. BALF was centrifuged at 10,000 rpm for 10 min, after which the supernatant was collected, and the pellet resuspended in PBS. A 250 µL resuspension of the cell pellet was spun down onto slides using the Cytospin™ 3 Centrifuge for 3 minutes at 800 rpm. Slides were stained using Wright's Stain and imaged at 1000x on the Olympus BX51 Microscope[22]. A total of 200 cells were differentially counted for macrophages, lymphocytes, and polymorphonucleocytes in duplicate and cell counts were averaged[23].

**BALF cytokine array**

The supernatant of the BALF was collected as stated above and stored at -80°C prior to processing. When thawed on ice, HALT™ Protease and Phosphatase Inhibitor Cocktail 100X (48446, ThermoFisher Scientific) was added to a final concentration of 1X and then total protein concentration was determined using the Pierce™ BCA Protein Assay Kit (ThermoScientific). Each treatment group (NT, BLM, and B6_BP_dslf 185.2 µg/kg) had three supernatants pooled together at equal concentrations. The cytokine expression was measured through the Mouse Inflammation Antibody Array - Membrane (40 Targets) (ab133999, abcam) according to manufacturer's instructions. Membranes were imaged using FluorChem imaging system (Alpha Innotech, San Jose, CA) The membranes were quantified using ImageJ software as described in Schindelin et. al[24]. The resulting densitometry for the individual cytokines were then analyzed using one-way ANOVA.

**Immunofluorescence staining**

Paraffin-embedded lung sections were deparaffinized using Histo-Clear (HS-200, National Diagnostics) three times for 5 minutes each time, after which the slides were rehydrated in serial ethanol concentrations in water for 5 minutes each step (that is, 100%-100%-80%-80%-70%-70%-30%-30%-0%). Antigen retrieval was done as described previously using citraconic anhydride solution[25]. The slides were washed three times with PBS-T then blocked using 50% Mouse-to-Mouse blocking reagent (MTM500, ScyTek Laboratories) in Seablock blocking buffer (37527, ThermoFisher Scientific) for an hour. Anti-α Smooth Muscle Actin (1:200, 48938, Cell Signaling Technology) and anti-Collagen type I (1:250, PA5-95137, Invitrogen) primary antibodies were diluted in 10% Seablock PBST and incubated with the samples overnight at 4°C. The slides were then washed three times with 10% Seablock PBST. The samples were then incubated with corresponding secondary antibodies and a DAPI counterstain (14285, Cayman). The slides were then mounted using diamond antifade (P36961, Thermo Fisher Scientific) and cure at room temperature for at least 24 h prior to imaging. Confocal imaging was done using a Leica TCS SP8 Hyvolution confocal microscope with LASX software (Leica, version 3.1.1.15751). Argon laser power was consistently set to 25%. Use of the HyD detectors were preferred over the PMT detectors and gains for the detectors are set to less than 100. The images were taken at 1024x1024 pixel format. The final images were exported as TIFF files and gamma was not adjusted for any images.

**Sandwich ELISA to quantify B6_BP_dslf in lung homogenate and serum samples**

To extract B6_BP_dslf from lung tissue, pre-weighed entire lung was homogenized in 1 mL tissue protein extraction reagent (T-PER, ThermoFisher, #78510) + 1X Halt protease/phosphatase inhibitor cocktail (ThermoFisher, #78440) using Omni ceramic bead tubes (Omni cat # 19-627) and an Omni electric homogenizer for 3 rounds of 20 sec at full speed (8.0 m/s). Homogenized samples were then incubated on ice for 2 h, centrifuged at 19,000 × *g* for 20 min at 4°C, and isolated supernatants were stored at -80°C until use in the sandwich ELISA.

Recombinant human integrin αvβ6 or αvβ8 headpieces at 1.0 μg/mL in TBS with $Ca^{2+}/Mg^{2+}$ (20 mM Tris pH 8.0, 150 mM NaCl, 1 mM $Ca^{2+}Cl_2$, 1 mM $Mg^{2+}Cl_2$) were incubated overnight at 4°C on 96-well Nunc MaxiSorp plates (ThermoFisher, #442404) (100 μL/well). Plates were blocked with 200 μL/well of blocking buffer (2% [w/v] bovine serum albumin [BSA] in TBS with 0.05% Tween20 [TBST]) and incubated for 1 h at room temperature. Plates were washed 3× in TBST using a robotic plate washer (BioTek). 100 μL of lung homogenate supernatant or serum dilutions for each mouse starting at 1:100 and serially diluting 5-fold seven times using TBS with $Ca^{2+}/Mg^{2+}$ and 0.5% (w/v) BSA (binding buffer) (8 total dilutions) were added to each well and incubated for 1 h at room temperature. Each 96-well plate with biological sample dilutions also included wells with 100 μL of B6_BP_dslf standard starting at 1 nM and serially diluting 2-fold ten times using binding buffer (11 total dilutions). After washing plates 3× in TBST, plates were incubated for 30 min at room temperature with 100 μL/well of 10 nM rabbit anti-B6_BP_dslf mAb called "A1" diluted in binding buffer. After washing plates 3× in TBST, plates were incubated for 30 min at room temperature with 100 μL/well of goat anti-rabbit IgG HRP-linked mAb (Cell Signaling Technology, #7074) diluted 10,000-fold in binding buffer. Following a final 3× TBST plate wash, 100 μL of TMB (3,3′,5′,5-tetramethylbenzidine, SeraCare, #5120-0083) was added to each well and rested for 2 min. TMB was quenched with 100 μL of 1 N HCl. Absorbance at 450 nm was immediately collected for each well on an Agilent BioTek Epoch 2 plate reader. Data were plotted in Prism 9 (GraphPad Software, San Diego, CA) to determine the 50% effective concentration ($EC_{50}$) using a four-parameter logistic regression model. A logarithmic equation fit to the linear portion of the sigmoidal curve of the B6_BP_dslf standard curve, lung weights, and $EC_{50}$ values for lung homogenate supernatant and serum were used to calculate B6_BP_dslf concentration (nM) in mouse lung tissue and undiluted sera.

**Quantification of collagen, fibronectin, and p-SMAD2 in lung homogenates**

Whole lung tissue was homogenized using 300 uL RIPA buffer (10 mM Tris-Cl (pH 8.0), 1 mM EDTA, 1% Triton X-100, 0.1% sodium deoxycholate, 0.1% SDS, 140 mM NaCl, and 1 mM PMSF) and HALT™ Protease and Phosphatase Inhibitor Cocktail 100X (48446, ThermoFisher Scientific) was added to a final concentration of 1X, and ran at 60 Hz for a total of 30 seconds per manufacturer instructions(). The tissue lysate was collected, the total protein concentration determined using Pierce ™ BCA Protein Assay Kit. Protein samples were separated through 5-10% SDS-PAGE gels and transferred to 0.2 µm nitrocellulose membranes (Bio-Rad, Cat No.1620150, Hercules, CA) through electro blotting. To block, 1% BSA in PBS and Mouse-to-Mouse Blocking Reagent (MTM500, ScyTek Laboratories Inc.) were used and the following primary antibody probes were used overnight: Collagen I Polyclonal Antibody (Cat. No. PA5-95137), Fibronectin Polyclonal Antibody (Cat. No. PA5-29578, ThermoFischer Scientific), and Phospho-SMAD2 (Ser465/Ser467) (E8F3R) Rabbit mAb (Cat. No. 18338, Cell Signaling Technology). After washing the primary antibodies off, secondary antibodies IRDye® 800CW Donkey Anti-Rabbit IgG (H + L) (Cat No. 926–3221, Licor, Lincoln, NE, 1: 15,000) and Goat anti-Mouse IgG (H+L) Highly Cross-Adsorbed Secondary Antibody, Alexa Fluor 680 (Cat. No. A-21058, Invitrogen, 1: 5,000) were used. The blots were developed using the Odyssey CLx (Model No. 9140, Li-Cor, Lincoln, NE). Quantifications were done by using ImageJ. Three biological replicates were used for each treatment group.

**Collagen quantification using Sircol ™ soluble and insoluble collagen assay**

Whole lung tissue was homogenized as described above then 100 µL of homogenate was incubated with 1 mL of acetic acid-pepsin digest overnight and the rest of the Biocolor Sircol™ Soluble Collagen Assay (S1000, Ilex Life Science) and Biocolor Sircol™ Insoluble Collagen Assay (S2000, Ilex Life Science) were done following manufacturers instructions. Three biological replicates were used for each treatment group.

**Human fluorescent lung organoid triculture (hFLO) bleomycin model**

A 3D human alveolar triculture model was created as previously described.[26] Alveolar type II cell line A549, endothelial cell EAHy, and human normal lung fibroblast cell HFL1 were incorporated in a suspension culture supplemented with 300 µg mL -1 basement membrane extract (BME). These aggregates were grown until day 7 of culture and then treated with 20 µg mL -1 bleomycin (13877, Cayman Chemical Company). Media was changed after 3 days of culture with bleomycin. As a treatment, we used 10 nM B6_BP_dslf. The cultures were kept in treatment media for 4 more days with media changing every other day. Cell aggregates were collected and fixed in 4% paraformaldehyde for 15 minutes, washed with PBS-T three times, 15 minutes each time and resuspended in 15% and 30% sucrose in PBS prior to embedding in OCT compound (Sakura FineTek). 10 µm sections were then stained, imaged, and analyzed as described previously.[26]

**Statistical analysis**

All values were reported as mean ± SD. Data were analyzed by one-way ANOVA followed by Tukey's post-hoc test for multiple comparisons. Outliers were identified using Grubb's test (Q=5%). Analysis and graphs were done with GraphPad Prism 9.0 (GraphPad, San Diego, CA. USA). Results with p-value<0.05 were considered statistically significant. Analysis was done blinded to the treatment group assigned.

**Software**

*In silico* design and the analysis of the inhibitors was performed using a combination of bash, python, and Rosetta. All scripts used in this paper have been uploaded to a github account (https://github.com/aroy10/avb6-publication). Data analyses were performed with custom code in Python and IPython.
